# Neurotransmission and neuromodulation systems in the learning and memory network of *Octopus vulgaris*

**DOI:** 10.1101/2021.11.12.468341

**Authors:** Naama Stern-Mentch, Gabriela Winter, Michael Belenky, Leonid Moroz, Binyamin Hochner

## Abstract

The vertical lobe (VL) in the octopus brain plays an essential role in its sophisticated learning and memory. Early anatomical studies suggested that the VL is organized in a “fan-out fan-in” connectivity matrix comprising only three morphologically identified neuron types; input axons from the superior frontal lobe (SFL) innervating *en passant* millions of small amacrine interneurons (AMs) which converge sharply onto large VL output neurons (LNs). Recent physiological studies confirmed the feedforward excitatory connectivity: a glutamatergic synapse at the first SFL-to-AM synaptic layer and a cholinergic AM-to-LNs synapse. SFL-to-AMs synapses show a robust hippocampal-like activity-dependent long-term potentiation (LTP) of transmitter release. 5-HT, octopamine, dopamine and nitric oxide modulate short- and long-term VL synaptic plasticity. Here we present a comprehensive histolabeling study to better characterize the neural elements in the VL. We generally confirmed glutamatergic SFLs and cholinergic AMs. Intense labeling for NOS activity in the AMs neurites fitted with the NO-dependent presynaptic LTP mechanism at the SFL-to-AM synapse. New discoveries here reveal more heterogeneity of the VL neurons than previously thought. GABAergic AMs suggest a subpopulation of inhibitory interneurons in the first input layer. Clear GABA labeling in the cell bodies of LNs supported an inhibitory VL output yet the LNs co-expressed FMRFamide-like neuropeptides suggesting an additional neuromodulatory role of the VL output. Furthermore, a group of LNs was glutamatergic. A new cluster of cells organized in a “deep nucleus” showed rich catecholaminergic labeling and may play a role in intrinsic neuromodulation. *In situ* hybridization and immunolabeling allowed characterization and localization of a rich array of neuropeptides and neuromodulatores, likely involved in reward/punishment signals. This analysis of the fast transmission system, together with the newly found cellular elements helps integrate behavioral, physiological, pharmacological and connectome findings into a more comprehensive understanding of an efficient learning and memory network.

## Introduction

Cephalopods are critical reference species in neuroscience, with multiple examples of neuronal innovations and convergent evolution (Albertin et al. 2012; Striedter et al. 2014; Yoshida et al. 2015; Nesher et al. 2020; Turchetti-Maia et al. 2017). Octopuses are known for their highly flexible behavior, which relies on various forms of associative learning, including observational learning (Alves et al. 2008; Amodio and Fiorito 2013; Boal 1996; Boycott 1954; Fiorito and Scotto 1992; Mackintosh 1965; Maldonado 1963; Maldonado 1965; Moriyama and Gunji 1997; Papini and Bitterman 1991; Sutherland 1959; Wells 1978). Their behavioral flexibility also includes solving the problems of complex motor tasks through learning strategy (Fiorito et al. 1990; Gutnick et al. 2011; Gutnick et al. 2020; Richter et al. 2015; Richter et al. 2016); see review (Nesher et al. 2020) Behavioral lesion and stimulation studies have implicated the vertical lobe (VL, **Fig. 1**) as a major part of the octopus learning system (Boycott 1961; Boycott and Young 1958; Fiorito and Chichery 1995; Graindorge et al. 2006; Shomrat et al. 2008). In addition, the anatomical organization of the VL resembles other well-studied brain structures involved in learning and memory, like the insect mushroom body and the mammalian hippocampus (see (Young 1995). The development of the VL slice preparation allowed physiological investigation of the VL (Hochner et al. 2003; Hochner et al. 2006), revealing a robust activity-dependent long-term potentiation (LTP), whose expression is very similar to that in the hippocampus. This LTP is important for acquiring long-term memory, as saturation of the LTP mechanism by electrical stimulation prior to training in a passive avoidance task, impaired the transition of short-into long-term memory (Shomrat et al. 2008).

**Figure 1.**
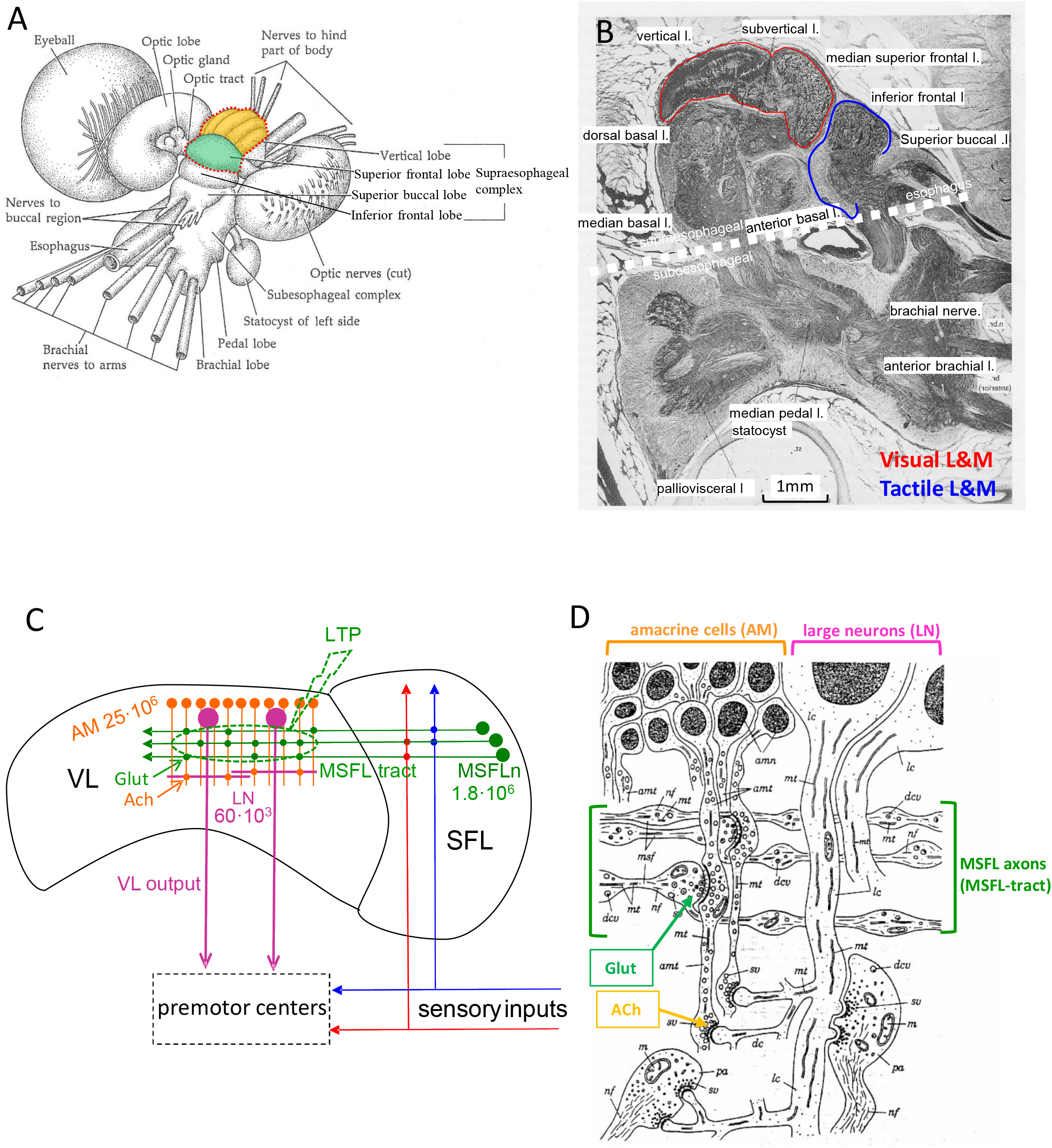
The morphological organization of the octopus brain. (A) The centralized brain of *Octopus vulgaris* in anterior-dorsal view. The supraesophageal brain complex includes the superior frontal lobes (SFL, green) and the vertical lobe (VL, orange), which consists of five cylindrical gyri. Red dotted line outlines the area of the visual learning system at the most dorsal part of the supraesophageal complex. The inferior frontal lobe (IFL), part of the touch learning system, lies ventrally and slightly anterior to the SFL (modified from Brusca and Brusca, 1990). (B) Sagittal section of the sub- and supraesophageal brain complexes showing the dorsally located median SFL (MSFL)-VL system (red line), the median IFL (MIFL)-subFL system (blue line), and the organization of the other lobes (modified from Nixon and Young, 2003). (C) A schematic wiring diagram of the neural elements of the VL system, showing the basic connectivity, types, and numbers of cells in the MSFL-VL system. Neurotransmitters and synaptic areas where long-term potentiation (LTP) occurs are marked (after Shomrat et al., 2011). (D) A classic diagram summarizing the basic circuitry of the VL (adapted from Gray, 1970) with superimposed neurotransmitters as inferred from pharmacological studies (Shomrat et al 2011). *amn*, amacrine interneurons; *amt*, amacrine trunk; *dc*, dendritic collaterals of large neurons; *dcv*, dense core vesicle; *co*, cortex of vertical lobe; *h*, hila of vertical lobe; *lc*, body or trunk of the large neurons; *m*, mitochondrion; *mt*, microtubule; *nf*, neurofilaments; *pa*, possible nociceptive axon input to the large neurons; *sv*, synaptic vesicles.

### The MSFL-VL memory system

The median superior frontal lobe (MSFL), a brain region thought to integrate sensory information (**Fig. 1**), appears to contain only one morphological type of neuron. Their axonal projections form the distinct MSFL tract into the VL neuropil (Young 1971). The tract runs in an outer neuropil layer between the deep neuropil and the external cell bodies creating the cortices of the five VL lobuli (**Fig. 1C,D**). The VL cortices contain two main classes of cell bodies, a majority of small amacrine cells (AMs; ∼3-6 μm dia.), whose neurites run radially into the center of the VL medulla, and a relatively small number of large neurons (LNs; ∼10-17 μm dia.) lying in the inner zones of the cortex, either singly or in clusters of up to six (**Fig 1C,D**). Both AMs and LNs are morphologically typical invertebrate monopolar neurons (Bullock and Horridge 1965; Young 1971).

The afferent axonal bundles from the MSFL are distributed throughout the whole VL (Young 1971), with each MSFL axon making *en passant* synapses with a yet unknown number of AM interneurons along its length (**Fig. 1C,D**). A second input to the VL is less defined but may carry pain (punishment) and reward signals. These axons enter the VL from the subVL (**Fig. 1B**) and ramify in the VL neuropil (Gray 1970). The LN trunks run radially into the medulla, where they give off dendritic collaterals, which branch profusely in the more central regions of the neuropil. Their axons project to the subvertical lobe (subVL), but little is known about other possible targets.

The general connectivity of the VL can be described as a feedforward fan-out fan-in, bi-synaptic network (Shomrat et al., 2011), whose suggested connectivity is shown in **Figure 1C,D** (Gray 1970; Young 1971, 1995). The 1.8 million MSFL neurons diverge (‘fan out’) to innervate 25 million AM interneurons (**Fig. 1C,D**; Gray, 1970; Young, 1971). The fan-in connection of the AMs onto merely 65,000 efferent LNs is mediated by serial synapses, in which the AM neurites are postsynaptic to the MSFL axon terminals and presynaptic to the spines of the LN dendrites (**Fig. 1D**; Gray, 1970). The LNs are presumed to be the only output of the VL (Young, 1971), their axons leave the VL ventrally in organized bundles or roots.

The VL bi-synaptic fan-out fan-in network, like the presumed association neural networks of the mammalian hippocampus and the insect mushroom bodies (Heisenberg 2003; Wolff and Strausfeld 2016; Young 1991, 1995), is arranged similarly to feedforward two-layered artificial “perceptron” networks (Rosenblatt 1958). Provided that one of the two synaptic layers is endowed with long-term synaptic plasticity, these produce computational functions such as association and classification (Shomrat et al. 2011; Vapnik 1998).

The MSFL-VL system is important for visual learning, while the median inferior frontal lobe (MIFL) and subfrontal lobe (subFL) form the main part of the touch learning system (Wells 1978). The morphological organization of the MIFL-subFL system resembles that of the MSFL-VL complex, but it is significantly smaller (**Fig. 1A, B**;.(Young 1971, 1991)).

Different ultrastructure of the synaptic vesicles in this system (Young, 1971) indicates the recruitment of different neurotransmitters. Pharmacological experiments were performed to elucidate putative neurotransmitters and neuromodulators of the VL system (**Fig. 1C, D**; for review (Turchetti-Maia et al. 2017). The fan-out input to the AMs appears to be glutamatergic, while the fan-in input to the LNs is cholinergic (Shomrat et al. 2011), with no direct connections from the MSFL neurons to the LNs. (Hochner et al. 2003) found no indication that the LTP induction mechanism in the *Octopus* VL is NMDA-mediated. This is not surprising since the pharmacology of NMDA receptors in mollusks often differs significantly from that in vertebrates (for example, (Moroz et al. 1993a) and a number of ionotropic glutamate receptors have been described in cephalopods (Moroz et al. 2021; Di Cosmo et al. 2006; Di Cosmo et al. 1999; Di Cosmo et al. 2004). This likely NMDA-independent LTP involves an exclusively presynaptic mechanism resembling the presynaptic expression of the non-associative LTP of mossy fiber synaptic input to the CA3 pyramidal cells of the hippocampus (Kandel et al. 2012; Yeckel et al. 1999). Yet, the LTP in the *Octopus vulgaris* VL appears to be mediated by a novel mechanism, whereby an activity-dependent constitutive elevation in nitric oxide (NO) mediates the LTP expression through NO-mediated presynaptic facilitation of transmitter release from the MSFL axon terminals (Turchetti-Maia et al. 2018).

Various neuromodulators appear to be active in the *Octopus* VL. 5-HT causes presynaptic facilitation of the glutamatergic MSF-AM synapses (Shomrat et al. 2010), possibly also indirectly enhancing the activity-dependent induction of LTP (Shomrat et al. 2010). Octopamine (OA), an excitatory neuromodulator in mollusks (Vehovszky et al. 2004; Wentzell et al. 2009) provokes a short-term facilitatory effect in the VL, like 5-HT. However, unlike 5-HT, OA attenuates LTP induction. Therefore, it was proposed that 5-HT and OA transmit punishment and reward signals into the VL where they enhance or suppress, respectively, the associative strengthening of synaptic connections (see Turchetti-Maia et al 2017).

Although specific neurotransmitters and neuromodulators have been found in the octopus brain (Messenger 1996; Shomrat et al. 2010; Shomrat et al. 2011; Taghert and Nitabach 2012; Tansey 1979; Winters et al. 2020; Shigeno and Ragsdale 2015), little is known about their precise distribution in the MSFL-VL learning and memory system. Here we characterize the anatomical distribution of candidates for neurotransmitter and neuromodulation systems in the MSFL-VL with special attention to those identified in the physiological analysis of VL connectivity, plasticity and neuromodulation (see (Shomrat et al. 2015; Turchetti-Maia et al. 2017). Our anatomical findings advance the understanding of the functional organization of the ‘fan-out fan in’ network. We show here that this network is more complex than the previously reported two simple homogenous neuron layers. Lastly, we characterize the distribution of possible neuromodulators and identify and generally localize candidate neuropeptides involved in learning.

## Materials and Methods

Adult *Octopus vulgaris* weighing ∼150-450 grams were captured off the coast of Israel. They were kept individually in closed synthetic seawater aquaria (80-100 liters) with biological and chemical filters and maintained at 18°C on a 12h light/dark cycle (Hebrew University) or in semi-open seawater systems, both light- and temperature-controlled (Ruppin Faculty of Marine Sciences, Michmoret, Israel). Animals were acclimatized to the laboratory for at least two weeks before experiments, conforming to the ethical principles of EU Directive 2010/63/EU, the principle of the 3Rs (Replacement, Reduction and Refinement), and minimization of suffering (see (Fiorito et al. 2014; Fiorito et al. 2015).

Animals were deeply anesthetized in fresh seawater supplemented with 2% ethanol and 55 mM MgCl_2_ (Shomrat et al. 2008). The brain was removed through a dorsal opening of the cartilaginous brain capsule. For most histochemical procedures, the brain was immediately fixed by immersion 4hr-overnight in 4% PFA in artificial sea water (ASW) (Shomrat et al. 2008) or 0.1 M phosphate-buffered saline (PBS), pH 7.4 at 4 °C. **Table 1** gives the fixation and preservation solutions used for each primary antibody. Brain slices, in vivo and in vitro, were obtained as described in Shomrat et al. (2008). After washing with ASW or PBS, the fixed supraesophageal mass was glued with acrylic glue to a vibratome stage (Leica, VT1000 S). 50-90 µm sagittal, transverse or horizontal sections were used for immunohistochemistry (Shomrat et al. 2010)

**Table 1:**
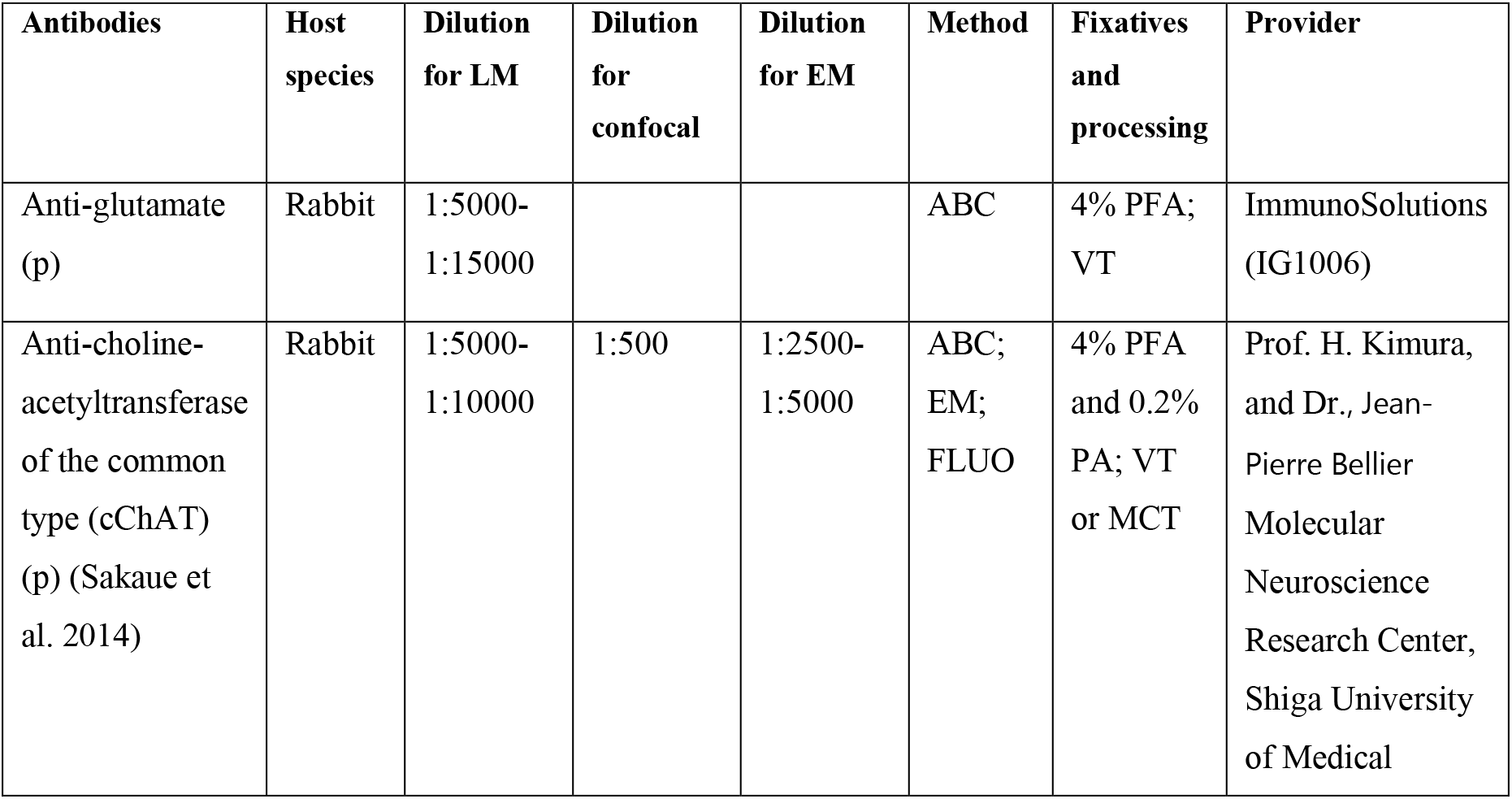
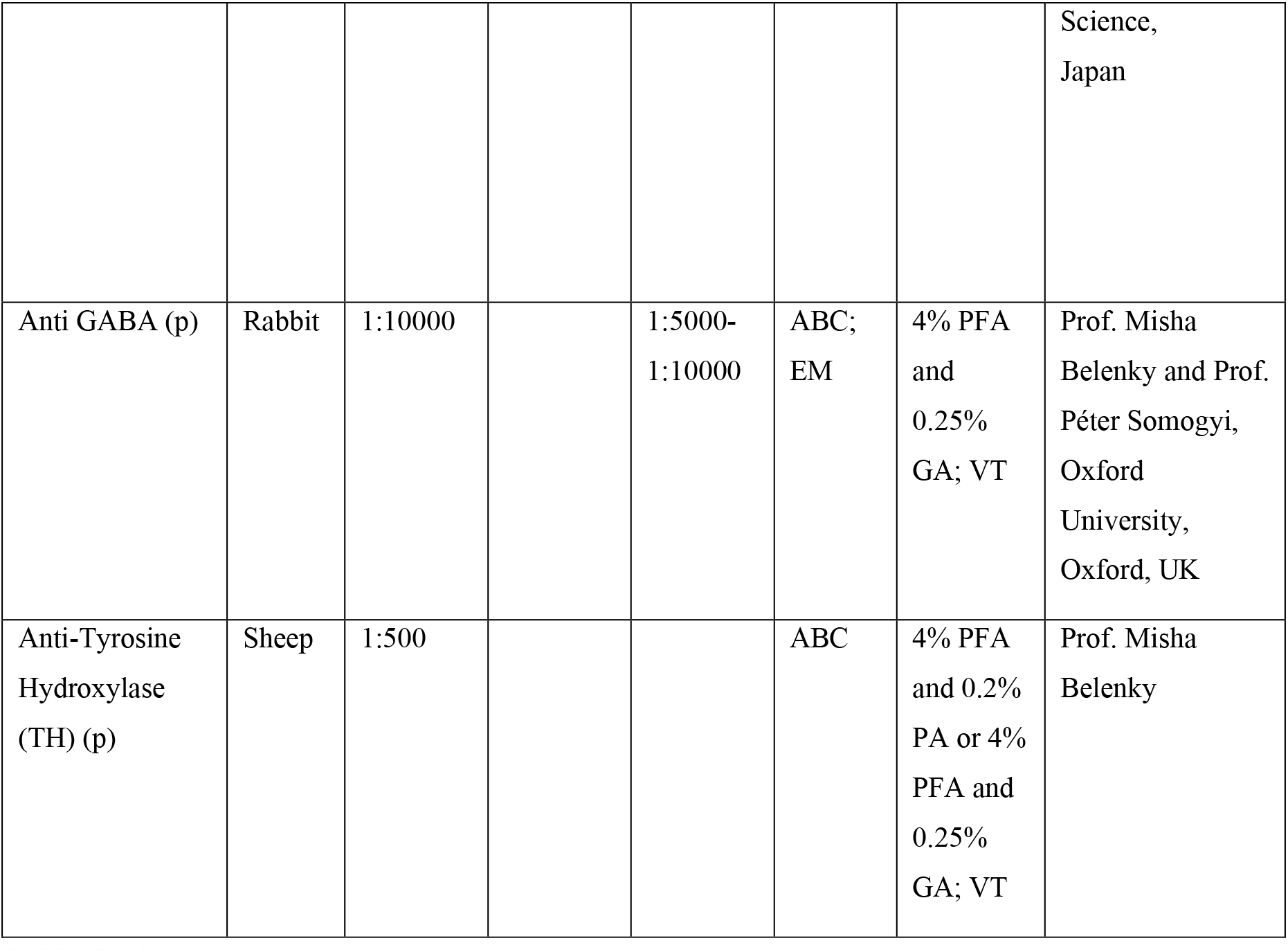
Antibodies, concentrations, information on methods, fixation, sample processing and provider. *p* polyclonal *ABC* avidin-biotin complex, *GA* glutaraldehyde, PFA paraformaldehyde, PA picric acid, EM electron microscope, FLUO immunofluorescence method, VT vibratome, MCT microtome

For cChAT confocal microscopy experiments, after fixation and washing, tissues were immersed for at least 24 hours in PBS containing 30% sucrose at 4°C, frozen on the microtome base with dry ice, and sectioned sagittally on a sliding microtome (40 µm). Free-floating sections were collected in ice-cold PBS. Each type of marker was tested on a minimum of two brains, eight slices per brain.

### Immunohistochemistry

An immunoperoxidase procedure was performed on free-floating sections for the immunohistochemical detection of target epitopes in EM and LM sections, similar to the protocol in Shomrat et al. (2010; differences are noted in **Table 1**).

#### Immunoperoxidase labelling

After slices were incubated in primary antibody (**Table 1**) and washed, they were incubated for 3 h, RT in biotinylated secondary antibody (1:600 goat anti-rabbit, Vector Laboratories, USA). The sections were placed in avidin-biotin peroxidase complex (ABC Elite, Vector Laboratories, USA) for 1h at RT. PBS was used after each step. Peroxidase activity was visualized as a brown precipitate by reacting the sections for 3-7 minutes at RT with a solution containing 0.04% 3-3diaminobenzidine-tetrachloride (DAB) and 0.006% H_2_O_2_. When picric acid was used for pre-staining tissue preservation, the DAB solution also included 0.4% nickel ammonium sulfate in 50 mM Tris-HCl buffer (pH 7.6) to yield a dark blue precipitate. The reaction was stopped by rinsing with PBS or Tris-HCl buffer, accordingly. For EM, the reaction product was intensified and substituted with silver/gold particles, as described by (Livneh et al. 2009). For control experiments, sections were processed as described above but without the primary antibody, resulting in no specific staining.

LM sections were mounted on SuperFrost Plus slides (Menzel Glaser, Germany) and air-dried. The sections were then dehydrated in a graded ethanol series (70–100%), cleared with xylene, and cover-slipped with Enthellan for observation under an Olympus BX43 microscope.

#### Immunofluorescence

Slices were incubated in rabbit anti-cChAT primary antibody (**Table 1**) and washed (omitting incubation in H_2_O_2_). Samples were then incubated for 2 h with anti-rabbit Alexa Fluor 594 secondary antibody (1:500; Thermo Fisher Scientific). Slices were washed and mounted on microscope slides. Fluorescent counterstaining of cell nuclei was carried out in a PBS solution with 0.1 µg/ml 4’,6-diamidino-2-phenylindole (DAPI; Roche Molecular Biochemicals, Indianapolis, IN, USA). Fluorescence was detected, analyzed and photographed with an Olympus BX43 microscope or with an Olympus FV-1200 confocal microscope. Alexa Fluor 594 was excited with the 561 nm laser and the emission wavelength was 570-620 nm. Images were prepared using NIH ImageJ software (Bethesda, MD, USA).

For EM, the sections were post-fixed, treated and observed according to Shomrat et al. (2010). Photomicrographs were unaltered except for brightness/contrast enhancement.

#### NADPH-diaphorase staining

The brain tissue was fixed by immersion in 4% PFA in ASW pH 7.4 at 4 °C overnight. In later experiments 0.25% glutaldehyde (GA) was added to the fixation solution. After slicing (50-100 µm), sections were analyzed for NADPH-diaphorase activity according to (Hope and Vincent 1989) and modifications by Moroz (Moroz 2000; Moroz et al. 2000). At the final stages, slices were dehydrated in ethanol, cleared in xylene, mounted on SuperFrost Plus slides (Menzel Glaser, Germany) and viewed with Olympus SZX16 stereo and Olympus BX43 microscopes. Specificity of NADPH-diaphorase staining was tested in control experiments in which tissue sections were incubated in the reaction solution as described, except that β-NADPH or NBT were omitted. All chemicals for this procedure were purchased from Sigma–Aldrich unless otherwise indicated.

### In situ hybridization

All procedures were carried out at RT unless otherwise stated. Samples were agitated only during antibody incubation and washes. Particular attention was paid to maintaining an RNase-free environment. To help avoid contamination, solutions prepared according to Jezzini et al. (2005) were made in small batches in disposable sterile 50 mL plastic centrifuge tubes (Corning Incorporated, NY, USA; Cat. No. 430921). The specimens were incubated in disposable sterile 24-well cell culture plates (Corning Incorporated, NY, USA; Cat. No. 3524).

#### In situ hybridization probe preparation

The molecular information for target mRNAs and sequences is summarized in **Supplementary Data.** Digoxigenin-labeled antisense RNA probes were transcribed *in vitro* using SP6 or T7 polymerases (according to the insert’s specific orientation into the plasmid) from full-length cDNA clones, ligated into p-GEM T vector (Promega) and linearized with the appropriate restriction endonucleases. Full-length sense probes were used for negative controls. One µl of the restriction digest ran on a 1% agarose gel with ethidium bromide staining to check for quality of the reaction, and the remainder was purified using a PCR purification kit (Qiagen). Typically, 13 µL of the purified linearized plasmid (approx. 1µg plasmid) was used as a template in the probe synthesis reaction. This was carried out using the DIG RNA Labeling Kit (SP6/T7) (Roche; Cat. No. 1175025) according to the manufacturer’s instructions (13µL template, 2µL NTP labeling mix, 2 µL 10× transcription buffer, 1µl RNase inhibitor, 2µL SP6 or T7 RNA polymerase, 37 ◦C for 2 h). The reactions were stopped by adding 2µL 0.2 M EDTA, pH 8.0. The NTP labeling mix contained either DIG-11-UTPs (DIG RNA Labeling Mix, Roche; Cat. No. 1277073) for synthesis of DIG-labeled probes or Fluorescein RNA Labeling Mix (Roche; Cat. No. 1685619) for synthesis of fluorescein-labeled probes. The quality of plasmids, restriction digests, and synthesized probes was checked on a 1% agarose gel with ethidium bromide staining prior to use. Probes up to 1 kb or longer were used at full length (see CNS preparation and details in Supplementary Data).

#### CNS processing and probe hybridization

In situ hybridization (ISH) experiments were performed as described previously (Jezzini et al. 2005; Winters et al. 2020; Moroz and Kohn 2010) with minor modifications. Immediately following removal from anesthetized animals, the supraesophageal mass was fixed in 4% formaldehyde in PBS, pH 7.4 overnight at 4 °C. The brain mass was then washed in three 15 min changes of PBS before being placed in a tub with fresh PBS and sectioned at 300-450 µm. All slices were washed three times with PBS, transferred to PTW (0.1% Tween 20 in PBS) for 10 min and subsequently dehydrated in sequential 10 min incubations in methanol (30-70%) in PTW, followed by 100% methanol at −80 ◦C overnight. The slices were then shipped from the Hebrew University to the University of Florida, St. Augustine, on dry ice. They were rehydrated by sequential 10 min incubations in methanol (70-30%) in PTW followed by 100% PTW for 15 min. They were then washed with 0.3% Triton X-100 in PBS for 10 min, PBS for 10 min, and PTW for 5 min. Proteinase K (Roche Diagnostics) was added to the PTW to a final concentration of 10 µg Proteinase K per 1 mL PTW (typically around 0.6 µl Proteinase K per 1 ml PTW). The slices were incubated at RT for 20 min (or until they started to appear slightly translucent around the edges). Proteinase K activity was terminated by transferring slices to 4% formaldehyde in PBS for 20 min at 4°C. Following post-fixation in 4% formaldehyde, the slices were washed in two changes of PTW followed by two changes of PTW and three changes of TEA HCl (0.1 M triethanolamine hydrochloride adjusted to pH 8.0 with NaOH). With the slices in a 1 mL volume of 0.1 M triethanolamine hydrochloride adjusted to pH 8.0 using sodium hydroxide, 2.5µL of acetic anhydride was added slowly to the TEA HCL while stirring. The slices were left for 5 min before adding an additional 2.5µL acetic anhydride while stirring, followed by another 5 min incubation. Next, the slices were washed in three changes of PTW before being transferred to hybridization buffer (HB: 50% formamide, 5 mM EDTA, 5× SSC (20× SSC: 3 M NaCl, 0.3 M sodium citrate, pH 7.0), 1× Denhardt solution (0.02% ficoll, 0.02% polyvinylpyrrolidone, 0.02% bovine serum albumin), 0.1% Tween 20, 0.5 mg/mL yeast tRNA (GIBCO BRL)). The slices were then left overnight in HB at −20◦ C before prehybridization incubation for 6–8 h at 50 °C the next day. Next, 2–6 µL (∼1µg/mL) of each probe was added and hybridization allowed to proceed for 12–14 h at 50 °C.

#### Immunological detection

Immunological detection was performed using antidigoxygenin-AP Fab antibody fragments at a dilution of 1:2000 (Roche Diagnostics, Mannheim, Germany). After probe hybridization, the slices were washed in 50% formamide/5× SSC/1% SDS (sodium dodecyl sulfate, Fisher 20% solution BP1311) for 30 min, then 50% formamide/2× SSC/1% SDS for 30 min at 60 °C, and two 30 min changes of 0.2× SSC at 55° C. Slices were transferred to PBT (0.1% Triton-X 100, 2 mg/ml bovine serum albumin, in PBS; pH 7.4) for 20 min followed by two 20 min changes of PBT at RT. Goat serum was added after the third change to a concentration of 10% by volume and the slices were then incubated for 90 min at 4° C with gentle shaking, after which the sections were placed in 1% goat serum in PBT. Alkaline phosphatase-conjugated antibodies were then added and incubation proceeded for 12–14 h at 4° C with gentle shaking.

Development using the NBT/BCIP alkaline phosphatase substrate - single probe labeling: After incubation with antibody, the slices were transferred to PBT at 4° C and washed in three 30 min changes of PBT at 4°C followed by three 5 min changes of filtered NBT/BCIP detection buffer (NDB: 100 mM NaCl, 50 mM MgCl_2_, 0.1% Tween 20, 1 mM levamisol, 100 mM Tris–HCl; pH 9.5) at 4° C. 20 µl/ml of NBT/BCIP stock solution (NBT/BCIP: 18.75 mg/ml nitro blue tetrazolium chloride, 9.4 mg/ml 5-bromo-4-chloro-3-indolyl phosphate toluidine salt in 67% dimethyl sulfoxide, Roche; Cat. No. 1681451) was added to the third change of NDB while stirring thoroughly until completely dissolved. The slices were kept on ice in the dark, and every 10 min were monitored briefly for staining to avoid excessive exposure to light. Development was terminated after cell-specific labeling was clearly visible and before excessive background began to appear. Development was stopped by transferring the slices to 4% formaldehyde in methanol for 60 min at 4° C followed by a final transfer into 100% ethanol at 4° C, immediately followed by washing in two 10 min changes of 100% ethanol at 4° C. The slices were cleared in methylsalicylate (for about 1 min or until they sank to the bottom) and mounted on microscope slides in Permount (Fisher).

Images were acquired digitally with an Olympus DP-73 camera mounted on Olympus SZX16 stereo and Olympus BX43 binocular microscopes. The diameters of immunoreactive cells were measured on photomicrographs of sagittal and transverse sections with LITE (Leica) or FIJI (ImageJ) software.

## Results

### Canonical ‘fast’ transmitters

#### Glutamate

*In situ* hybridization (ISH) revealed that most of the cell bodies in the MSFL cortex and cortices of several other lobes expressed vesicular glutamate transporter VGLUT-encoding transcripts ( **Fig. 2A**). In contrast, the VL cortex was barely labeled. Light traces of VGLUT transcript expression appeared in the MSFL axonal projections in the outer ventral neuropil (**Fig. 2B, lower bracket**), suggesting that respective mRNAs could be transported to distant neurites as in *Aplysia* (Puthanveettil et al. 2013).

**Figure 2.**
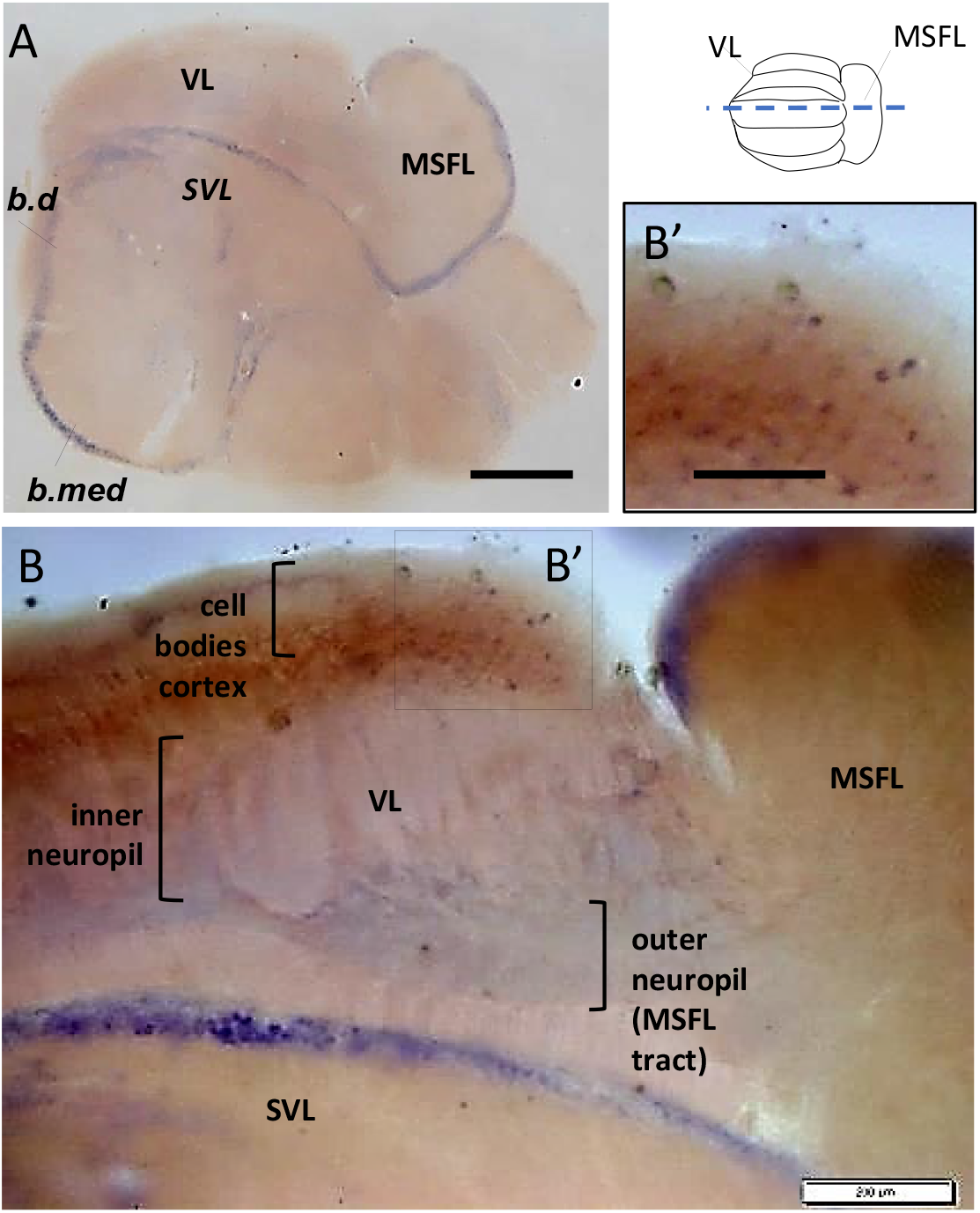
Glutamate-transporter mRNA expression highlights the MSFL cell body cortex. Inset, schematic dorsal view of the *Octopus* MSFL-VL lobes. Dashed blue line represents the plane of sections in figure B (and all figures thereafter). (A) *In situ* hybridization (ISH) on a sagittal slice reveals expression of vesicular glutamate transporter (VGLUT) mRNA in MSFL cell cortex that contrasts greatly with the generally unlabeled cells of the VL cortex. Expression of VGLUT can also be seen in the cell layers of the SVL, basal dorsal (b.d), and basal medial (b.med) lobes. (B) Photomicrograph of *in situ* hybridized sagittal section showing VGLUT mRNA expressions in the ventral outer VL neuropil (MSFL-tract, lower bracket) in large cells in anterior regions (marked region). (B’) Magnification of VGLUT expressing cells of the dashed square region marked in B. Scale bar: 500 µm (A), 200µm (B), 100 µm (B’)

**Fig. 4A** shows similar results with immunohistochemistry (IHC) labeling for L-Glut; the cell bodies in the MSFL cortex showed stronger glutamate immunoreactivity (L-glu-IR) than the generally pale L-Glut labeling in the neighboring AM cell bodies of the VL cortex. Thus, some MSFL neurons are indeed potential sources for glutamatergic input to the VL (Hochner et al. 2003; Shomrat et al. 2011). In addition, L-Glut antibody revealed clusters of L-Glut-IR LN (18-30 µm dia.) with positively labeled neurites detected in several individual cells. This group of cells was organized separately from the cell body cortex as a ‘deep nucleus’ located at the MSFL -VL border and likely interacts with the main MSFL-VL circuitry (**Fig. 3**).

**Figure 3.**
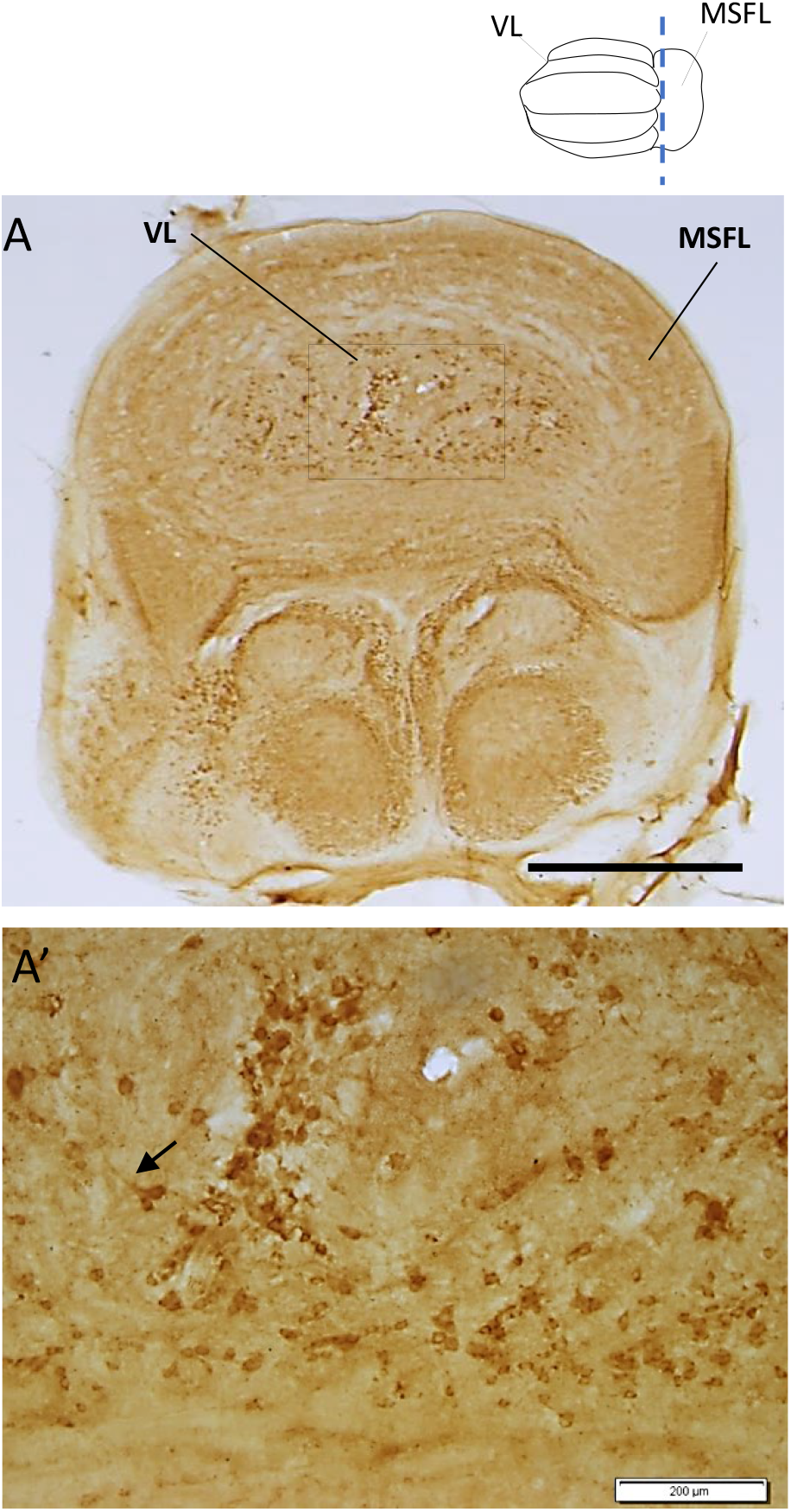
Glutamate immunohistochemistry reveals a ‘deep nucleus’ in the VL-MSFL system. (A) The transverse section shows the L-Glu positive large cell bodies (18-30µm in diameter) in the area between the MSFL and VL. They are distinctly organized as a ‘deep nucleus’. (A’) Higher magnification of the area marked in (A) showing large L-Glu-IR cell bodies. Positive neuronal processes projecting from these cells can also be seen (arrow). Scale bar: 1mm (A), 200µm (A’).

**Figure 4.**
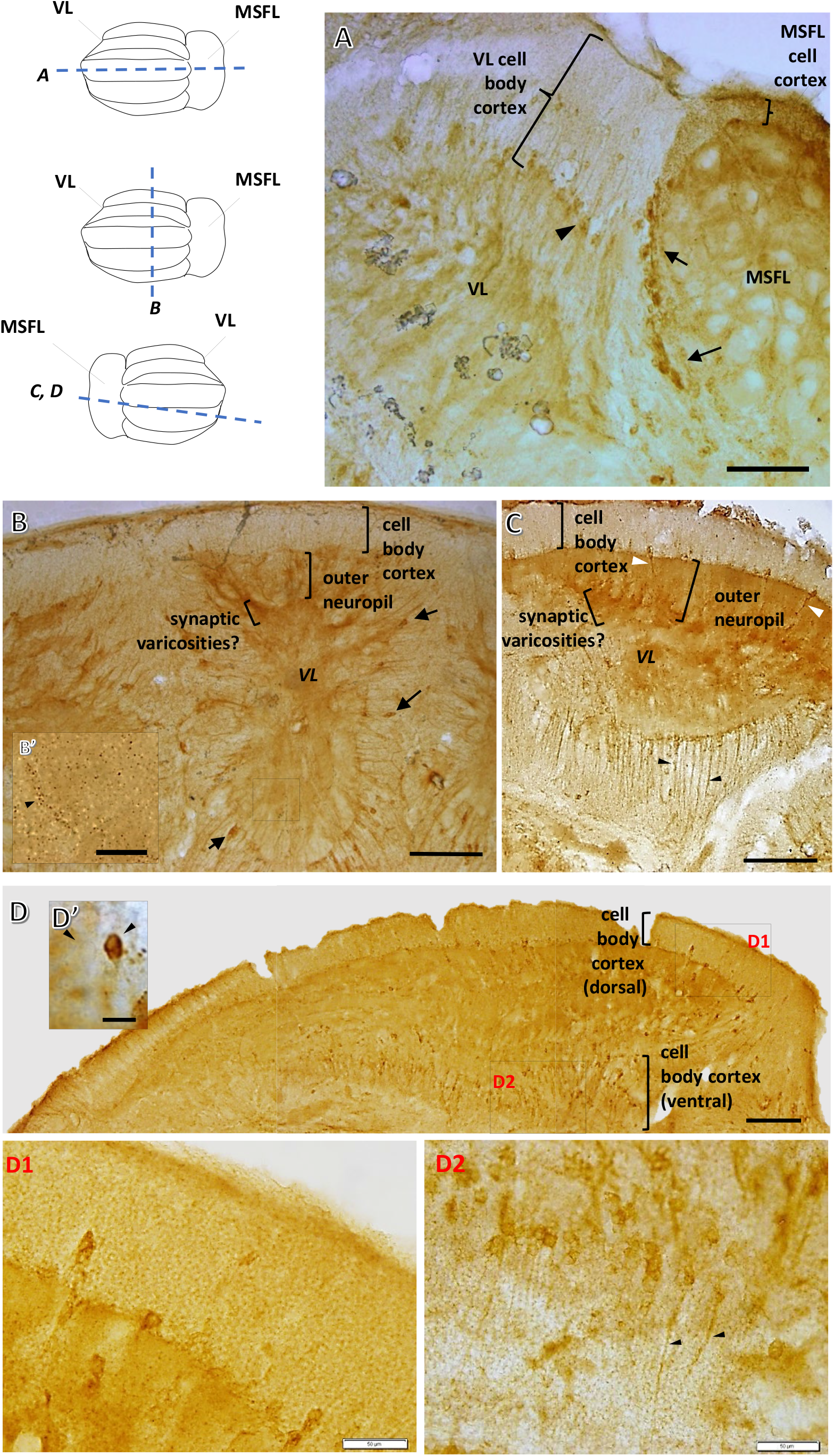
Distribution of glutamate-positive cell bodies and neuronal processes in the VL. (A) MSFL cell body cortex reveals stronger glutamate immunoreactivity (L-Glu-IR) than the global pale L-Glu labeling in the neighboring cell body cortex of the VL. L-Glu-IR also reveals a group of large cells at the border between the MSFL and VL (arrows). Arrowhead marks large cells positively labeled in the inner margin of the VL cell body cortex. (B) Photomicrograph of a transverse section of the VL showing L-Glu-IR in relatively large cells (6-13µm dia.). Intensified markings are localized at the border area between the outer and inner neuropil, suggesting this area as a site of greater synaptic densities (lower bracket). Large cell bodies are labeled (arrows). (B’) Magnifications of the area marked in B showing granular labeling in the VL neuropil. Intensified granular L-Glu labeling in the border zone between the outer and inner neuropil (circling the center of the medulla; arrowhead), again suggesting a unique compartmentalized area with high synaptic densities. (C) Light micrograph of the sagittal section showing darkened markings in the border area of the outer and inner neuropil similar to those seen in transverse section in B. L-Glu-IR large cell bodies are localized in the dorsal region, their processes projecting into the neuropil (white arrowheads), while glutamatergic processes from large cells in the ventral region mainly project inwards to the ventral cell body cortex or to the subVL (black arrowheads). (D) Photomicrograph of the sagittal section showing glut-IR of relatively large cells distributed in the inner layers of the dorsal and ventral cell cortices surrounding the VL lobuli. The L-Glu positively labeled cells in the dorsal cortical cell layer are organized in a typical row along the inner margin of the cell cortex, resembling the appearance and organization of the large neurons (LNs). The L-Glu-IR cells in the ventral region are more crowded and uniquely organized, differing from ‘typical’ LNs (lower bracket). (D’) Enlarged example of glutamate-IR cell (∼ 9µm; arrowhead, right) in the VL, taken from section similar to that in D, neighboring a cell similar in size but lacking glutamate-IR (arrowhead, left). (D1, D2) Magnification of L-Glu-IR large cells in the dorsal and ventral cortex layers seen in dashed squares marked in D. Scale bar: 200µm (A-D), 50µm (B’, D1,D2), 10µm (D’).

The VL showed intense granular L-Glut-IR in the area of the MSFL axonal projections (**Fig. 4B, C**). This particularly dark labeling at the lower region of the outer neuropil was localized at the area of the *en passant* synaptic connections right at the junction of the AM trunk and the MSFL tract (Gray 1970; Shomrat et al. 2011), probably marking the synaptic varicosities rich in glutamatergic vesicles.

L-Glu IR was also found in medium to large LN cells (6-12.5 µm dia.) in the inner half of the VL cell cortex (**Fig. 4**), either lying alone or as a small group among unstained cell bodies. Remarkably, these positively stained cells were located in the dorsal cell layer in classic locations of the LNs. Indeed, large VGLUT transcript-expressing cell bodies were revealed in similar cortical areas (**Fig 2B**). Thus, some of the LNs in the VL appear to be glutamatergic.

The L-Glu-IR LNs in the ventral cell cortex appeared more dense and visible neuronal processes projected mainly inward towards the cell layers and not into the neuropil (**Fig. 4C, D, D2**, arrowheads). These ventral glutamate-IR processes hint that some neurites do not necessarily exit with the common LN roots crossing the VL hila but, instead, may maneuver their way out of the VL through the small-cell body layer to the subVL ventral to the VL. These ventral LNs may thus represent a separate population or sub-type of LNs.

#### Cholinergic system

We used cChAT antibody, kindly provided by Prof Kimura and Dr Bellier to identify cholinergic neurons (see (Casini et al. 2012; D’Este et al. 2008; Sakaue et al. 2014). The semi-sagittal section in **Fig. 5** shows cChAT immunoreactivity in the MSFL-VL system compared with an unstained control section. While most positively labeled supraesophageal lobes did not exhibit a clear difference in the density of labeling between the neuropil and the cell body cortices, the VL is remarkable in the dense labeling of its inner neuropil, quite distinct from the surrounding cell bodies. The global cChAT-IR of the MSFL was much weaker than the richly stained VL neuropil mass (**Fig. 5A**).

**Figure 5.**
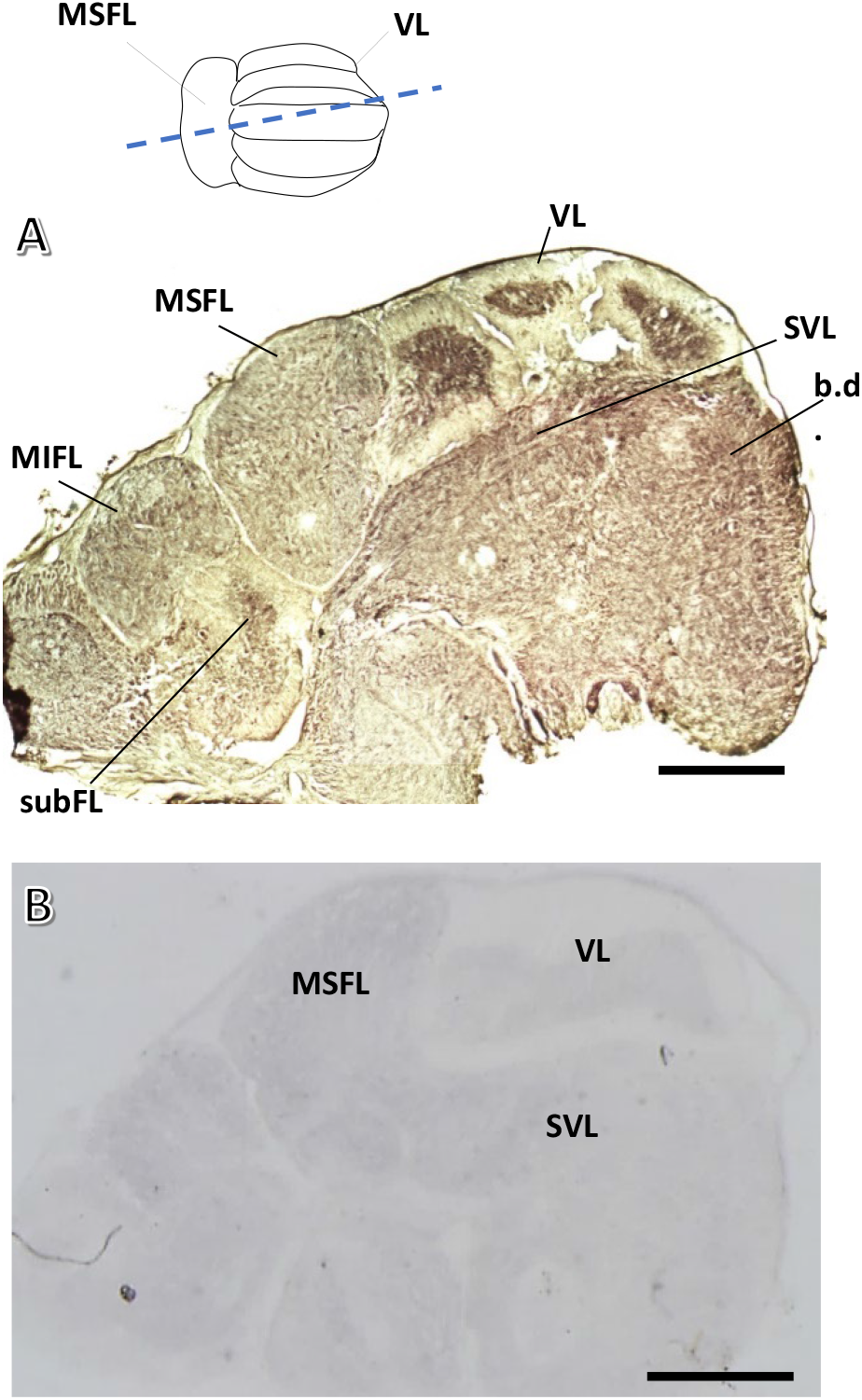
Light micrograph showing immunohistochemical staining for cChAT in the supraesophageal lobes of the octopus brain. (A) Low magnification of a stained semi-sagittal section of the supraesophageal brain mass (more than one lobulus is seen) reveals a wide distribution of cChAT immunoreactivity throughout many of the lobes, with a uniquely dense labeling of the VL and subFL inner neuropil (note that only small part of the subFL inner neuropil is included in this section). There is a similar but less intense pattern of labeling in the MSFL and MIFL. Unlike the VL there is no clear difference between the densities of the cell body cortices and the neuropilar structures. (B) Light micrograph of control sagittal section with secondary antibody only. Scale bar: 1mm

**Fig. 6A, A’** show scattered fibers labeled in the MSFL neuropil with a slightly increased density in the outer neuropil below the cell body cortex. Similar results were obtained using fluorescent labeling (**Fig. 6B**). Thick cChAT-IR neuronal processes were visualized mainly in the posterior area of the outer MSFL neuropil. At the same time, scattered fibers were labeled in other parts of the MSFL neuropil, indicating cholinergic innervation of the MSFL outer neuropil.

**Figure 6.**
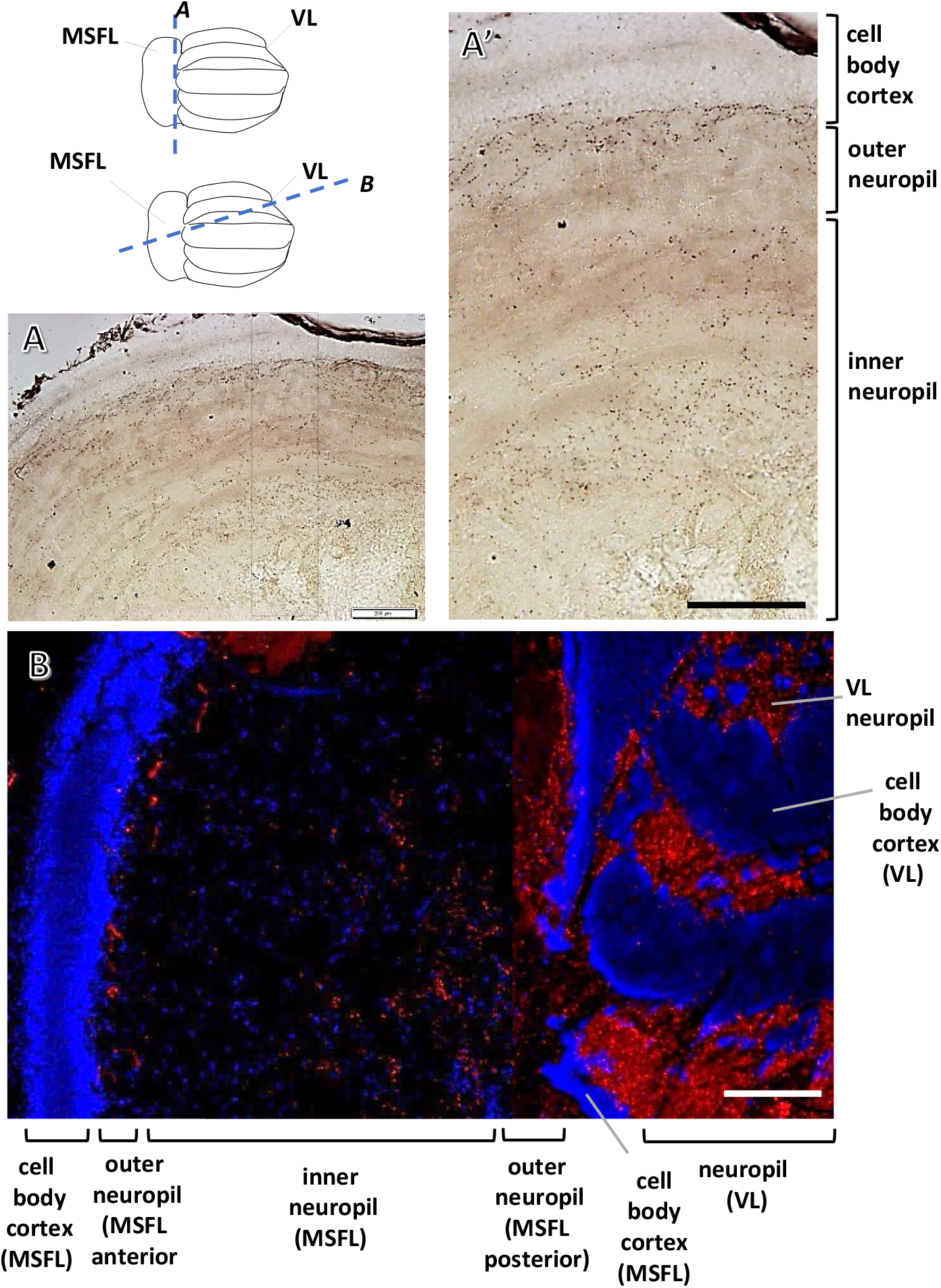
cChAT immunolabeling in the MSFL. (A) Light micrograph of a transverse slice sampled from a posterior region of the MSFL showing from where A’ was taken. (A’) Some fibrous cChAT-IR labeling can be detected in the MSFL neuropil, where labeling was slightly greater in the outer neuropil and in the center of the medulla where axons from other areas converge. (B) Fluorescent light micrograph of a sagittal-horizontal section showing VL lobuli at the right of the image, revealing similar MSFL cChAT labeling as in A’. The fluorescent labeling is scarce in the middle regions of the MSFL neuropil and slightly more intense in the outer neuropil (anterior and posterior) near the MSFL cell body cortex. Note the abundantly stained VL neuropil (red, cChAT; blue, DAPI). Scale bar: 200µm

The occasional cChAT granular markings detected along the MSFL tract axons in the outer neuropil are likely cChAT-IR AM trunks crossing through the unlabeled axons of the tract in the outer neuropil (**Fig. 7A,A’,C,C’**, arrows). Fluorescence microscopy revealed dense labeling in the outer VL neuropil, again seemingly belonging to AM trunks running vertically over the unlabeled MSFL axon into the deep neuropil (**Fig. 7A**). Associations of fluorescent labeling with presumed AMs trucks were seen by merging DIC and fluorescent images captured with a confocal microscope (**Fig. 7A’** arrow).

**Figure 7.**
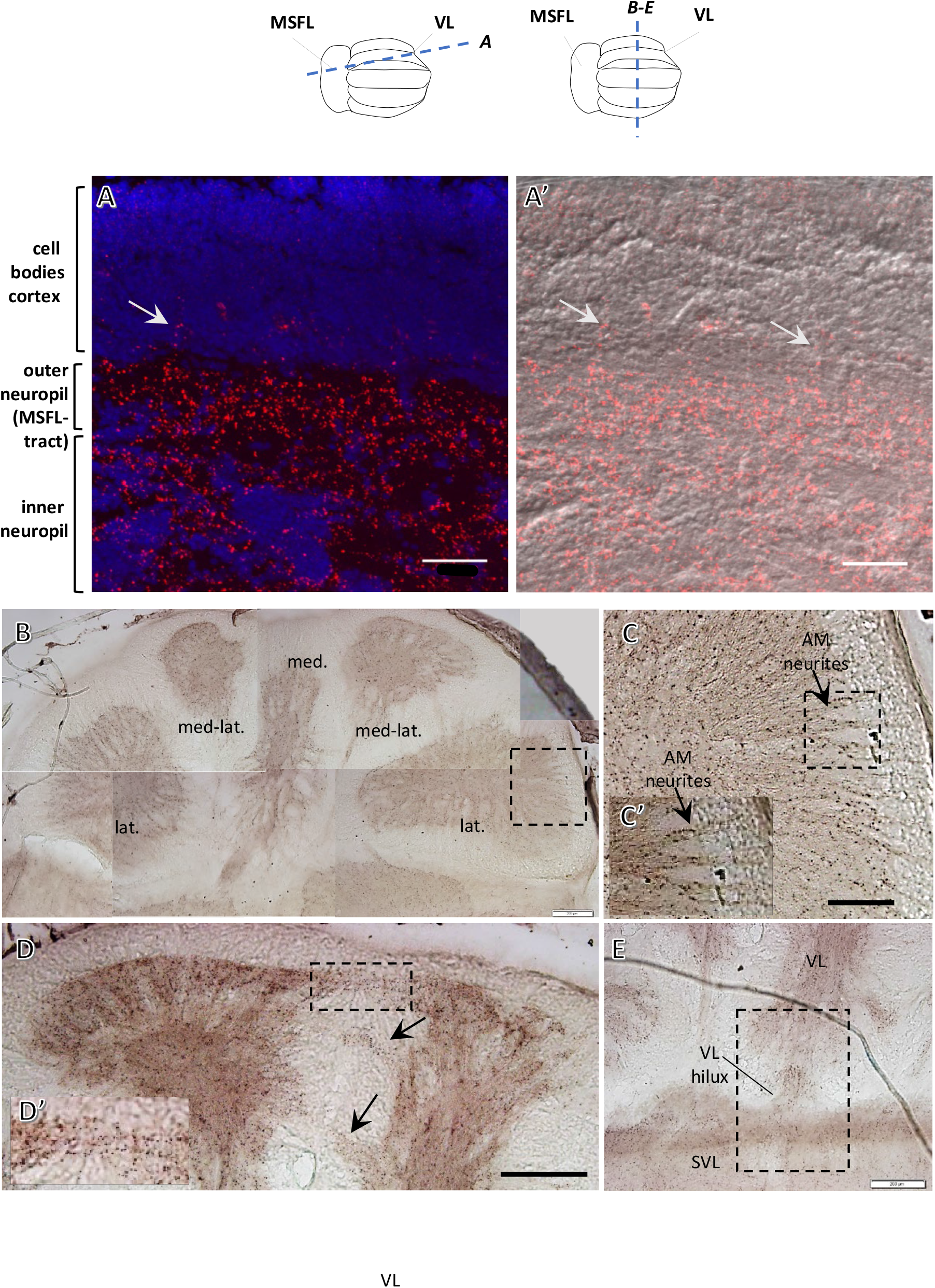
Distribution of strong cChAT positive labeling in neuronal processes in the VL. (A, A’) Confocal micrograph showing that the fluorescent signal of the processes in the VL neuropil can be followed vertically into the cell body cortex (arrows). Scattered punctate labeling in the outer cell layers is not of distinct cell bodies counterstained with DAPI (red, cChAT; blue, DAPI). (1:500). (A’) Differential interference contrast (DIC) of the cell body cortex and VL neuropil merged with the fluorescent signal (excluding DAPI) of the image in A. (B) Transverse slice of the VL shows the cChAT neuropil labeling is distributed evenly throughout the five lobuli. Marked region indicates area enlarged in C (1:15000). (C) Abundant cChAT immunoreactivity of processes most likely belonging to bundled AM neurites running vertically (example of three labeled neurite bundles are shown in inset C’). These bundles cross the apparently unlabeled MSFL tract with gaps of ∼25-30µm between them to pass into the neuropil. (D) cChAT-IR along inter-lobule connections between medial and medial-lateral lobuli in a transverse section (marked region and others marked with arrows), suggesting a cholinergic feed among the different lobuli. (1:15000). (D’) Enlargement of the marked area of crossing fibers. (E) Fibrous cChAT-IR crossing the VL hilux between the VL and the SVL in a transverse section (1:15000). 50µm (A, A’); 200 µm (B, E,); 100 µm (C).

The intense cChAT-positive fiber labeling was similarly distributed in the neuropil across all five VL lobuli (**Fig. 7C**). The cChAT-IR groups of bundled neurites running from the cell cortex crossed the unlabeled MSFL-tract fibers with gaps of ∼25-30 µm between them and projected into the homogenous densely labeled inner neuropil. These results further support physiological findings that at least some AMs are cholinergic (Shomrat et al. 2011).

cChAT-IR labeling was also observed along inter-lobulus connections running through the cell body cortex (**Fig. 7D,D’**), suggesting cholinergic processes crossing between the VL lobuli. Note that Young (1971) considered the inter-vertical tracts crossing between the lobuli to be separate packets of MSFL tract fibers rather than AM projections.

In addition, cChAT-IR labeling (**Fig. 7E**) indicates that cholinergic fibers may run between the VL and the SVL through the VL hilum.

Electron microscopy (EM) showed a field in the inner medullary zone packed with extremities of transversely cut AM trunks with their characteristic clear agranular vesicles (Gray, 1970), which exhibited cChAT-IR (**Fig. 8A**). Some of these amacrine processes showed synaptic connections with pale structures presumed to belong to the LN dendritic branches (**Fig. 8A**; double membrane, arrowhead). In addition, strong cChAT-IR was revealed in neuronal structures and processes (**Fig. 8B**). Some structures contained granulated vesicles varying in size up to 100 nm (**Fig. 8C**). These vesicles did not appear to belong to the MSFL neurons, nor did they fit the classifications of amacrine vesicles or vesicles of ‘pain’ fibers ascending into the lobules from below (Gray, 1970). Similar to the LM immunolabeling, visualization of the VL cell layers with EM barely revealed positive cChAT reactivity in the cytoplasm of the amacrine cell bodies (not shown).

**Figure 8.**
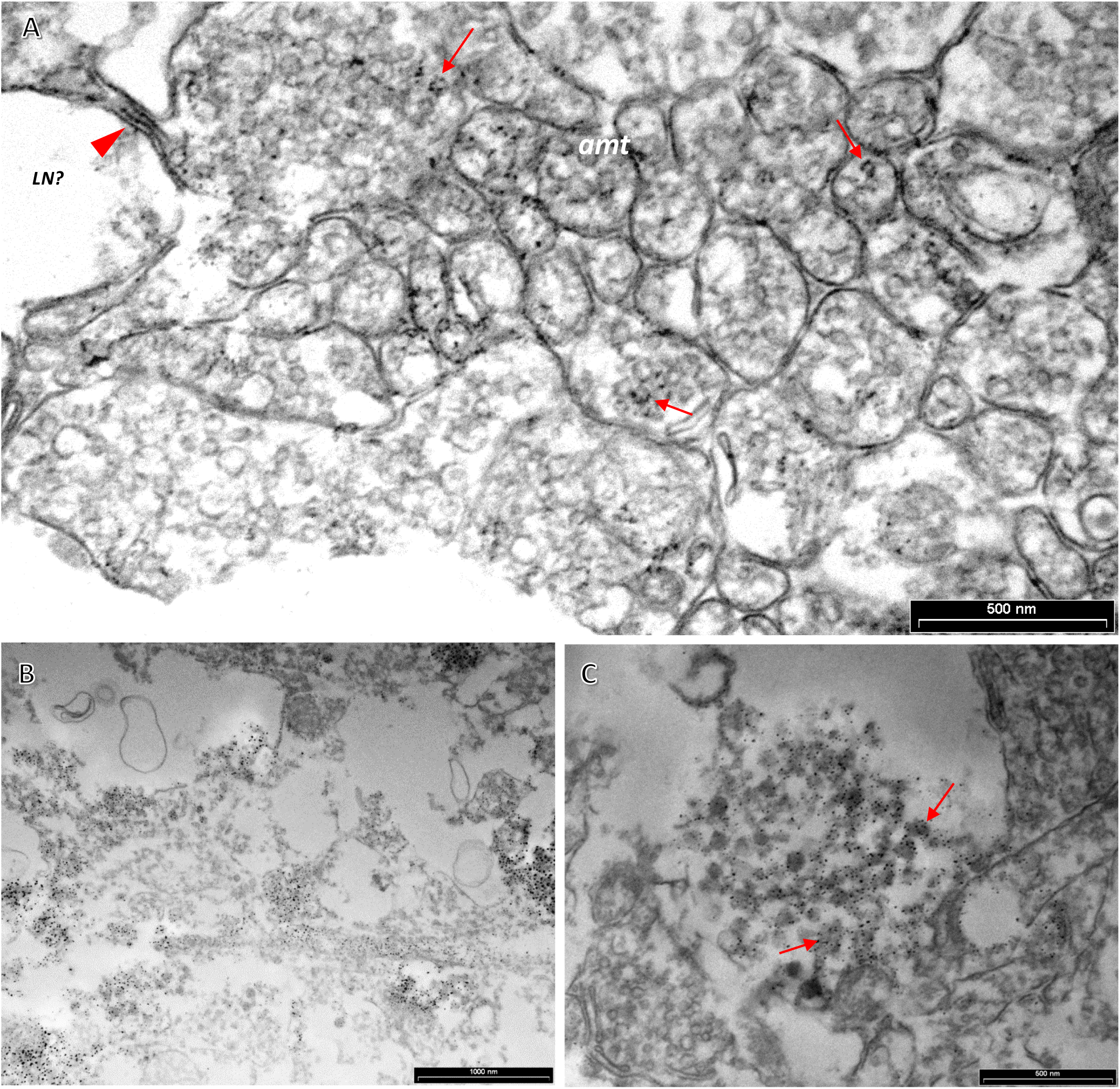
cChAT immuno-EM of neuronal profiles in the VL neuropil. (A) Electron micrographs showing cChAT-IR in agranular vesicles in profiles typical for amacrine trunks (*amt*) (arrows) in the inner neuropil (cut transversely). Note a pale unstained non-vesicular profile, possibly an LN (*LN?*), shows a synaptic connection density (thickened membrane, red arrowhead) with one of the positively stained trunks. (B) Fairly widespread cChAT immunoreactivity is possibly distributed in synaptic structures and processes in the neuropil mass. While it is not fully clear to which cells the labeled structures belong, neighboring profiles remain unlabeled. (C) Electron micrograph showing intensely labeled dark granulated vesicles of various sizes (arrows), which are probably not to be attributed to MSFL incoming fibers. Scale bar: 500nm (A, C); 1000nm (B).

**Fig. 9A** displays a relatively similar cChAT reactivity pattern in the MIFL and the MSFL neuropil and rather intense darker brownish labeling of the VL and subFL neuropils (**Fig. 9A’**, lower bar). Cholinergic properties in the visual and tactile learning systems appear comparable. Yet, possibly because of the larger cell bodies of the subFL cortex (up to ∼ 20µm), cChAT-IR cell bodies were clearly visible in the outer cell layers of the subFL (**Fig. 9A’**, upper bar).

**Figure 9.**
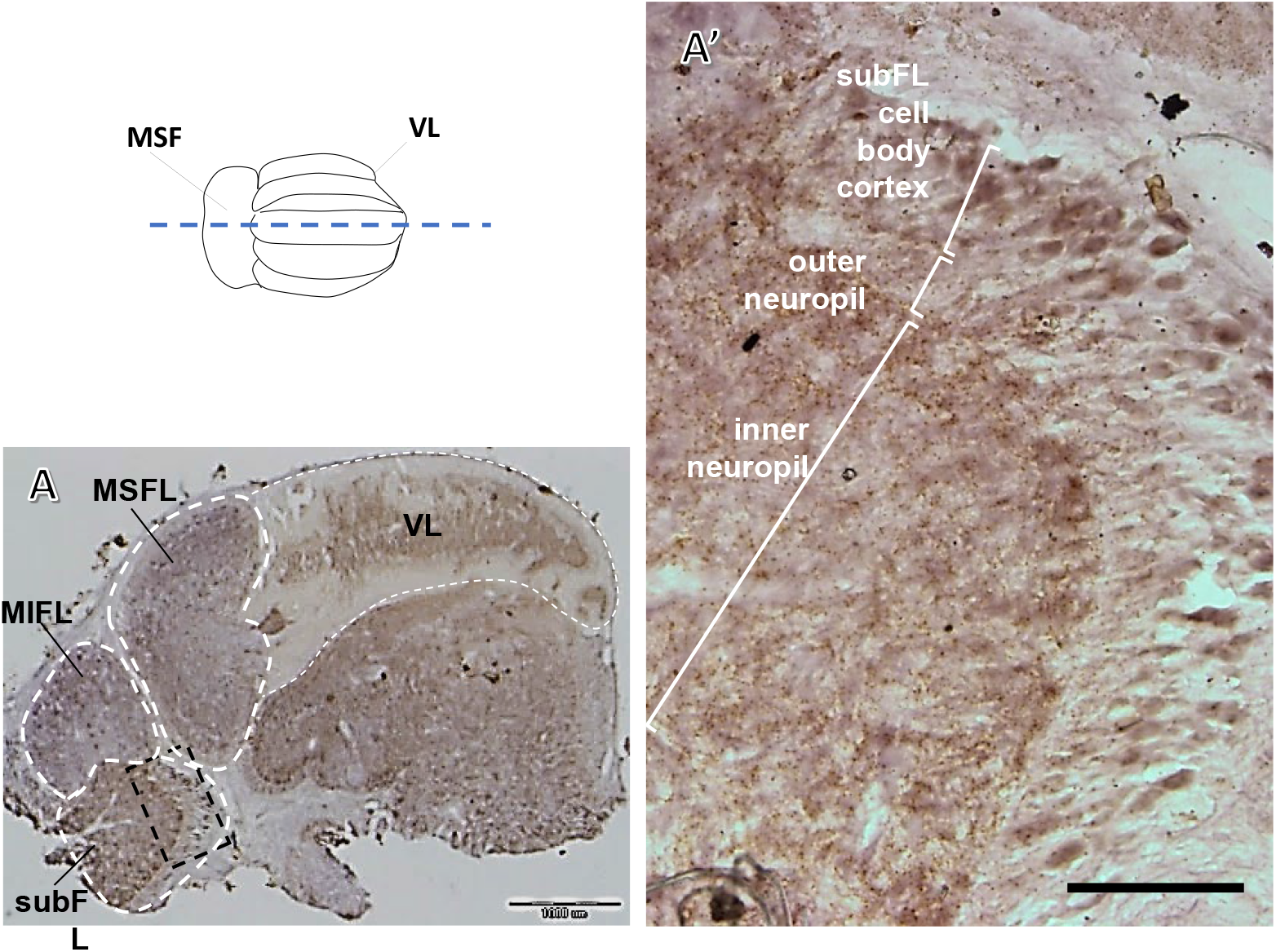
cChAT-IR distribution in the MIFL-subFL touch learning system. (A) Sagittal section from the middle area of the supraesophageal brain. Dotted rectangle gives the location of section A’. Note the similarity in intensity patterns of the MIFL and MSFL and in the intense, brownish labeling of the inner neuropil in VL and subFL (1:15000). (A’) Enlarged area of cell bodies and neuropil of the subFL. There is positive labeling of relatively large cell bodies (>20µm) localized in the outer cell cortex (top bracket). This location differs from that of the LNs mainly organized in the outer layers in the VL cell cortex. The MIFL-tract is barely apparent in this section (middle bracket), and dense labeling of processes is uniformly distributed in the inner neuropil (lower bracket). Scale bar: 1 mm (A); 200µm (A’)

#### γ-amino butyric acid (GABA)

**Fig. 10A** displays a general comparison between the GABA labeling in the MSFL, VL and the MIFL. GABA-IR processes were distributed mainly in the outer MSFL neuropil plexus. The intensely labeled processes appear to carry dark GABA-IR swellings, possibly synaptic varicosities (**Fig. 10A1’**). The dense and widespread GABA-IR probably derives from the external inputs to the MSFL described by Young (1971), rather than local MSFL innervation, suggesting inhibitory input to the MSFL. In both the MSFL and MIFL, the dense punctuated GABA-IR seemed to encircle fascicles of neuropilar structures (**Fig. 10A1, A3**).

**Figure 10.**
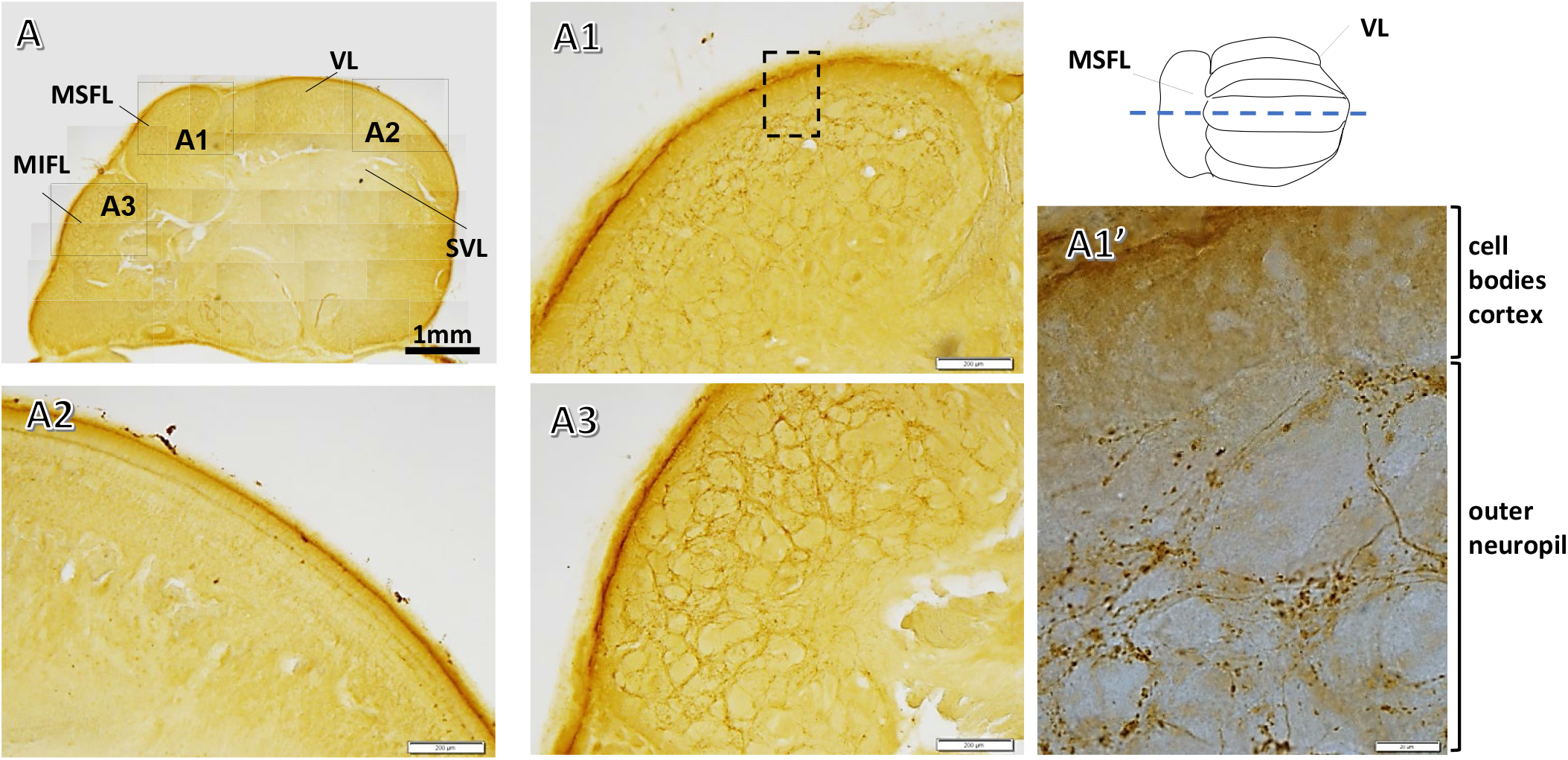
Light microscope micrographs of sagittal sections showing GABA-immunoreactivity labeling in supraesophageal lobes of the octopus brain. (A) Low magnification of supraesophageal brain preparation labeled for GABA, showing the MSFL (A1), VL (A2), and MIFL (A3) where specific labeling patterns were identified. (A1’) High magnification of the area marked in A1 showing GABA-IR labeling in the MSFL. No specific labeling was detected in the cell body cortex as opposed to the rich GABAergic area in the outer neuropil of the MSFL lobe with GABA-positive processes with dark varicosity-like labeling. GABA-positive processes and varicosities are scarcely detectable in the inner neuropil. Dark labeling of the sheath surrounding the brain and general faint staining was not observed in controls with secondary antibodies only (not shown). Scale bar: 1mm (A); 200µm (A1-A3); 20μm (A1’).

Strong GABA-IR labeling was detected in a distinct group of LN cell bodies (∼6-13 µm dia.) in the VL (**Fig. 11B**, arrows) and their neuronal processes (**Fig. 11B’**), which are remarkably organized along the dorsal and ventral inner margin of VL cell body cortex. These morphological characteristics fit the LN described by Young and Gray, and therefore our findings confirm that this group of LN mediates the inhibitory output of the VL (Shomrat et al., 2008). An intriguing, conspicuous, increased coloration was observed in the dorsal portion of GABA-stained slices, visualized as a thin, continuous darker sheath running at the border between the cell body cortex and the outer neuropil of the MSFL tract (**Fig. 11C**, rectangle).

**Figure 11.**
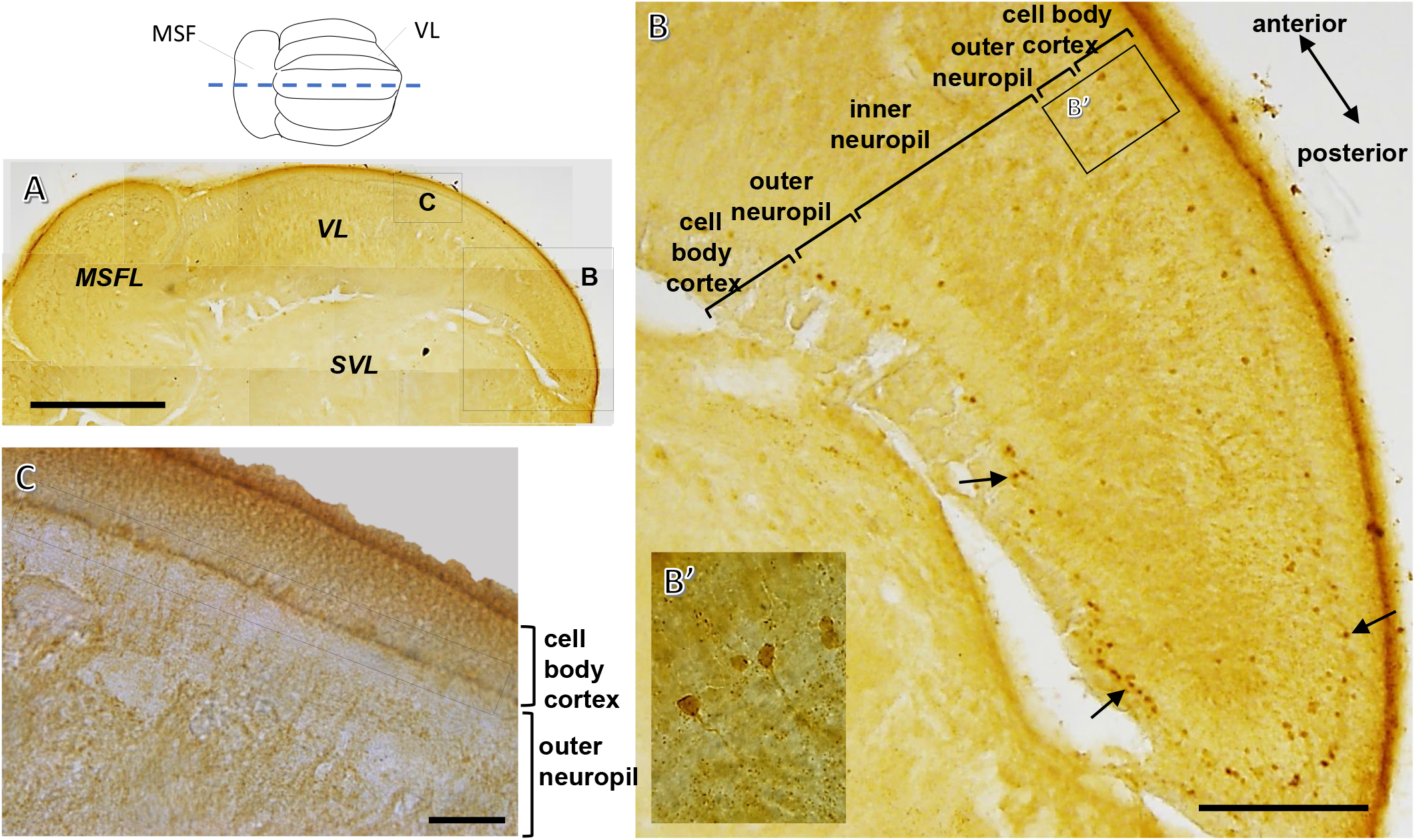
Light micrographs showing GABA-IR of large cell bodies in the VL. (A) Low magnification of the MSFL-VL system in a GABA-positive labeled sagittal slice, with locations of B and C marked. (B) The location and organization of GABA-IR large cell bodies (dia. 6-13 µm; arrows) around the inner dorsal and ventral margin of the cell body cortex strongly suggest that these are LNs. (B’) Higher magnification of individual GABA-IR large cells showing their positively labeled neurites emerging toward the inner neuropil, further indicating that these are ‘classical’ LNs. (C) A thin, anatomically yet undefined, dark layer runs along the border between the inner cell layer and the outer neuropil (rectangle). Scale bar: 1mm (A); 200µm (B); 50 µm (C).

Dense, punctuated GABA-IR was seen in processes in the middle-inner neuropil plexus, mainly in the medial lobule and medial-lateral lobes (**Fig. 12B**), while the outer neuropil, containing mainly fibers of the MSFL-tract, was scarcely labeled (**Fig. 12A**). **Figure 12B** shows a transverse section where such GABA labeling in the inner neuropil formed a distinct circular pattern with varicosities or thick granular labeling along the processes. These may represent presynaptic varicosities conveying inhibitory input, possibly to the LN dendrites. Such inhibitory inputs may originate from other LNs, GABAergic AMs (see below), or inhibitory afferents from the subVL, such as “pain fibers” described by Young (1971).

**Figure 12.**
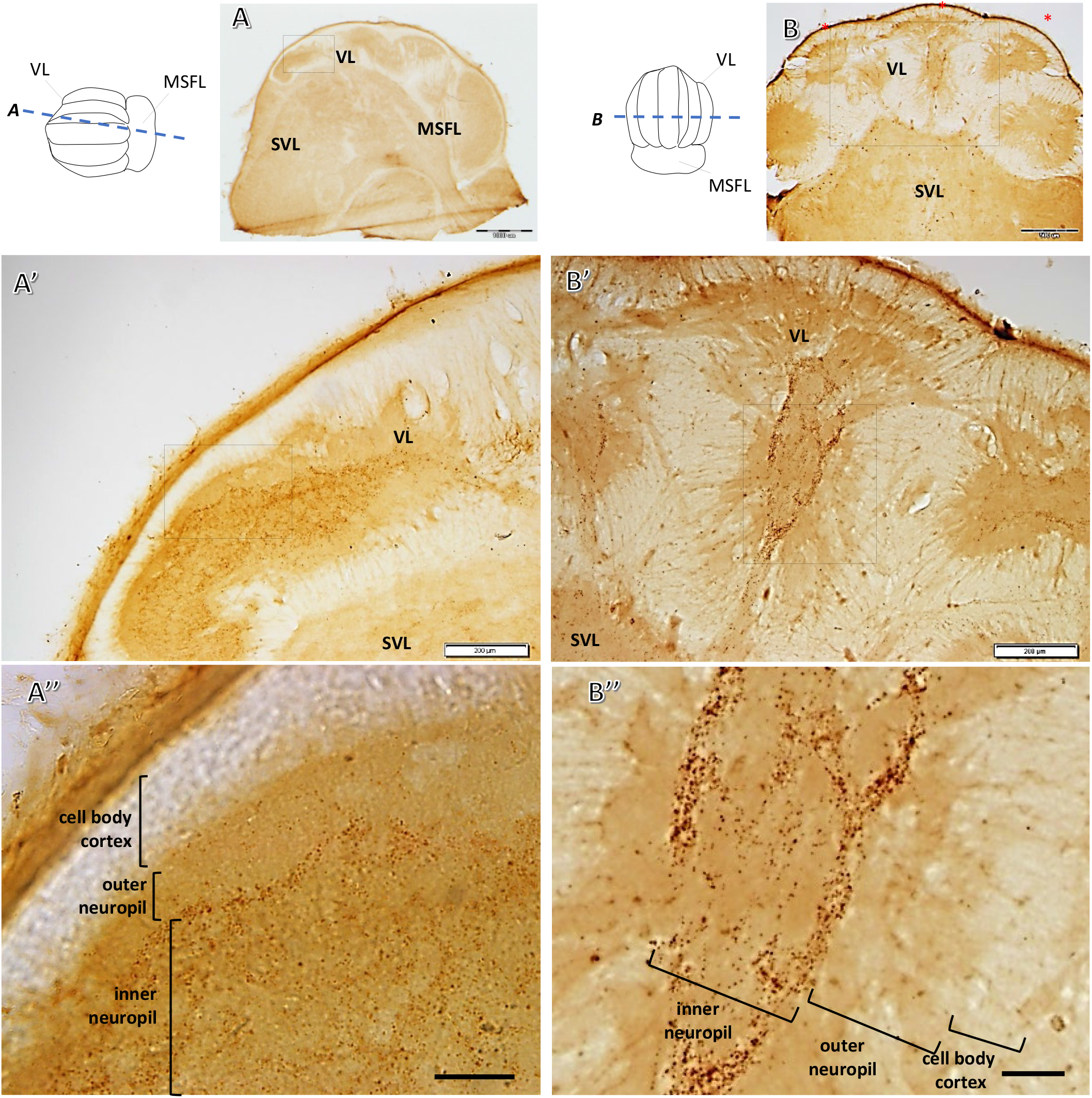
GABA-IR reveals a unique dense labeling pattern in a distinct area in the inner neuropil. (A) Low magnification of a semi-sagittal slice labeled for GABA, including the MSFL-VL system. Location of A’ is marked. (A’) GABA-IR labeling in the inner neuropil. (A’’) Higher magnification of the area marked in A’ showing barely noticeable GABA-IR in the AM cell layer and in the outer neuropil (MSFL-tract), while punctuated dark labeling is seen in the inner neuropil. (B) Low magnification of a transverse slice labeled for GABA, including the five VL lobuli. (B’) Enlargement of the area marked in B. The GABA-IR observed in transverse section forms a distinct loop-like pattern in the inner neuropil. (B’’) Enlargement of the area marked in B’ shows that the dense labeling carries varicose-like markings suggesting this area comprises GABAergic synaptic terminals. Scale bar: 1mm (A); 500µm (B); 200µm (A’, B’), 50µm (A’’, B’’)

Although physiological finding suggested that the excitatory input to the LNs derives from cholinergic AMs (Shomrat et al., 2011), GABA-IR was clearly seen in the region of the AM cell bodies in the VL cell cortex (**Fig. 13**). While unclear staining of the cell somata hindered verification that some of the AMs are indeed GABA-positive, GABA-IR neurite processes were clearly evident running from the AM cell body area and crossing the unlabeled outer neuropil (MSFL tract; **Fig. 13A,A’**). GABA-positive projections crossing the VL hila can be clearly seen in the medial and medial-lateral lobuli (**Fig. 13B, B’**, arrowhead).

**Figure 13.**
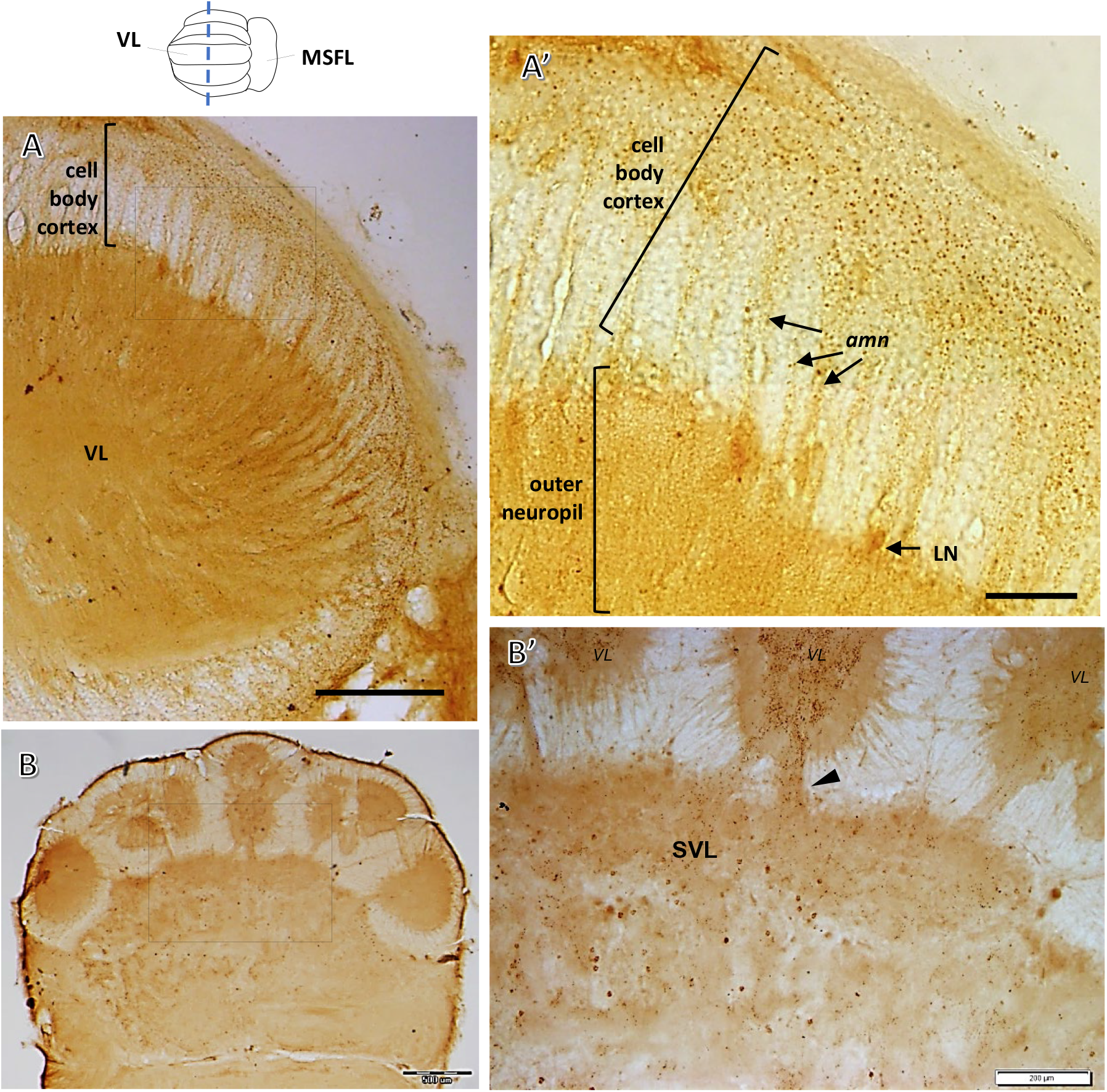
Light micrographs show GABA-positive processes crossing the cell body cortex within and outside the VL. (A) Low magnification of a transverse section of a lateral lobule labeled for GABA. The location of A’ marked is. (A’) Higher magnification of a lateral lobule showing GABA-IR in bundles of processes crossing the apparently unlabeled MSFL axons in the outer neuropil with consistent gaps of approximately 20µm between them (arrows). These processes appear to reach both outer cell layers and the VL neuropil and are most likely AM neurites (*amn*). Large cells positively labeled for GABA are situated in the inner margin of the cell layer cortex (arrow). (B) Low magnification of a transverse section of the supraesophageal brain. Location of B’ is marked. (B’) GABA-IR in processes (arrowheads) connecting the VL with subVL. Scale bar: 200µm (A); 50µm (A’); 500µm (B); 200µm (B’).

Support for selected GABAergic AMs was provided by EM observations in which GABA positivity was detected in what were most likely AM neurite bundles and synaptic connections (**Fig. 14B**). EM also revealed neuronal processes containing swarms of GABA-IR clear agranular vesicles (*cv*), 30-80 nm dia., associated with amacrine cells (**Fig. 14C**). In some cases, positive synaptic vesicles were seen neighboring unlabeled synaptic terminals (**Fig. 14D, E**). Taken together, these findings show that at least some of the AMs can be GABAergic and thus may provide the inhibitory inputs to the LNs (Shomrat et al. 2011 and unpublished results).

**Figure 14.**
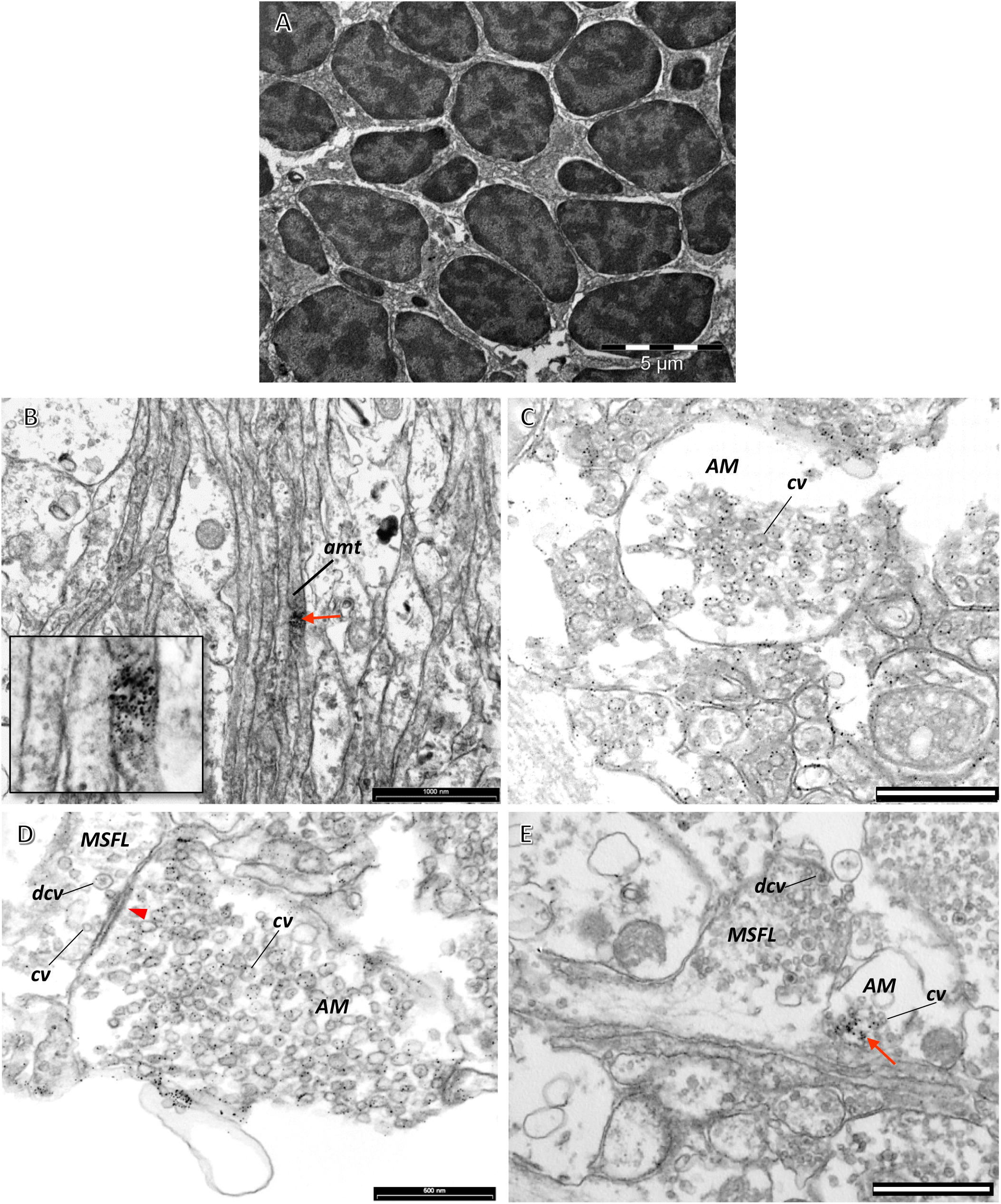
Electron micrographs showing GABA-IR in the VL. (A) Section showing the closely packed cell bodies of the small AMs. The nucleus occupies most of the cell body. (B) A longitudinal section in the outer neuropil. Although GABA-IR was not identified in the cell body cortex, labeling was observed in a process (red arrow) crossing over MSFL axons. (C) Section of an ovoid varicosity. GABA-IR can be seen in agranular clear vesicles (cv), associated with AMs ranging from 30-80 nm dia. (D) Neighboring cells sampled from the outer neuropil area. The cell containing cv (probably AM) reveals GABA-IR, while no labeling was identified in the adjacent MSFL varicosity containing cv and dense core vesicles (dcv). Note the membrane thickening (arrowhead) suggests a synaptic contact between the two but without clear directionality. (E) GABA-IR in cv docked near the membrane of a cell neighboring an MSFL axon synapse, which hints at inhibitory synapses in the area of the MSFL axon terminals. Scale bar: 5µm (A); 1µm (B, E), 500nm (C, D)

**Figure 15A,B** shows abundant GABA-IR matrix-like fibrous labeling in the medial inferior frontal lobe (MIFL) similar to that in the MSFL (cf. **Fig. 10A1**). The subFL clearly exhibited large GABA-IR cells distributed in the inner cell body cortex and positive granular processes throughout the neuropil (**Fig. 15C-E**), remarkably resembling the labeling patterns of the VL (**Figs. 11, 12**).

**Figure 15.**
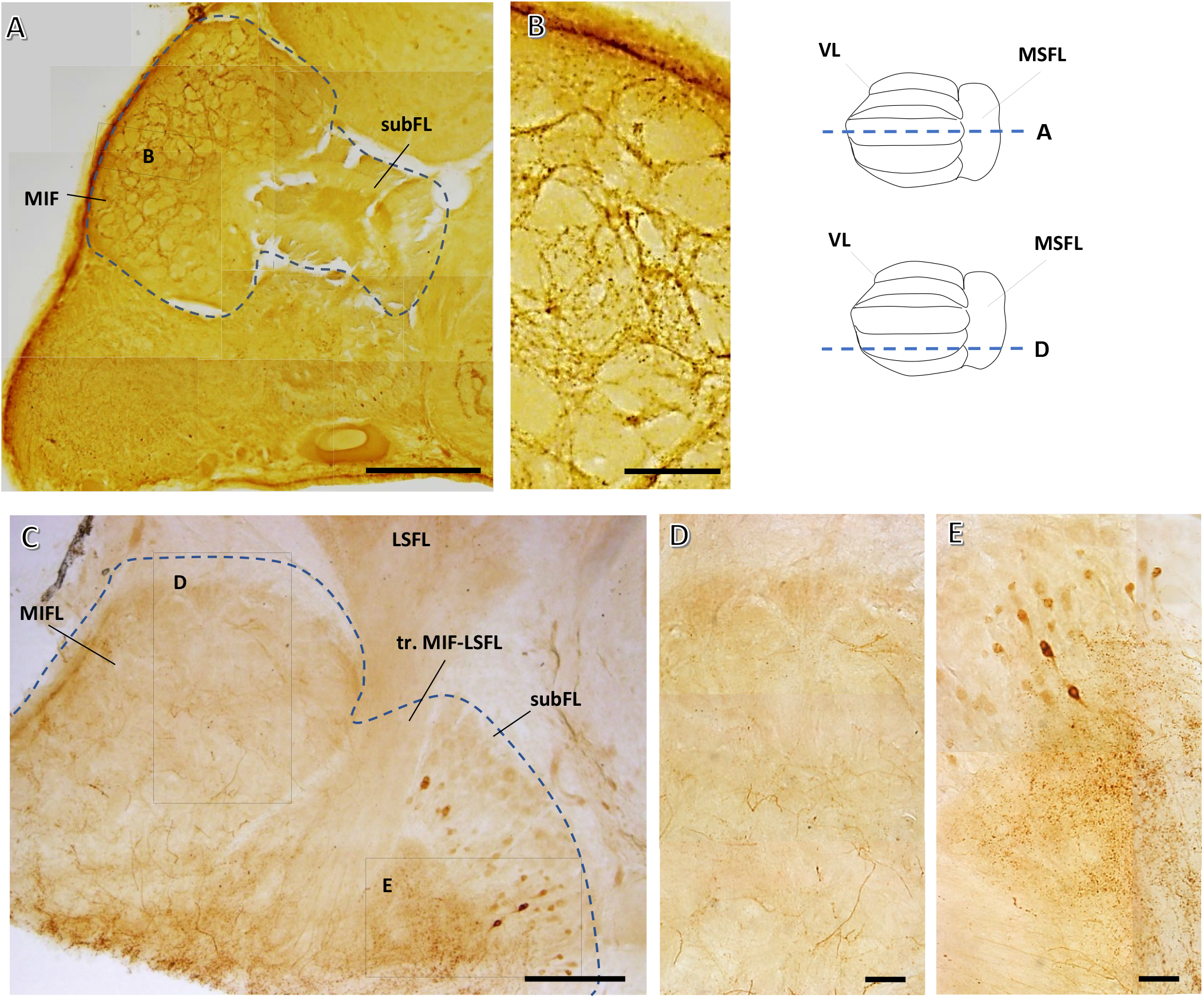
GABA-IR distribution in the touch learning system. (A, C) Sagittal GABA-labeled sections of the MIFL-subFL. Marked areas are shown at higher magnification in B, D, and E. (B) Prominent labeling of the MIFL neuropil comparable to the labeling pattern in the MSFL (see Fig. 10). (D) A more lateral section in which the labeled processes are still apparent but less intense. (E) The subFL mass clearly demonstrates large cell GABA-IR in the inner cell body cortex and positive labeling of the neuropil mass comparable to the labeling patterns in the VL (see Fig. 11). Scale bar: 500µm (A); 100µm (B-E)

Using ISH of the metabotropic GABA-B-like receptor mRNA gave especially strong expression in the MSFL cell cortex (**Fig. 16A**). In contrast, there were only some pale expressions in the VL, mostly in the dorsal cell body cortex and in the inner margin of the ventral cell body cortex (**Fig. 16B**, arrows). The cell cortex of the MIFL revealed GABA-B receptor transcript expressions similar to those in the MSFL cell cortex, and the faint expression in the SubF cell cortex corresponded with that in the cell layer of the VL (**Fig. 16A**).

**Figure 16.**
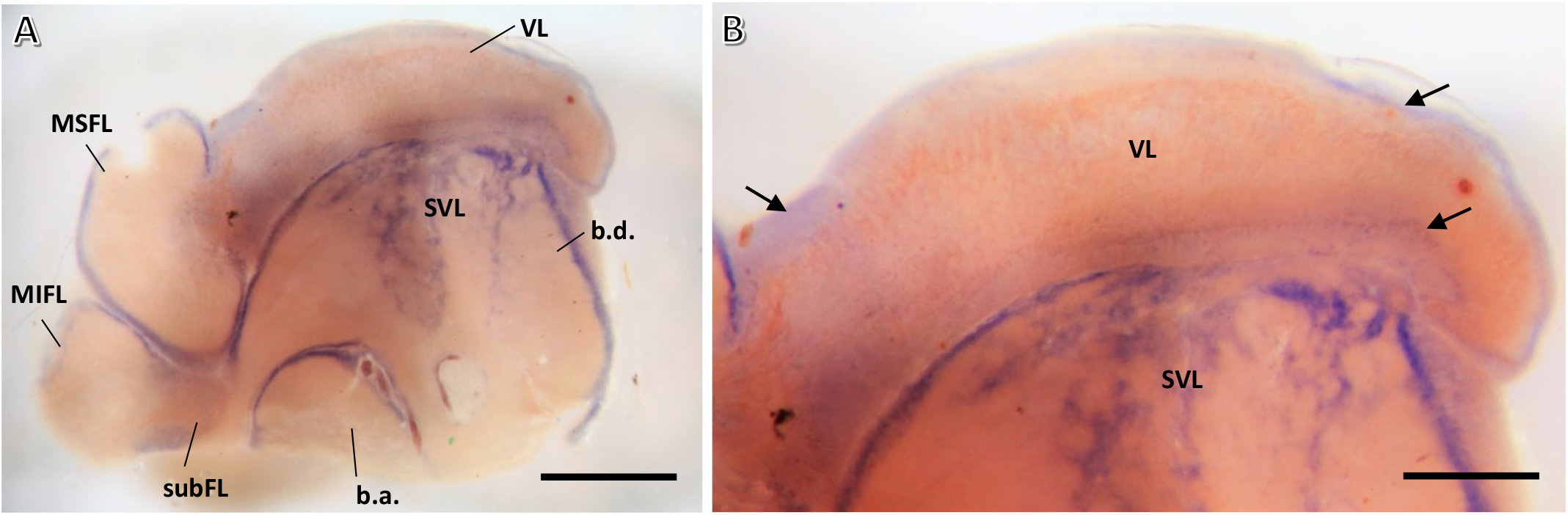
Photomicrographs of GABA-B expression detected by *in situ* hybridization in supraesophageal lobe sections. (A) Cells strongly expressing GABA-B mRNA can be seen in the MSFL cell cortex, SVL, and the basal lobes. (B) Higher magnification of the VL reveals faint expressions of GABA-B mRNA in the cell body cortex of the VL. The VL and the subFL appear to express lower levels of GABA-B receptors than the other lobes. Scale bar: 1mm (A); 500µm (B)

### Putative Neuromodulators

#### Recruitment of Nitric Oxide (NO) in the synaptic plasticity pathways

Fixative resistant NADPH-diaphorase reactivity is a reliable marker of NOS activity in molluscan preparations (Moroz et al. 2005; Floyd et al. 1998; Moroz et al. 1999). Fitting with the involvement of NO in LTP in the *Octopus* VL (Turchetti-Maia et al. 2018), intense NADPH-d staining revealed NOS activity in the neuropil of all five VL lobuli (**Fig. 17A,B,C1,C2**) and in the subFL (**Fig. 17A,E**). In the VL, staining was found in the inner zones where the synaptic connections between the AM and the LNs lie. The outer neuropil was more sparsely stained, probably because in this region unlabeled axons that run in the MSFL tract make sparse *en passant* connections with AM neurites. The AM neurites can be seen crossing the tract in faintly stained AM trunks with gaps of 2-10µm between them (arrows **Fig. 17D1**), suggesting the presence of NOS in the AM neurites. The patterns of labeling in the VL and the subFL were extremely similar (**Figs. 17D1,D2**). Note the clear but sparse labeling of cell bodies in the subFL cortex (**Fig. 17F** arrows).

**Figure 17.**
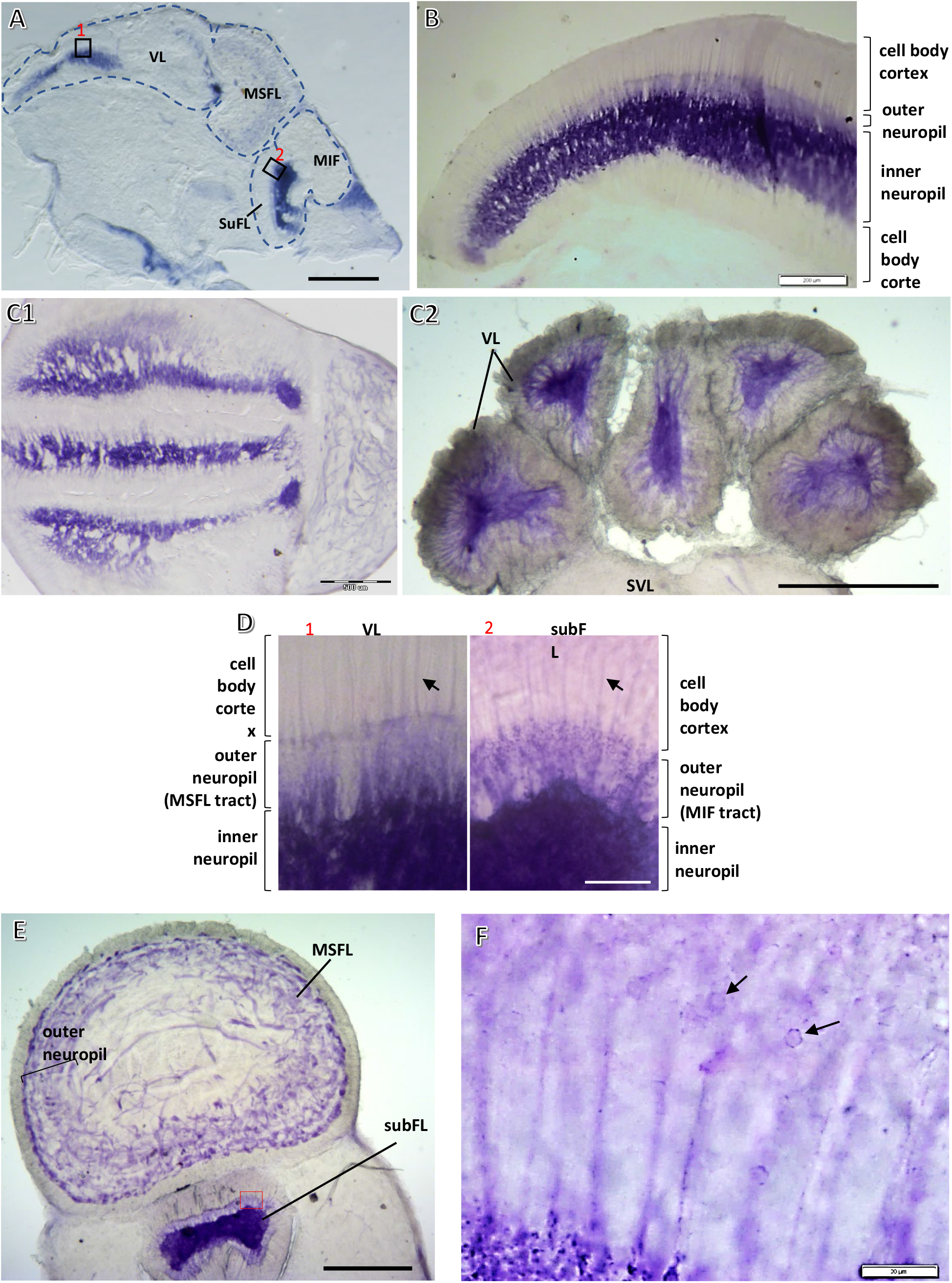
Intense NOS activity in the MSFL-VL and MIFL-subFL systems revealed by NADPH-d labeling in the supraesophageal lobes. (A) Sagittal section with intense labeling of the neuropil in VL and SubFL. (B) NADPH-d labeling in the VL showing different staining patterns in the cell body cortex, outer and inner neuropil. (C1,2) Horizontal (C1) and transverse (C2) sections of the VL lobuli showing a similar pattern of positive NADPH-d labeling in the different lobuli. (D) Higher magnification of the areas marked in A showing comparable labeling patterns between brain structures involved in visual learning (VL) and tactile learning (SuFL) systems. (E) Transverse section of the MSFL and SubFL area showing positive labeling. In the MSFL the staining in the outer neuropil is darker than in the center. (F) Enlargement of area marked in E (subFL) showing positive NADPH-d neurites and cell bodies (arrows). Scale bar: 500µm (A, C1, C2, E); 200µm (B); 50µm (D); 20µm (F) [Figs. A,C2,D from Turchetti-Maia et al. 2018].

The MSFL neuropil also contained NADPH-d positive processes (**Fig. 17E**), although staining was much less abundant than in the VL and subVL (**Fig. 17A**). Staining was distributed throughout most of the MSFL neuropil, especially at a thin layer below the cortex. Scarce labeling was seen in the plexiform arrangement in the deeper central region of the lobe (**Fig. 17E**), an area containing mainly incoming fibers (Young 1971).

#### Catecholaminergic system: tyrosine hydroxylase (TH)

TH-positive labeling showed prominently in a group of neurons with large cell bodies (>10 µm dia.) in the posterior MSFL lobe, bordering, or possibly belonging to, the VL (**Fig. 18A, B**). This cluster formed a ‘deep nucleus’, contrasting with the regular organization of cell bodies in an outer cortex. TH-IR could be followed for some distance in neuronal processes (**Fig. 18A’’**). Similar cells, in a similar location, also showed L-glutamate-IR (**Fig. 5**), and formed symmetrical tree-like structures feeding into the anterior region of the VL neuropil (**Fig. 18A, A’**). Only very few TH-positive large cells (∼12.5 µm dia.) were detected in the VL cortex; these lay in the inner margin of the VL cell cortex, where the LNs lie. Their projections also showed positive TH labeling and could be followed inward to the depth of the VL neuropil (**Fig. 19 A’**, arrows). It is not yet clear if these dopaminergic (or other catecholamines) neurons are special efferent LNs or constitute an internal neuromodulatory system (see below).

**Figure 18.**
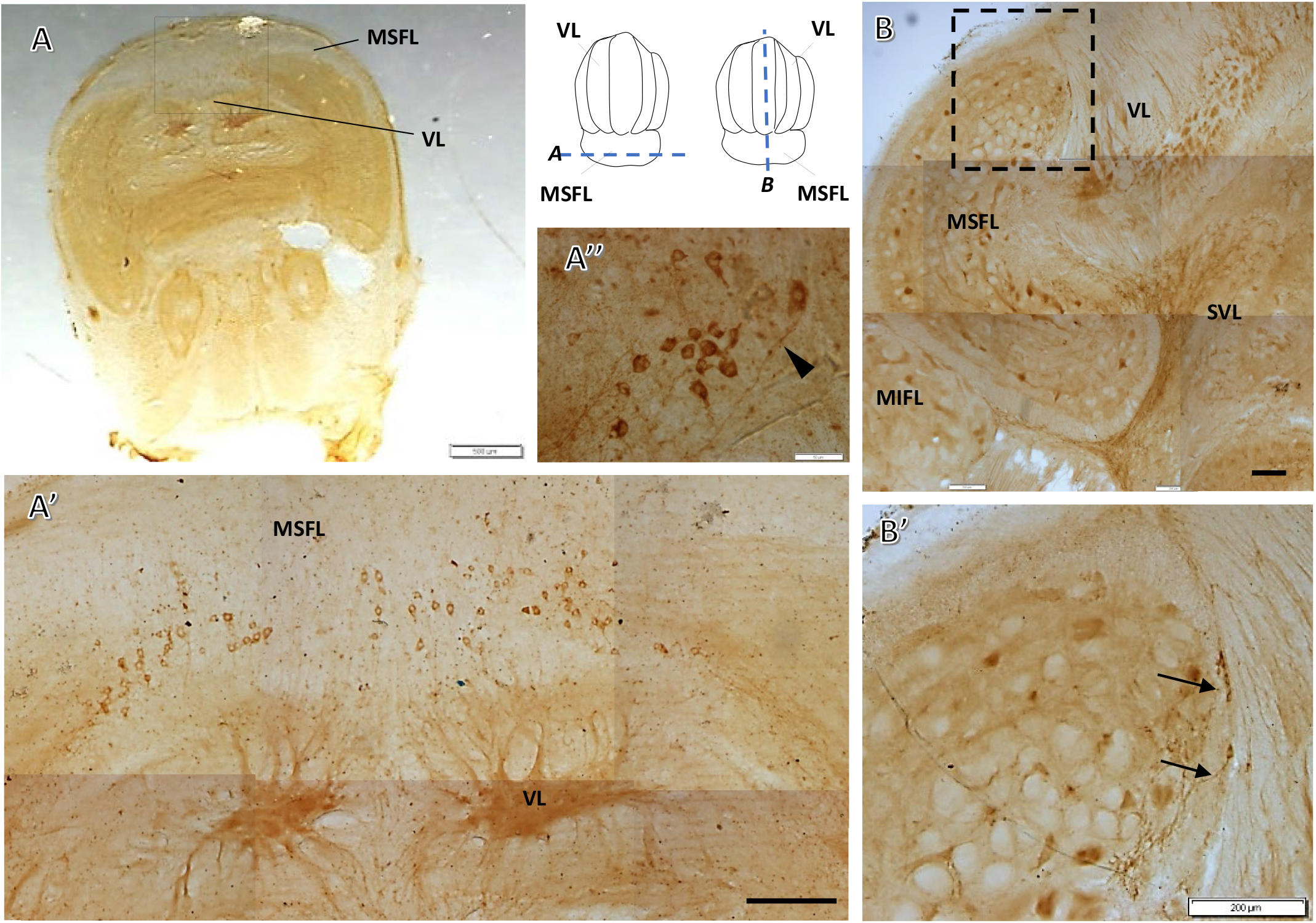
Tyrosine hydroxylase (TH) immunoreactivity reveals a “deep nucleus” in the VL-MSFL system. (A) Transverse TH-labeled slice from the border area between the MSFL and the VL. The location of A’ is marked. (A’) Grouped TH-IR cell bodies localized in the anterior VL area/posterior MSFL organized as a ‘deep nucleus’ (cell dia. ∼16-20 µm). (A’’) TH-IR projections from cells localized in the deep nucleus can be followed a certain distance (arrowhead; image from slice similar to A). (B) TH-labeled sagittal slice of the MSFL and neighboring areas. Location of B’ is marked. (B’) Higher magnification showing several cells that probably belong to the ‘deep nucleus’ (arrows). Scale bar: 0.5mm (A); 200µm (A’, B, B’); 50µm (A’’)

**Figure 19.**
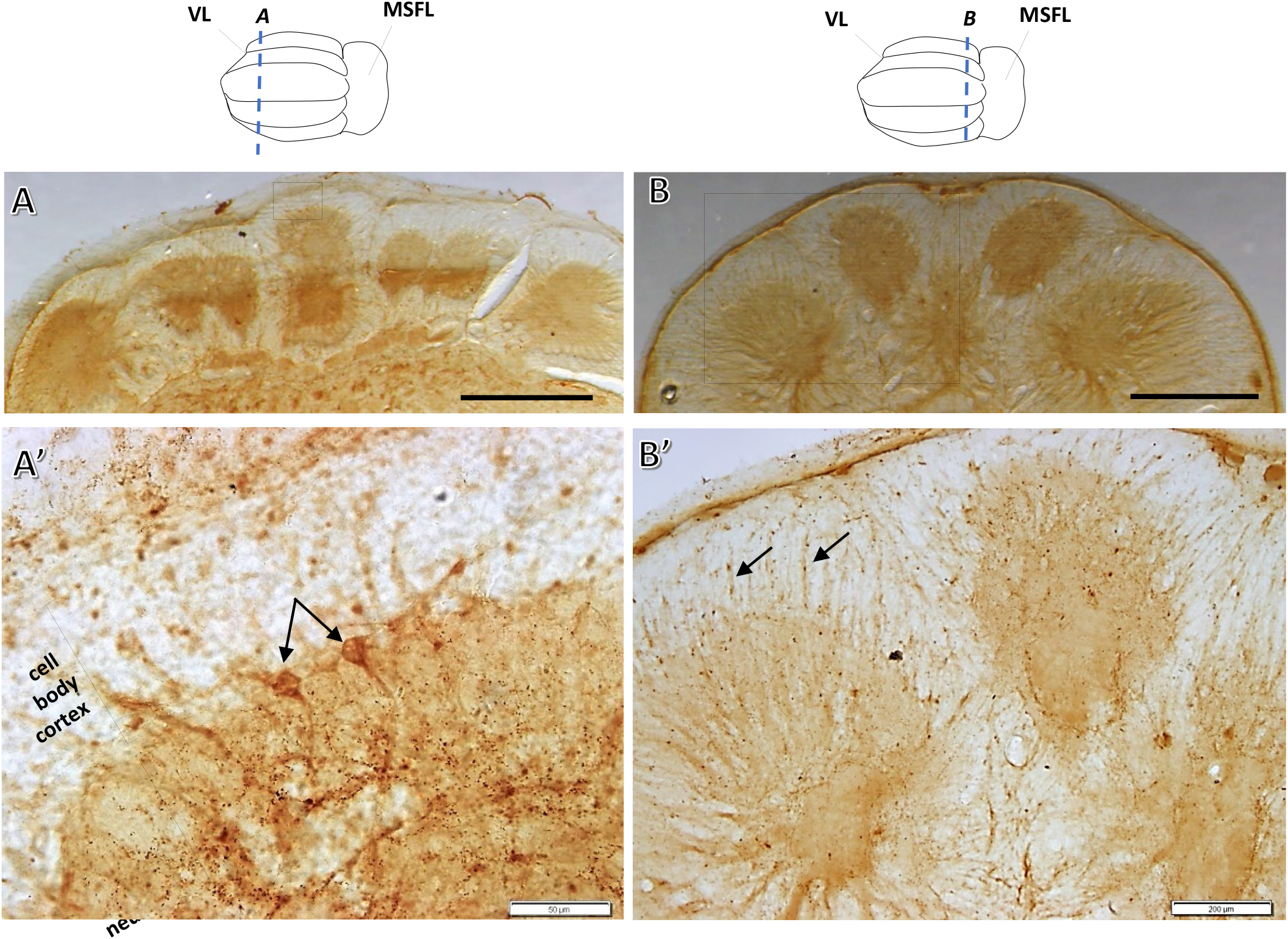
Scarce TH immunoreactivity in the cell body cortex of the VL. (A, B) Transverse TH-labeled slices. Locations of A’ and B’ are marked. (A’) TH-positive large cell bodies (∼13 µm dia.) located in the inner layers of the VL cell cortex. They project inwards to the neuropil. (B’) Transverse view of part of the VL lobuli showing faint staining of neuronal processes running between the cell body layers and the neuropil (arrows). There is clear granular labeling in the neuropil (see also Fig. 21). Scale bar: 500µm (A,B); 50µm (A’); 200µm (B’)

TH immunoreactivity was evident in both the inner and outer VL neuropil. The outer neuropil showed TH-positive processes interweaving in the lower part of the cell layer (**Fig. 19B,B’** arrows, **Fig. 20A**), while in the dorsal and ventral aspects of the inner neuropil a dark TH-IR impressively defined a thickened band carrying TH-positive fibers with varicosities along their length (**Fig. 20A**). Correspondingly, ring-like thickenings were seen in transversal sections (**Fig. 20B, B’**), suggesting dopaminergic innervation of a specific dendritic area of, for example, an LN. These ring-like patterns in the central inner neuropil and the VL-subVL crossing fibers were evident mainly in the medial and medial-lateral lobuli, while only scarce TH-IR fibers were seen crossing through the VL hila of the lateral lobuli. The spatial distribution of the TH-IR at the center of the inner VL neuropil suggests that the source of some of the labeled twigs may be afferent processes from ventral lobes originating or crossing through the subVL to the VL. Additional sources for these processes are the TH-positive cells organized as a ‘deep nucleus’ at the MSF-VL border (**Fig. 18**) and the TH-IR LN in the VL cell cortex (**Fig. 19A**).

**Figure 20.**
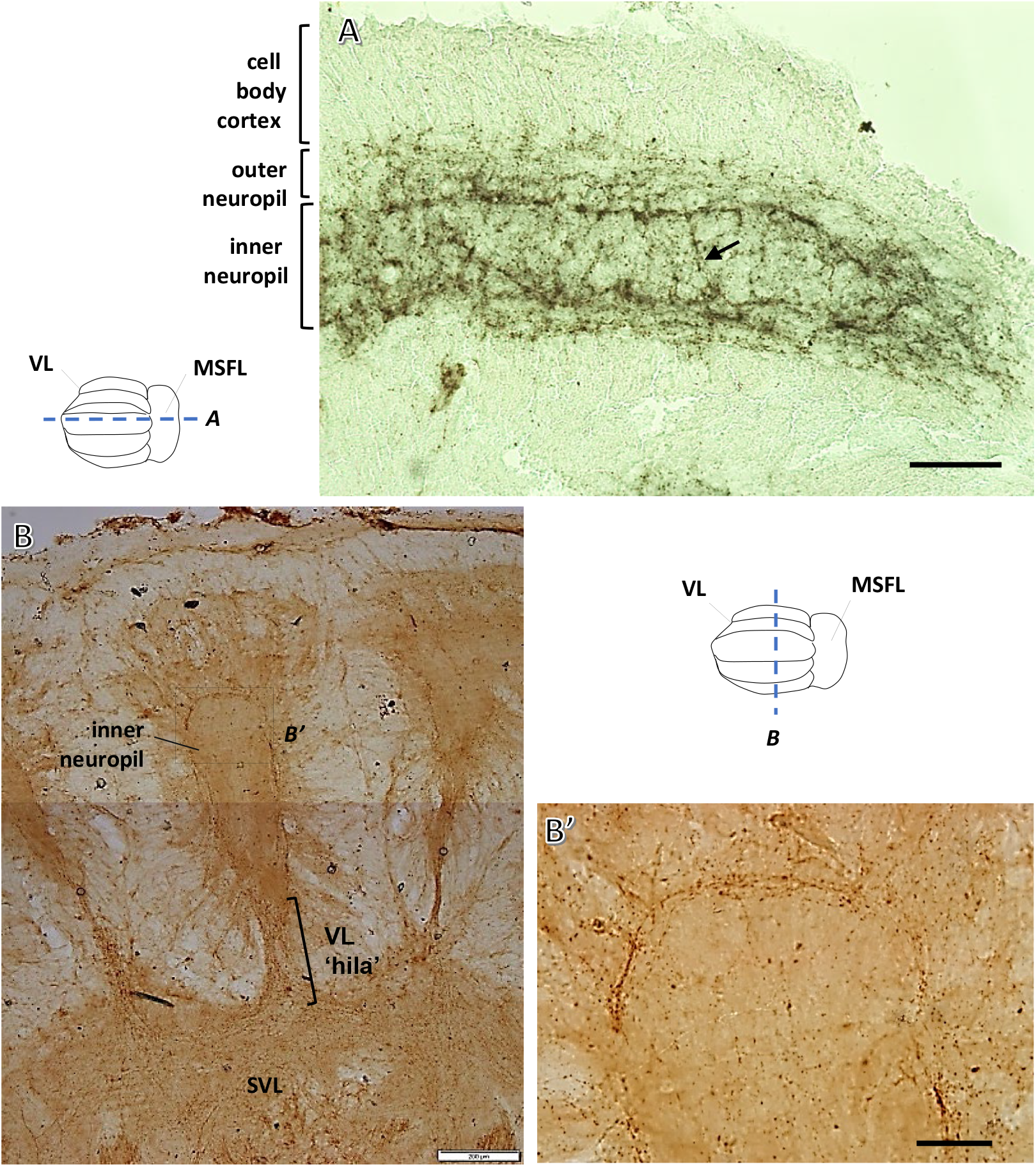
TH-immunoreactivity in the VL neuropil. (A) A sagittal slice of the VL showing TH-IR in both the inner and outer neuropil but no clear TH-IR cell bodies in the AM cell body layer. The inner neuropil is characterized by an impressively defined TH-IR ring-like band carrying positively stained thickenings and varicose fibers along its length. (B) The TH-IR ring-like labeling in the inner medial lobule neuropil (transverse section) corresponding to the labeling in A. TH-IR processes in SVL-VL tracts crossing the VL hila were observed mainly in the medial and medial-lateral lobes. Note the bottleneck-like morphology where the labeling is especially dense (brackets). Location of B’ is marked. (B’) TH-IR dense ring-like labeled processes in the inner neuropil. Scale bar: 200µm (A, B); 50µm (B’)

#### Predicted Neuropeptides

The expressions of some well documented molluscan neuromodulatory neuropeptides, such as FMRFamide-like neuropeptides (FLPs), buccalin, bradykinin and conopressin peptide were localized in the VL using ISH (**Figs. 21,22** and see (Winters 2018). FLRIamide-encoding transcript expression (**Fig. 21A1,A2**) was distributed in a distinctly organized pattern, clearly highlighting large cells bodies dispersed uniformly in the inner margin of the cortex in all five lobuli where the LNs lie and where GABAergic and glutamatergic LNs were found. The unique pattern of labeled cells expressing FLRIamide transcripts suggests that at least some of the LNs express FMRFamide-related peptide (FaRP) as a cotransmitter in addition to their conventional fast transmitters.

**Figure 21.**
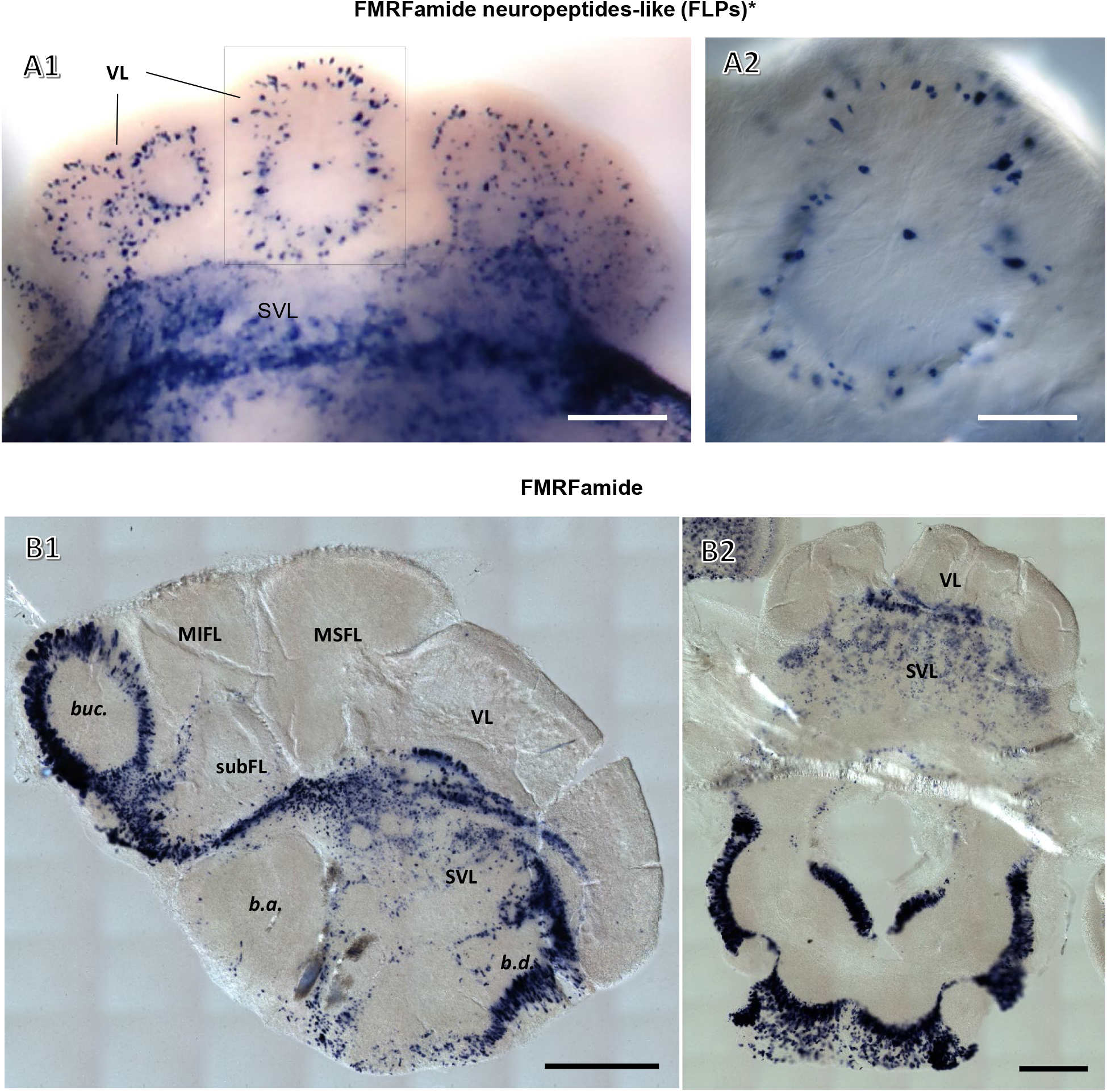
*In situ* staining of neuropeptide mRNA-expressing neurons in the VL system and other supraesophageal regions - FMRFamide neuropeptides-like. (A1) Transverse section showing FMRFamide-like peptide (FLRIamide) mRNA-expression in large cell bodies resembling LNs. The SVL also shows strong transcript expression. (A2) Enlargement of the area marked in A1 showing FLRIamide-expressing LNs in the medial lobule. (B1) Sagittal section showing FMRFamide mRNA-expressing cells in the supraesophageal brain. In addition to the abundant expression throughout the SVL, dorsal basal lobe (b.d) and buccal lobe (buc), some cells were detected in the ventral cell body cortex of the VL and subFL. (B2) Transverse slice showing FMRFamide mRNA-expressing cells in the SVL and subesophageal brain areas. Sparse labeling is also seen in the ventral cell cortex of the VL lobuli. Scale bars: 500µm (A1, B2); 1mm (B1), 200µm (A2)

**Figure 22.**
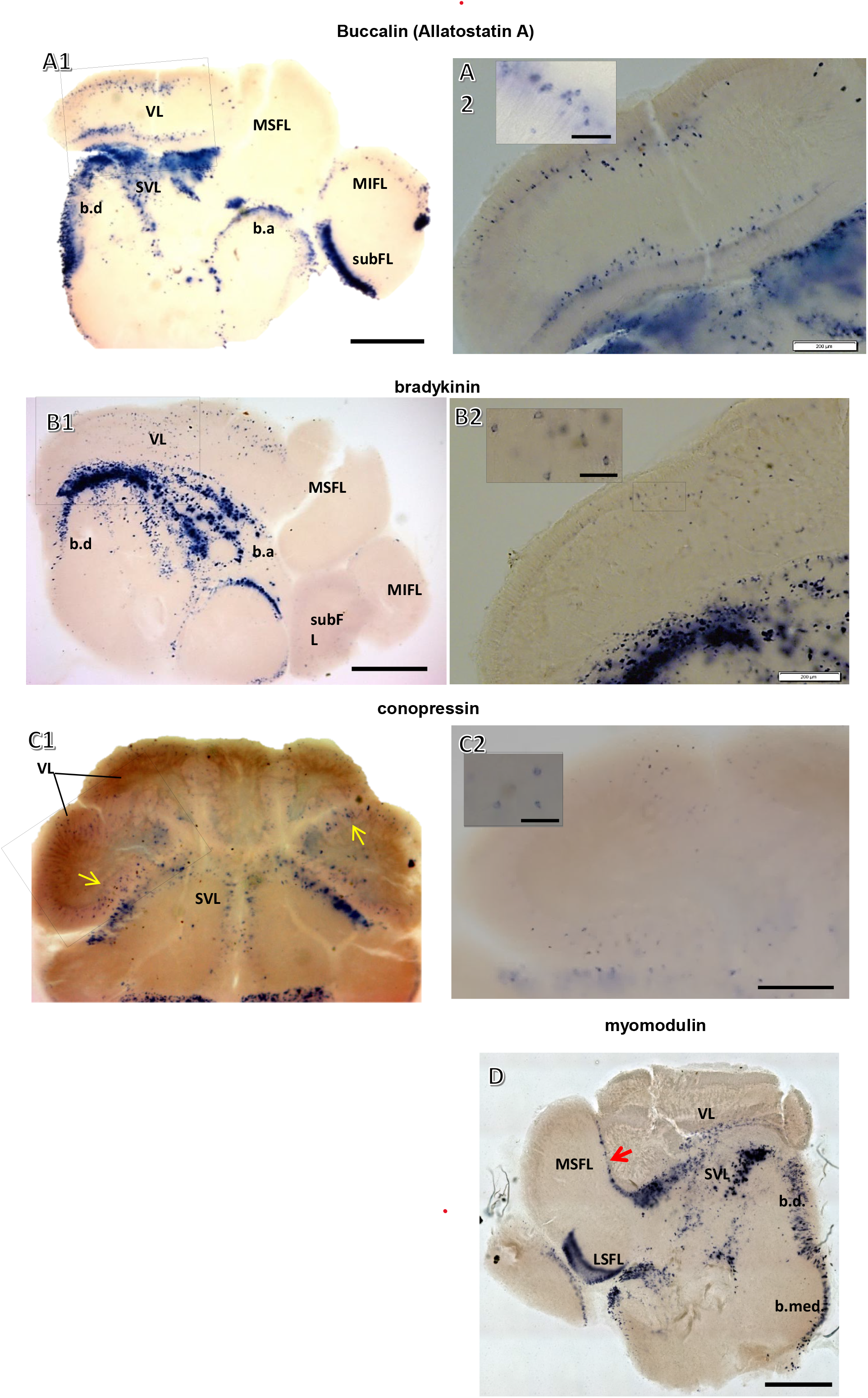
*In situ* staining of neuropeptide mRNA-expressing neurons in the VL system and other supraesophageal regions – buccalin, bradykinin, conopressin, myomodulin. (A1) Sagittal section showing buccalin mRNA-expressing neurons in the dorsal and ventral cell layers in the VL. There is also marked expression in the SVL, dorsal and anterior basal lobes and subFL cortices. (A2) Enlargement of the area marked in A1. Buccalin mRNA expressing individual cells are enlarged in the inset image. (B1) Sagittal section showing bradykinin mRNA-expressing cell bodies in the VL. Expression is especially strong in the SVL. (B2) Enlargement of the area marked in B1. Individual cells expressing bradykinin mRNA are enlarged in inset image. (C1) Transverse section showing conopressin mRNA-expressing neurons mainly, but not exclusively, in the lateral lobuli of the VL (arrows). (C2) Enlargement of the area marked in C1. Individual bradykinin-expressing cells are enlarged in inset image. Intense expression was observed in cells throughout the SVL. (D) Myomodulin transcript-expressing neurons are present in the border between the medial SFL (MSFL) and the VL (arrow), in the SVL, basal lobes and lateral SFL (lSFL). There were no clear indications for expression within the VL itself. Scale bar: Scale bars: 500µm (A1, B2); 1mm (A1,B1,C1,D); 200µm (A2,B2,D2); 50µm (Insets)

FMRFamide mRNA expression was seen in the MSFL at the ventral areas of the VL lobuli, MIFL and subFL. Cell expressions of FMRFamide-encoding transcripts were more widely distributed in other supraesophageal lobes, including the subVL (**Fig. 21B1,B2**).

Buccalin mRNA was identified in the inner margin of the VL cell body cortex but not in the MSFL (**Fig. 22A1,A2**). As reported in (Winters 2018) cells expressing bradykynin mRNA were found in the VL cell cortex, but not in the MSFL (**Fig. 22B1,B2**). Cell bodies expressing bradykinin were also scattered in inner areas of the VL, in what appeared to be cortical folds between lobuli. Conopressin mRNA was expressed in cells throughout the cell body cortex of the VL lobules, especially in the lateral lobules (**Fig. 22C1,C2**). In contrast to the scattered distribution of bradykinin and conopressin mRNAs in the VL, these transcripts were expressed in district areas of other regions of the brain.

Myomodulin peptide mRNA was expressed in cells at the MSFL-VL border (**Fig. 22D**), seemingly belonging to the cell bodies in the ‘deep nucleus’ (see **Fig. 3**, **Fig. 18**). Some positively labeled cells were also seen at the VL ventral cell cortex-subVL border.

## Discussion

The VL is arranged as a matrix in a fan-out fan-in network configuration (see Fig. 1C), in which the incoming MSF axons innervate *en passant* a very large group of minute amacrine interneurons (AM). These, in turn, converge onto a relatively small group of large efferent neurons (LNs; Gray, 1970; Young, 1971). Physiological experiments (Hochner et al., 2003, Shomrat et al., 2011) revealed an excitatory feedforward connectivity in which MSF afferents connect to the AMs via excitatory glutamatergic-AMPA-like receptor type synapses (Hochner et al. 2003), while the AMs show converging cholinergic excitatory synaptic connections to LNs. Plasticity is expressed as a robust activity-dependent LTP at the glutamatergic MSFL-AM connections. This LTP is important for behavioral learning and acquisition of long-term memory (Shomrat et al. 2008).

Even after decades of study the neuroanatomical distribution of neurotransmitters and neuromodulators of the learning and memory system of the octopus VL remained elusive. This study, therefore, aimed to gain a better understanding of the VL system, mainly motivated by the notion that neurons should be classified into cell types not only by location and shape but also according to their neurotransmitter, neuropeptides, and neuromodulators. Some of our results fit the physiological models (e.g., glutamatergic MSFL neurons, cholinergic AMs, GABAergic LNs). At the same time, other findings are novel (e.g., putative GABAergic AM) and suggest that the apparently simple input-output relationship of the VL is mediated by a more complex network than the fan-out fan-in feedforward excitatory connections found previously (Shomrat et al. 2011).

### MSF fiber inputs to the VL via L-glutamate synapses

ISH revealed VGLUT-encoding mRNA in the cell body cortex of the MSFL (**Fig 2A**), supporting L-glutamate (L-Glu) as the transmitter for these neurons. Immunolabeling of L-Glu (**Fig 4**) revealed especially densely labeled varicosities, such as those localized at the lower margin of the MSFL tract where the MSFL terminals synapse with the AM neurites (Gray 1970). These results support the physiological findings of glutamatergic connectivity at the first VL input fan-out synaptic layer (Hochner et al., 2003; Shomrat, et al., 2011). No VGLUT labeling was seen in the VL cell cortex, suggesting that the majority of the cells in the VL (e.g., AMs) use different neurotransmitters from the MSFL.

### AMs input to LNs via both cholinergic and GABAergic synapses

Specific cChAT labeling was seen in the AM neurites, with especially strong immunoreactivity in the inner neuropil, the site of the serial synapses of amacrine cells with LNs (Gray, 1970; Shomrat et al. 2011). The cholinergic processes were restricted to the inner neuropil region, emphasizing an evenly distributed cholinergic innervation in the dendritic area of the input to the LNs **(Figs 6,7)**. These results confirm the physiological findings of cholinergic AM→LN synapses (Shomrat et al. 2011).

Synaptic plasticity in this learning network occurs at predicted glutamatergic connections between the MSFL axon terminals and the AMs. Yet, in the related cuttlefish (*Sepia*), the plasticity occurs in the second layer, where ACh is the likely transmitter (Shomrat et al., 2011). This dichotomy suggests that, in cephalopods, the molecular mechanism of LTP is not associated with a single specific neurotransmitter.

Morphologically the AMs appear to make up a largely homogeneous population (Young 1971; Gray 1970). Yet, while some AM are cholinergic, we found that some AMs could be GABAergic **(Figs 13, 14)**. The proposed transmitter diversity of AMs may explain previous findings that stimulation of the MSF tract evoked IPSPs in some LNs (Shomrat et al., 2011). Thus, the AMs appear to be functionally and chemically heterogeneous. Indeed, an ongoing connectome study (Bidel et al., 2019) has been able to divide the AMs into two distinct groups. More than 95% are “simple” AMs, receiving synaptic input from MSFL neurons and sending a single non-bifurcating neurite into the neuropil. The remaining AMs are “complex”, seeming to integrate inputs from several MSF neurons and several simple AMs, their neurites bifurcating extensively in the outer neuropil at the level of the SFL tract where they receive inputs from the SFL axonal varicosities. These processes run into the inner neuropil where they innervate LNs processes. It is reasonable to propose that some interneurons are inhibitory and thus stained positively for GABA. It remains to be clarified whether this population of inhibitory AMs is the group of complex AMs discovered in the EM study or whether some other low molecular weight transmitters or small peptides are involved.

The restriction of the GABAergic varicosities to the inner neuropil, mainly in the medial and medial-lateral lobuli (**Figs 12**), indicates that dendritic branches may compartmentalize GABAergic signaling in general or, particularly, inhibition. If these GABAergic varicosities originate from GABAergic AMs, the proposed GABA-mediated inhibition may provide a feedforward inhibition of LNs. An immunohistochemical study in *Octopus bimaculoides* suggested that serotonin is distributed unevenly in the five lobuli indicating functional differentiation among them (Shigeno and Ragsdale 2015). In our study, the uneven distribution of GABAergic varicosities in the five lobuli was the most indicative result supporting further functional differentiation among the VL regions and their neurons.

### Heterogeneity and co-transmission of the LNs

Strikingly organized GABA-IR LN cell bodies were localized in the inner cortex of the VL (**Figs 11**), confirming that these neurons can provide the predicted inhibitory output from the VL as previously postulated from staining (Cornwell et al. 1993), lesioning (Boycott and Young 1955) and behavioral and physiological experiments (Shomrat et al. 2008). This provides support for the VL model in which the output has an inhibitory control over other circuits, such as those for attack behavior (see Turchetti-Maia et al. 2017).

Yet, glutamatergic LNs were also detected in the cell cortex (**Fig 4**), suggesting that like the AMs, the LNs do not comprise a homogenous population. Even taking into account that glutamatergic-IR may also label GABAergic cells, as glutamate is a metabolic precursor for the synthesis of GABA (Villar-Cerviño et al. 2013), the distribution and morphological characteristics of the group of glutamate-labeled LN cell bodies seemed to differ from the classic GABAergic LNs.

It is not clear if the axons of these LNs project out of the VL to form a parallel excitatory output that may facilitate behaviors like the attack behavior. Or they could be part of recurrent excitatory connections between the VL and the MSFL forming reverbatory cyclic networks as postulated by Young (1991, 1995). Recurrent reverbatory circuits may subserve working memory by maintaining ongoing electrical activity. Tracing techniques have revealed such possible connections in cuttlefish, though the nature of their transmission system is not yet clear (Graindorge et al. 2008).

LNs, and possibly other large cells in similar areas also express neuromodulatory neuropeptides. A FMRFamide-like neuropeptide, FLRIamide, showed the most prominent labeling pattern in specific VL neurons (**Fig 21A**). This suggests that, in addition to the fast transmitter GABA, the LNs contain a small neuropeptide FaRP as a transmitter or co-transmitter. Members of a class of neuropeptides ending in RFamide are known to be depressive/inhibitory neuromodulators in mollusks (Zhang et al. 2012; Baux et al. 1990; Cottrell 1993; van Golen et al. 1995), activating PKC and regulating cholinergic synapses (Baux, et al., 1990), and they appear similar in other cephalopods (Chrachri & Williamson 2003). Thus, this neuromodulator may be involved in long-term modulation – likely protein synthesis-dependent – in regions outside the VL, where long-term memories are stored. Similarly, FMRFamide has long-term inhibitory effects on the *Aplysia* sensory-motor synapse, a classical model for synaptic processes involved in learning and memory. However, the specific roles of Rfamide-related peptides still need to be investigated in detail.

### Neuromodulation systems in the VL - localization of putative NOS activity

NADPH-diaphorase is a reliable reporter of NOS activity in molluscs (Moroz 2000; Moroz et al. 2005; Moroz et al. 1999; Cruz et al. 1997; Floyd et al. 1998). The NADPH-diaphorase method produced intense labeling in the AM neurites and VL neuropil, indicating NOS activity in these structures (**Fig 17**). NO is a well-known anterograde neurotransmitter in sensory and motor circuits of mollusks (Moroz and Gillette 1995; Moroz and Kohn 2011; Moroz et al. 1993b; Hatcher et al. 2006; Moroz et al. 2000; Bodnarova et al. 2005; Moroz 2006). NO also mediates synaptic plasticity in mollusks, including its own release from interneurons (Antonov et al. 2007; Katzoff et al. 2002; Kemenes et al. 2002; Korshunova and Balaban 2014) and is involved in the LTP in octopus VL (Turchetti-Maia et al. 2018). The localization of NOS here fits with physiological results that suggested the retrograde mediation of LTP by NO increasing the probability of glutamate release from the presynaptic terminals of the MSFL neurons. This accords with the view of a retrograde message inducing presynaptic expression of plasticity, a commonly postulated scheme for NO-mediated plasticity, including associative (Hebbian) learning in mammals (Arancio et al. 1996; Garthwaite 2008; Prast and Philippu 2001; Turchetti-Maia et al. 2017). The similar pattern of expression of NOS and cChAT in the VL and subFL neuropils supports the presence of NOS in the cholinergic AMs. As the AMs are the postsynaptic targets of the MSFL synapses, this supports NO as a retrograde messenger in the presynaptic expression of LTP (Turchetti-Maia et al., 2018).

We could not demonstrate NOS activity in the neuropil of cuttlefish *Sepia officinalis* VL (not shown), suggesting that the molecular mechanism mediating LTP in cephalopods evolved independently in these phylogenetically close species as did the site of LTP (Shomrat et al. 2011).

### Tyrosine hydroxylase marker suggests catecholaminergic reward signaling in the VL system

Serotonin and octopamine have short-term facilitatory effects in the VL, reinforcing and suppressing LTP induction, respectively (Shomrat et al. 2010; Turchetti-Maia et al. 2017). TH, the enzyme which catalyzes the conversion of tyrosine into L-DOPA, the precursor of dopamine, was widely distributed in the MSFL and VL neuropil, indicating the involvement of catecholamines, particularly dopamine, in the MSF-VL neural network **(Figs 18, 19, 20)**. These results confirm those of (Tansey 1980), who reported scattered dopamine and/or other catecholamines in the MSFL and certain lobules in the VL neuropil, but not in the cell body layers. Here, a meshwork of thin TH-IR processes was seen in the outer neuropil in the region of the MSFL-AM synaptic connections (**Fig. 20**). This agrees with findings that dopamine could mediate a short-term facilitatory effect on the synaptic input to the AMs, while blocking the development of activity-dependent LTP (Weber I. 2018 MSc Thesis, Hebrew University).

TH staining was distributed in a stereotypical pattern in the inner VL neuropil, suggesting a specific interaction with the proximal dendrites of the LNs (**Fig. 20B**). There was a strong resemblance between the TH and the GABA labeling in the area into which the LN dendrites project in the inner VL neuropil (see **Fig. 12**). Catecholaminergic modulation and GABAergic innervation may thus occur at the same location, such as particular regions of the LN dendrites. Co-transmission, similar innervation, and synaptic locations of GABA and a catecholamine (probably dopamine) have been found in learning systems in mollusks (Díaz-Ríos et al. 2002) and mammals (Maher and Westbrook 2008).

The TH staining contrasts with the NADPH-d labeling and cChAT-IR, which showed homogenous widespread distributions throughout the entire inner VL neuropil. A finer spreading of TH-labeled process into the cell cortex in the region of the synaptic connection between the MSFL terminals and the AMs (**Fig. 19B’, 20A**) accords with the physiological finding of dopamine-dependent modulation of short- and long-term synaptic plasticity of these synaptic connections (Weber I. 2018 MSc Thesis, Hebrew University).

The abundantly labeled TH-IR cell bodies in the ‘deep nucleus’ at the MSFL-VL border (**Fig. 18**), may give rise to the TH-positive fibers running antero-posteriorly in the inner VL neuropil. This would imply that 5-HT inputs convey modulatory signals from other brain structures into the VL (Shomrat et al. 2010), like the global dopamine, noradrenaline, and acetylcholine fibers innervating the hippocampus (Matsuda et al., 2006) and modulatory inputs to the insect mushroom bodies (Fiala 2007). But, in contrast, in the octopus VL at least part of the TH modulation uniquely originates from within the MSFL-VL system itself.

Intense TH-IR neuronal processes crossing through the VL lobule hila (**Fig 20B**) showed broad interactions between the VL and surrounding lobes that may be related to the consolidation of long-term memory reinforced through LTP induction in the VL (Shomrat et al. 2010; Turchetti-Maia et al. 2017).

### Neuropeptides in the learning system

Several neuropeptide mRNAs were revealed in the MSFL and VL (**Figs. 21**-**24**); buccalin, bradykinin, conopressin, and myomodulin showed expression mainly in the cell bodies within different areas. Such differential expression of neuropeptides may play an important role in setting the specific neurophysiological properties of the lobes and beyond. The functional roles of these and other neuropeptides deserve careful attention and separate exploration.

### Striking similarities between visual and tactile learning structures

The MSFL-VL, the visual learning system, and the MIFL-subFL, the tactile learning system, both show a fan-out fan-in network organization (Sanders 1975; Young 1971). The distribution of the other neuropeptides and NADPH-d staining in the MSFL-VL system (**Fig 17**) was strikingly similar to that in the MIFL-subFL system. Similar cChAT labeling (**Fig. 9**) and GABA immunohistochemistry patterns (**Fig 15**) were also found in the two lobes. The similar structure of the MIFL-subFL assisted us in interpreting the labeling in the VL, especially because the small cell bodies of the VL AMs, almost devoid of cytoplasm, seem not to express detectable amounts of synaptic proteins. Thus, the NADPH-d reactivity of the small cell bodies in the cell layer of the subFL supported the analysis of which interneurons in the VL probably also contained NOS activity (see Fig 14A); this approach gave a fit with our interpretations of AM neurite labeling in the VL. Analyzing the separate visual and tactile learning systems, with their similar organization, in terms of cell types, neurotransmitters, neuromodulators and physiology (Shomrat, personal communication), could provide a better understanding of the overall functional organization of biological learning and memory systems controlling specific behaviors.

## Conclusion

Our results confirm previous anatomical and physiological findings of a feedforward fan-out fan-in network in the VL (Shomrat et al., 2011). The input *en passant* MSF→AM synapses are glutamatergic synapses that undergo LTP. A large group of AMs are cholinergic and mediate the fan-in excitatory input to the LNs.

Yet, immunohistochemical labeling revealed that the VL feedforward connectivity is not exclusively simpler and excitatory. Our findings suggest that MSFL glutamatergic inputs also innervate a GABAergic group of AMs, which likely feed modulatory or inhibitory inputs to the LNs. As mentioned above, an ongoing connectome study has revealed that a small proportion of the AMs have a complex bifurcating neurite tree. These “complex” AM, interneurons integrate synaptic inputs from the MSFL and “simple” AMs. This connectivity scheme makes these AMs plausible candidates for the inhibitory AMs. Furthermore, in contrast to previous assumptions, not all LNs are inhibitory; some neurons with large cell bodies were positively labeled for glutamate (and possibly ACh). Thus, the VL control of behavior seems to be intricately orchestrated by both excitatory and inhibitory pathways, providing more elaborated association modes for memory acquisition. Further physiological, anatomical and behavioral studies are needed to fully understand these pathways.

This study also contributes important insights into neuromodulatory systems, showing distributed innervation of TH-positive process in the VL, which suggest an elaborated catecholinergic neuromodulatory system. Physiological experiments have revealed the importance of serotonergic, dopaminergic, and octopaminergic modulatory reward and punishment signals in enhancing or suppressing LTP (reviewed in Turchetti-Maia et al., 2017).

Finally, in view of the proposed involvement of NO in the octopus LTP, the finding of robust expression of NADPH-d/NOS activity in the VL neuropil supports NO as a retrograde signal from the postsynaptic AMs to the presynaptic MSFL terminal leading to an increase in presynaptic glutamate release (Turchetti-Maia et al. 2018).

## Acknowledgments

This work was supported by: United States - Israel Binational Science Foundation (BSF) Grants Number: 2007-407; 2011-466. Israel Sciences Foundation (ISF) Grants Number: 1425-2011; 1928- 2015. We wish to thank Prof. Jenny Kien for editing the manuscript.

## References

1. Albertin CB, Bonnaud L, Brown CT, Crookes-Goodson WJ, da Fonseca RR, Di Cristo C, Dilkes BP, Edsinger-Gonzales E, Freeman RM, Jr., Hanlon RT, Koenig KM, Lindgren AR, Martindale MQ, Minx P, Moroz LL, Nodl MT, Nyholm SV, Ogura A, Pungor JR, Rosenthal JJ, Schwarz EM, Shigeno S, Strugnell JM, Wollesen T, Zhang G, Ragsdale CW (2012) Cephalopod genomics: A plan of strategies and organization. Stand Genomic Sci 7 (1):175–188. doi:10.4056/sigs.3136559

2. Alves C, Boal JG, Dickel L (2008) Short-distance navigation in cephalopods: a review and synthesis. Cognitive Processing 9 (4):239–247

3. Amodio P, Fiorito G (2013) Observational and other types of learning in Octopus. In: Handbook of Behavioral Neuroscience, vol 22. Elsevier, pp 293–302

4. Antonov I, Ha T, Antonova I, Moroz LL, Hawkins RD (2007) Role of Nitric Oxide in Classical Conditioning of Siphon Withdrawal in Aplysia. The Journal of Neuroscience 27 (41):10993–11002. doi:10.1523/jneurosci.2357-07.2007

5. Arancio O, Kiebler M, Lee CJ, Lev-Ram V, Tsien RY, Kandel ER, Hawkins RD (1996) Nitric Oxide Acts Directly in the Presynaptic Neuron to Produce Long-Term Potentiationin Cultured Hippocampal Neurons. Cell 87 (6):1025–1035. doi:http://dx.doi.org/10.1016/S0092-8674(00)81797-3

6. Baux G, Fossier P, Tauc L (1990) Histamine and FLRFamide regulate acetylcholine release at an identified synapse in Aplysia in opposite ways. The Journal of Physiology 429 (1):147–168

7. Boal JG (1996) A review of simultaneous visual discrimination as a method of training octopuses. BiolRev 71 157–190

8. Bodnarova M, Martasek P, Moroz LL (2005) Calcium/calmodulin-dependent nitric oxide synthase activity in the CNS of Aplysia californica: biochemical characterization and link to cGMP pathways. J Inorg Biochem 99 (4):922–928. doi:10.1016/j.jinorgbio.2005.01.012

9. Boycott BB (1954) Learning in Octopus vulgaris and other cephalopods. . Pubbl Staz Zool, Napoli 25:67–93

10. Boycott BB (1961) The functional organization of the brain of the cuttlefish Sepia officinalis Proceedings of the Royal Society of London B Biological Sciences 153 503–534

11. Boycott BB, Young JZ (1955) A memory system in Octopus vulgaris Lamarck. Proc R Soc Lond B Biol Sci 143 (913):449–480

12. Boycott BB, Young JZ (1958) Reversal of learned responses in Octopus vulgaris Lamarck. Animal Behaviour 6:45–52

13. Brusca & Brusca (1990). Invertebrates, First Edition (Sunderland: Sinauer Associates).

14. Bullock TH, Horridge GA (1965) Structure and function in the nervous systems of invertebrates. A Series of books in biology. W. H. Freeman, San Francisco,

15. Casini A, Vaccaro R, D’Este L, Sakaue Y, Bellier JP, Kimura H, Renda TG (2012) Immunolocalization of choline acetyltransferase of common type in the central brain mass of Octopus vulgaris. European Journal of Histochemistry 56 (3):215–222. doi:10.4081/ejh.2012.e34

16. Cornwell CJ, Messenger JB, Williamson R (1993) Distribution of GABA-like immunoreactivity in the octopus brain. Brain Research 621 (2):353

17. Cottrell G (1993) The wide range of actions of the FMRFamide-related peptides and the biological importance of peptidergic messengers. Comparative molecular neurobiology:279–285

18. Cruz L, Moroz LL, Gillette R, Sweedler JV (1997) Nitrite and nitrate levels in individual molluscan neurons: single-cell capillary electrophoresis analysis. J Neurochem 69 (1):110–115. doi:10.1046/j.1471-4159.1997.69010110.x

19. D’Este L, Kimura S, Casini A, Matsuo A, Bellier JP, Kimura H, Renda TG (2008) First visualization of cholinergic cells and fibers by immunohistochemistry for choline acetyltransferase of the common type in the optic lobe and peduncle complex of Octopus vulgaris. Journal Of Comparative Neurology 509 (6):566–579

20. Di Cosmo A, Di Cristo C, Messenger JB (2006) L-glutamate and its ionotropic receptors in the nervous system of cephalopods. Curr Neuropharmacol 4 (4):305–312. doi:10.2174/157015906778520809

21. Di Cosmo A, Nardi G, Di Cristo C, De Santis A, Messenger JB (1999) Localization of L-glutamate and glutamate-like recaptors at the squid giant synapse. Brain Res 839:213–220

22. Di Cosmo A, Paolucci M, Di Cristo C (2004) N-methyl-D-aspartate receptor-like immunoreactivity in the brain of Sepia and Octopus. J Comp Neurol 477 (2):202–219

23. Díaz-Ríos M, Oyola E, Miller MW (2002) Colocalization of γ-aminobutyric acid-like immunoreactivity and catecholamines in the feeding network of Aplysia californica. Journal of Comparative Neurology 445 (1):29–46

24. Fiala A (2007) Olfaction and olfactory learning in Drosophila: recent progress. Current Opinion in Neurobiology 17 (6):720

25. Fiorito G, Affuso A, Anderson DB, Basil J, Bonnaud L, Botta G, Cole A, D’Angelo L, De Girolamo P, Dennison N (2014) Cephalopods in neuroscience: regulations, research and the 3Rs. Invertebrate Neuroscience 14 (1):13–36

26. Fiorito G, Affuso A, Basil J, Cole A, de Girolamo P, D’angelo L, Dickel L, Gestal C, Grasso F, Kuba M (2015) Guidelines for the care and welfare of cephalopods in research–a consensus based on an initiative by CephRes, FELASA and the Boyd Group. Laboratory animals 49 (2_suppl):1–90

27. Fiorito G, Chichery R (1995) Lesions Of The Vertical Lobe Impair Visual-Discrimination Learning By Observation In Octopus-Vulgaris. Neuroscience Letters 192 (2):117–120

28. Fiorito G, Scotto P (1992) Observational-Learning in Octopus-Vulgaris. Science 256 (5056):545–547

29. Fiorito G, von Planta C, Scotto P (1990) Problem solving ability of Octopus vulgaris Lamarck (Mollusca, Cephalopoda). Behav Neural Biol 53 (2):217–230

30. Floyd PD, Moroz LL, Gillette R, Sweedler JV (1998) Capillary electrophoresis analysis of nitric oxide synthase related metabolites in single identified neurons. Anal Chem 70 (11):2243–2247. doi:10.1021/ac9713013

31. Garthwaite J (2008) Concepts of neural nitric oxide-mediated transmission. European Journal of Neuroscience 27 (11):2783–2802. doi:10.1111/j.1460-9568.2008.06285.x

32. Graindorge N, Alves C, Darmaillacq AS, Chichery R, Dickel L, Bellanger C (2006) Effects of dorsal and ventral vertical lobe electrolytic lesions on spatial learning and locomotor activity in Sepia officinalis. Behav Neurosci 120 (5):1151–1158

33. Graindorge N, Jozet-Alves C, Chichery R, Dickel L, Bellanger C (2008) Does kainic acid induce partial brain lesion in an invertebrate model: Sepia officinalis? Comparison with electrolytic lesion. Brain Research 1238:44

34. Gray EG (1970) The fine structure of the vertical lobe of octopus brain Phil Trans R Soc Lond B 258:379–394

35. Gutnick T, Byrne RA, Hochner B, Kuba M (2011) Octopus vulgaris uses visual information to determine the location of its arm. Curr Biol 21 (6):460–462. doi:S0960-9822(11)00108-4 [pii]

36. Gutnick T, Zullo L, Hochner B, Kuba MJ (2020) Use of Peripheral Sensory Information for Central Nervous Control of Arm Movement by Octopus vulgaris. Current Biology 30 (21):4322–4327.e4323. doi:10.1016/j.cub.2020.08.037

37. Hatcher NG, Sudlow LC, Moroz LL, Gillette R (2006) Nitric oxide potentiates cAMP-gated cation current in feeding neurons of Pleurobranchaea californica independent of cAMP and cGMP signaling pathways. J Neurophysiol 95 (5):3219–3227. doi:10.1152/jn.00815.2005

38. Heisenberg M (2003) Mushroom body memoir: from maps to models. Nat Rev Neurosci 4 (4):266

39. Hochner B, Brown ER, Langella M, Shomrat T, Fiorito G (2003) A learning and memory area in the octopus brain manifests a vertebrate-like long-term potentiation. J Neurophysiol 90 (5):3547–3554

40. Hochner B, Shomrat T, Fiorito G (2006) The octopus: a model for a comparative analysis of the evolution of learning and memory mechanisms. Biol Bull 210. doi:10.2307/4134567

41. Hope B, Vincent S (1989) Histochemical characterization of neuronal NADPH-diaphorase. Journal of Histochemistry & Cytochemistry 37 (5):653–661

42. Jezzini SH, Bodnarova M, Moroz LL (2005) Two-color in situ hybridization in the CNS of Aplysia californica. J Neurosci Methods 149 (1):15–25. doi:10.1016/j.jneumeth.2005.05.007

43. Kandel E, Schwartz J, Jessell T, Siegelbaum S, Hudspeth AJ (2012) Principles of Neural Science, Fifth Edition. McGraw-Hill Education,

44. Katzoff A, Ben-Gedalya T, Susswein AJ (2002) Nitric Oxide Is Necessary for Multiple Memory Processes after Learning That a Food Is Inedible in Aplysia. J Neurosci 22 (21):9581–9594

45. Kemenes I, Kemenes G, Andrew RJ, Benjamin PR, O’Shea M (2002) Critical Time-Window for NO–cGMP-Dependent Long-Term Memory Formation after One-Trial Appetitive Conditioning. The Journal of Neuroscience 22 (4):1414–1425

46. Korshunova T, Balaban P (2014) Nitric oxide is necessary for long-term facilitation of synaptic responses and for development of context memory in terrestrial snails. Neuroscience 266:127–135

47. Livneh Y, Feinstein N, Klein M, Mizrahi A (2009) Sensory Input Enhances Synaptogenesis of Adult-Born Neurons. Journal Of Neuroscience 29 (1):86–97

48. Mackintosh NJ (1965) Discrimination learning in the octopus. Animal Behaviour Suppl. 1 129–134

49. Maher BJ, Westbrook GL (2008) Co-transmission of dopamine and GABA in periglomerular cells. Journal of neurophysiology 99 (3):1559–1564

50. Maldonado H (1963) The visual attack learning system in Octopus vulgaris. Journal of Theoretical Biology 5 (3):470–488

51. Maldonado H (1965) The positive and negative learning process in Octopus vulgaris Lamarck. Influence of the vertical and median superior frontal lobes. Zeitschrift fur vergleichende Physiologie 51 185–203

52. Messenger JB (1996) Neurotransmitters of cephalopods. Invertebrate neuroscience (2):95–114

53. Moriyama T, Gunji YP (1997) Autonomous learning in maze solution by Octopus. Ethology 103 (6):499–513

54. Moroz LL (2000) Giant identified NO-releasing neurons and comparative histochemistry of putative nitrergic systems in gastropod molluscs. Microsc Res Tech 49 (6):557–569. doi:10.1002/1097-0029(20000615)49:6<557::AID-JEMT6>3.0.CO;2-S

55. Moroz LL (2006) Localization of putative nitrergic neurons in peripheral chemosensory areas and the central nervous system of Aplysia californica. J Comp Neurol 495 (1):10–20. doi:10.1002/cne.20842

56. Moroz LL, Dahlgren RL, Boudko D, Sweedler JV, Lovell P (2005) Direct single cell determination of nitric oxide synthase related metabolites in identified nitrergic neurons. J Inorg Biochem 99 (4):929–939. doi:10.1016/j.jinorgbio.2005.01.013

57. Moroz LL, Gillette R (1995) From Polyplacophora to Cephalopoda: comparative analysis of nitric oxide signalling in mollusca. Acta Biol Hung 46 (2-4):169–182

58. Moroz LL, Gillette R, Sweedler JV (1999) Single-cell analyses of nitrergic neurons in simple nervous systems. J Exp Biol 202 (Pt 4):333–341

59. Moroz LL, Gyori J, Salanki J (1993a) NMDA-like receptors in the CNS of molluscs. Neuroreport 4 (2):201–204. doi:10.1097/00001756-199302000-00022

60. Moroz LL, Kohn AB (2010) Do different neurons age differently? Direct genome-wide analysis of aging in single identified cholinergic neurons. Front Aging Neurosci 2. doi:10.3389/neuro.24.006.2010

61. Moroz LL, Kohn AB (2011) Parallel evolution of nitric oxide signaling: diversity of synthesis and memory pathways. Front Biosci (Landmark Ed) 16:2008–2051. doi:10.2741/3837

62. Moroz LL, Nikitin MA, Poličar PG, Kohn AB, Romanova DY (2021) Evolution of glutamatergic signaling and synapses. Neuropharmacology:in press

63. Moroz LL, Norekian TP, Pirtle TJ, Robertson KJ, Satterlie RA (2000) Distribution of NADPH-diaphorase reactivity and effects of nitric oxide on feeding and locomotory circuitry in the pteropod mollusc, Clione limacina. J Comp Neurol 427 (2):274–284

64. Moroz LL, Park JH, Winlow W (1993b) Nitric oxide activates buccal motor patterns in Lymnaea stagnalis. Neuroreport 4 (6):643–646. doi:10.1097/00001756-199306000-00010

65. Nesher N, Levy G, Zullo L, Hochner B (2020) Octopus Motor Control. Oxford University Press. doi:10.1093/acrefore/9780190264086.013.283

66. Nixon M, Young JZ (2003) The Brains and Lives of Cephalopods. Oxford University Press, Oxford

67. Papini MR, Bitterman ME (1991) Appetitive conditioning in Octopus cyanea. J Comp Psychol 105 (2):107–114

68. Prast H, Philippu A (2001) Nitric oxide as modulator of neuronal function. Progress in Neurobiology 64 (1):51–68. doi:http://dx.doi.org/10.1016/S0301-0082(00)00044-7

69. Puthanveettil SV, Antonov I, Kalachikov S, Rajasethupathy P, Choi YB, Kohn AB, Citarella M, Yu F, Karl KA, Kinet M, Morozova I, Russo JJ, Ju J, Moroz LL, Kandel ER (2013) A strategy to capture and characterize the synaptic transcriptome. Proc Natl Acad Sci U S A 110 (18):7464–7469. doi:10.1073/pnas.1304422110

70. Richter JN, Hochner B, Kuba MJ (2015) Octopus arm movements under constrained conditions: adaptation, modification and plasticity of motor primitives. The Journal of Experimental Biology 218 (7):1069–1076. doi:10.1242/jeb.115915

71. Richter JN, Hochner B, Kuba MJ (2016) Pull or Push? Octopuses Solve a Puzzle Problem. PloS one 11 (3):e0152048

72. Rosenblatt F (1958) The perceptron: A probabilistic model for information storage and organization in the brain. Psychological Review 65 (6):386–408. doi:10.1037/h0042519

73. Sakaue Y, Bellier J-P, Kimura S, D’Este L, Takeuchi Y, Kimura H (2014) Immunohistochemical localization of two types of choline acetyltransferase in neurons and sensory cells of the octopus arm. Brain Struct Funct 219 (1):323–341. doi:10.1007/s00429-012-0502-6

74. Sanders GD (1975) The Cephalopods. In: Corning WC, Dyal JA, Willows AOD (eds) Invertebrate learning,, vol Vol:3: Cephalopods and Echinoderms Plenum Press, New York, pp 139–145

75. Shigeno S, Ragsdale CW (2015) The gyri of the octopus vertical lobe have distinct neurochemical identities. J Comp Neurol 523 (9):1297–1317. doi:10.1002/cne.23755

76. Shomrat T, Feinstein N, Klein M, Hochner B (2010) Serotonin is a facilitatory neuromodulator of synaptic transmission and “reinforces” long-term potentiation induction in the vertical lobe of Octopus vulgaris. Neuroscience 169 (1):52–64

77. Shomrat T, Graindorge N, Bellanger C, Fiorito G, Loewenstein Y, Hochner B (2011) Alternative Sites of Synaptic Plasticity in Two Homologous Fan-out Fan-in Learning and Memory Networks. Current biology : CB 21 (21):1773–1782

78. Shomrat T, Turchetti-Maia AL, Stern-Mentch N, Basil JA, Hochner B (2015) The vertical lobe of cephalopods: an attractive brain structure for understanding the evolution of advanced learning and memory systems. J Comp Physiol A 201 (9):947–956. doi:10.1007/s00359-015-1023-6

79. Shomrat T, Zarrella I, Fiorito G, Hochner B (2008) The octopus vertical lobe modulates short-term learning rate and uses LTP to acquire long-term memory. Current Biology 18 (5):337–342. doi:10.1016/j.cub.2008.01.056

80. Striedter GF, Belgard TG, Chen CC, Davis FP, Finlay BL, Gunturkun O, Hale ME, Harris JA, Hecht EE, Hof PR, Hofmann HA, Holland LZ, Iwaniuk AN, Jarvis ED, Karten HJ, Katz PS, Kristan WB, Macagno ER, Mitra PP, Moroz LL, Preuss TM, Ragsdale CW, Sherwood CC, Stevens CF, Stuttgen MC, Tsumoto T, Wilczynski W (2014) NSF workshop report: discovering general principles of nervous system organization by comparing brain maps across species. J Comp Neurol 522 (7):1445–1453. doi:10.1002/cne.23568

81. Sutherland NS (1959) Visual discrimination of shape by octopus. Circles and squares, and circles and triangles. Q J Exp Psychol 11 24–32

82. Taghert PH, Nitabach MN (2012) Peptide neuromodulation in invertebrate model systems. Neuron 76 (1):82–97

83. Tansey EM (1979) Neurotransmitters in the cephalopod brain. Comp Biochem Physiol 64C:173–182

84. Tansey EM (1980) Aminergic Fluorescence In The Cephalopod Brain. Philosophical Transactions Of The Royal Society Of London Series B-Biological Sciences 291 (1046):127–&

85. Turchetti-Maia A, Shomrat T, Hochner B (2017) The vertical lobe of cephalopods—a brain structure ideal for exploring the mechanisms of complex forms of learning and memory. The Oxford handbook of invertebrate neurobiology Oxford University Press, Oxford:1–27

86. Turchetti-Maia AL, Stern-Mentch N, Bidel F, Nesher N, Shomrat T, Hochner B (2018) A novel NO-dependent ‘molecular-memory-switch’ mediates presynaptic expression and postsynaptic maintenance of LTP in the octopus brain. bioRxiv:491340. doi:10.1101/491340

87. van Golen FA, Li KW, de Lange RP, Jespersen S, Geraerts WP (1995) Mutually exclusive neuronal expression of peptides encoded by the FMRFa gene underlies a differential control of copulation in Lymnaea. Journal of Biological Chemistry 270 (47):28487–28493

88. Vapnik VN (1998) Statistical learning theory. John Wiley &Sons, Inc., New York

89. Vehovszky Á, Szabó H, Elliott CJ (2004) Octopamine-containing (OC) interneurons enhance central pattern generator activity in sucrose-induced feeding in the snail Lymnaea. J Comp Physiol A 190 (10):837–846

90. Villar-Cerviño V, Barreiro-Iglesias A, Fernández-lópez B, Mazan S, Rodicio MC, Anadón R (2013) Glutamatergic neuronal populations in the brainstem of the sea lamprey, Petromyzon marinus: an in situ hybridization and immunocytochemical study. Journal of Comparative Neurology 521 (3):522–557

91. Wells MJ (1978) Octopus. Chapman and Hall, London

92. Wentzell MM, Martínez-Rubio C, Miller MW, Murphy AD (2009) Comparative Neurobiology of Feeding in the Opisthobranch Sea Slug, Aplysia, and the Pulmonate Snail, Helisoma: Evolutionary Considerations. Brain, Behavior and Evolution 74 (3):219–230. doi:10.1159/000258668

93. Winters GC (2018) Molecular mapping of the octopus brain. PhD Dissertation, University of Florida. https://ufdc.ufl.edu/UFE0054028/00001.

94. Winters GC, Polese G, Di Cosmo A, Moroz LL (2020) Mapping of neuropeptide Y expression in Octopus brains. J Morphol 281 (7):790–801. doi:10.1002/jmor.21141

95. Wolff GH, Strausfeld NJ (2016) Genealogical correspondence of a forebrain centre implies an executive brain in the protostome–deuterostome bilaterian ancestor. Phil Trans R Soc B 371 (1685):20150055

96. Yeckel MF, Kapur A, Johnston D (1999) Multiple forms of LTP in hippocampal CA3 neurons use a common postsynaptic mechanism. Nature Neuroscience 2 (7):625–633

97. Yoshida MA, Ogura A, Ikeo K, Shigeno S, Moritaki T, Winters GC, Kohn AB, Moroz LL (2015) Molecular Evidence for Convergence and Parallelism in Evolution of Complex Brains of Cephalopod Molluscs: Insights from Visual Systems. Integr Comp Biol 55 (6):1070–1083. doi:10.1093/icb/icv049

98. Young JZ (1971) The anatomy of the nervous system of Octopus vulgaris. Clarendon Press, Oxford

99. Young JZ (1991) Computation In The Learning-System Of Cephalopods. Biological Bulletin 180 (2):200–208

100. Young JZ (1995) Multiple matrices in the memory system of octopus. In: Abbott JN, Williamson R, Maddock L (eds) Cephalopod Neurobiology. Oxford University Press, Oxford, pp 431–443

101. Zhang Z, Goodwin E, Loi PK, Tublitz NJ (2012) Molecular analysis of a novel FMRFamide-related peptide gene (SOFaRP2) and its expression pattern in the brain of the European cuttlefish *Sepia officinalis*. Peptides 34 (1):114–119. doi:http://dx.doi.org/10.1016/j.peptides.2011.07.011

