## Supplementary Data for "Neurotransmission and neuromodulation systems in the learning and memory network of *Octopus vulgaris*"

| Gene name in this Manuscript | Gene reference from Winters, 2018. | Clone Sequence | Published RNA Accession Number |
| --- | --- | --- | --- |
| FMRFamide neuropeptides-like (FLPs)* | FLRIamide | TCAAAGATCAACAACTTCAGTGCCTGGAACTCCAACAGAAGAATCTTATTAAAGAAAATGGCCGGTCTTTGGAGAATCGTTCTTTTAGCAACAATTACTTTGATGTGTGTGACATCACAATGGGTTATCTATGTCCAGGCTGAAGAAGCTTCAAGCAGCGACGAATTGGTGAACAGTGCGAATGAAGCTGAACCTGAAATGGATAGTGAAGATGATCTGAAACGAGCAAATACATTTCTTCGAATAGGTAAAGCAAACGGAATTTTACGTTTAGCTCGAAGCCCTAGTTCTTTCCTGAGAATTGGACGTCGCCCCCCATTGCACTTTGTTAGAATCGGTAAAGCTCCATCAAGCATGTTTCTAAGAATTGGCAAAAGAGTTGATGATAGCGAAAATGAAGATCTAAACGACTACGATGCAGATAAAATGTCAGGGCGTGATACCAGAGCCAGTCCATCTTCGTTTCTTAGAATTGGAAAATCAGGAAATGAAGTAGAAGATGAAANNNNNNNNNNNNNNNNNNNNNTTAAACGTGTTAACGCATTTCTACGTATTGGTCGACAAAACGATCCATCTTCATTCCTGAGAATTGGTAAAAGTTTGAATAATGAAGATCTCTCTAAGGACAAACGT | Published in O. bimaculoides as: XM_014935811.1 (93.93% sequence similarity) |
| FMRFamide | FMRFamide | CCCATAATGAGTGGCTTGAGTCCCTTCAGCTTGTTAATTTTTATTTTCATCTACTGTCTCACGGTGTATGCTGCTACAGCTATGAGCCTGGCTCAGGCATGTATGGAGTCACCCAGTATGTGCGAAAGTATCTCGCTTTCTCATTTAGGTTCTGAAGACGGATTAAAGAGCAAACGCTTTCTTCGTCCAGGAAGAGCTCTTTCTGGAGATGCTTTCTTGAGATTTGGTAAAAATGGTCTAAATTTACCATTTGAAGACAAACGATTCCTTCGATTTGGTAGAACTGATCGACAATTTGAAGATATGCTAAAGGAAGTTTTACAGAGAGCTGAAACCGTTGACAATAGACAAAAGAGATCCACTGACAACATTCCTCAAAGCTCAGTTCAAGATGACAGTTCGAAAATTACGAAGAGAAATGCTGATGCCAGTGAAAACGGAATGGACAAGAGATTTATGAAATTTGGTAAATCCGGTGATCCTTACGTCAATGAATGGAGTGACAAGCGTTTCATGAGATTCGGTCGGGAACCTGACAAGCGATTCATGCGATTTGGAAAATCTGATGATAAACGATTCATGCGCTTTGGTAGAAACCCAAGTGAACTTGAAGATAGAATCGAGGAAGGAAAACGCTTTATGAGATTTGGACGGGGCAATGATGAAGAAGAAAAACGGTTTATGAGGTTTGGACGTGATCCTGACAGTAAGTTAATGGGTTATGGTAATAGTGACGAAGAAAAACGCTTTATGCGCTTTGGTAGAATGTCAGATGAAGCTGATGCTCAAAAAAGGTTCATGAGATTTGGCCGAAGTGTTGATGATGACAAGGCGAACAGGTTGAAAAAATCAAACGATCAGACCGACTGCG | [MN081862.1](https://www.ncbi.nlm.nih.gov/nucleotide/MN081862.1?report=genbank&log$=nucltop&blast_rank=2&RID=CX1BVENZ013) |
| Bradykinin | Bradykinin | ACATTCATTGCCGAAGCGGAATGACCATGTTATTACTCTGCGTTCGTTACGACAAACCGG  CCTCAGTGATTCTGACAGTCGAGCATTACTTCAAGCATACATTCTCGGAAAGCTATCCAA  CGTAGACATTGCCGGTAACAAGGAAGCAGACAATTCAGATTATACTACCATTAAGAGGAA  AGCATTTTGGAGACCTATGGGTTACCTTCCTTTTGATAACCATGGCGGTAGTGGCGCTAG  CAGCAGCAACGATAACTCTCCCAGTGGAGGCACAGGATCTGCTGTATTCAGATATGGTTA  AGCGACTCTGAAATAAAAAAAAAATCAAATATATAATGAGGAGAGAATTAATGAGTATCA  GTATGAACTAAGTAAAGACCTAACGAGTTGGTTATTGTCTTCAGCGAGGTTGATTCAATT  AAAATAGTTATTCTTC | Published in O. bimaculoides as: [XR_001409235.1](https://www.ncbi.nlm.nih.gov/nucleotide/XR_001409235.1?report=genbank&log$=nucltop&blast_rank=2&RID=CX1FKTA0013)  (97.25% sequence similarity) |
| conopressin | Conopressin/  Cephalotocin | ATATTCTGCCGTAGTCAGGCGGAAGAAGTGGTAAAGAGAAAACCAGCAAAGAAAAAAAATATCTAGATAATACTACAAGA  GATAACTATAAGTACAACCACACCACAAGAATTAAACAAGAGACCTATTTATATATTTGCATAGTATACAGCATATATCT  TTAAAATACCTTTATTAATTATATATCTATATCTGAAAAAAATTATACAGAACAACTGTGAAAATGTCTCAAAATTGTTT  CGCCATCGTCCAACTTTTGTTCGTGTTGTTCACAGTATGCAGTCTCTTCATTGCCACTACAGATGGTTGCTATTTCCGAA  ACTGTCCCATTGGTGGCAAACGAGCGACACCCATGTCAGAGCAGGGATCAAATCAAAAGTGTATGTCTTGCGGACCAAAT  GGTGAAGGCCAGTGTGTTGGATCCAACATATGCTGCCACAAAGACGGTTGCATCATAGGCACACTCGCAAAGGAATGCAA  TGAGGAAAACGAGAGCACGACAGCATGTTCTGTGAAAGGGGTGCCGTGTGGAACTGACGGACAAGGACGATGTGTTGCCG  ACGGTGTTTGCTGCGATGAATCCTCCTGTTTTACAACCGATAGATGTGACAGAGAGAATCATCGCAGTATGGCAATGCAG  AAATTATTGGAAATACGTGATGGAATTTATTACAAGAAATAAATGAAAGAATCATAAAAGGACCATTCTCAAAACGAGAT | [AB162925.1](https://www.ncbi.nlm.nih.gov/nucleotide/AB162925.1?report=genbank&log$=nucltop&blast_rank=1&RID=CX1NXSE701R) |
| Myomodulin | Myomodulin-1 | TGATAATATACTTTTCCCCCGCCAAATAACCCTCCCACGATATGGAAAAGATGAGGGATCTGATGCAGCAGATGCCGAACAATATTTGTCATCGGATTGTGAAATCTTTGATGCCTATGGCACATGTGTGCAGTATAAAATACCAAGAGAGAATGTGAAGAGGCAAGTTAGAATGTTGCGATTAGGAAAGCGTCAGATGCATCCAATAAAAATAGATGATGAAACAGGGATGGACAAAAGGTCAGAAGATTTTAACGTGGATGGAAACTCCGGTGAAAGTAAACGGGCAGTCTCAATGTTGCGTCTCGGAAGAAGTTTCAATGCTGGTGGGGAATCTGAAGAAAATGAAATTCCAGTTTTCGACGAAGCCAAACGAGCAGTTTCTATGTTACGGCTTGGCCGCAGTGGACCATTCCTTGATAAACGTGCACTTTCAATGCTCCGACTTGGTCGAAGCGAATTTGATCAGGAAAACACAGCTCTCCCAATAATGTATCCAGGACCAAATGGTTTCGAAAGCAACAAGCGTGCTGTTTCAATGTTGCGCCTCGGTCGCAGTATGTCTGCTGATGATAAAAGAGCTGTGTCTATGTTACGTCTCGGTCGCAGCGGATTTGATACAATGAAGCGAGCAGTTTCAATGTTACGCCTTGGAAGAAACAGTGGATATCCATCAGAAAAACGAGCAGTGTCCATGTTGAGATTAGGACGATCAGGATCAGATGAAGACAAGAGAGCAGTGTCTATGTTGCGGTTAGGACGATCAGGATCAGATGAAGACAAGAGAGCAGTGTCTATGTTGCGGTTAGGACGTAGTGGCGCAGATATAGAAGACGAAAAGCGAGCAGTGTCGATGTTAAGGCTTGGTCGTGGTGGTGCT  GATAATATGAAGAGAG | Published in O. bimaculoides as: [XM_014930314.1](https://www.ncbi.nlm.nih.gov/nucleotide/XM_014930314.1?report=genbank&log$=nucltop&blast_rank=6&RID=CX1WBG2W013)  (93.98% sequence similarity) |
| Sensorin | Sp191 | CATATATATATATAATTCTCGCTATACCAAGCGCTAGCAGAGACCTCTGTATGTGTGTATATATATATATTTCCGAAGCTAACTCCAGCAAAGGCATTCTGCCGATCACTGCCGAATAGTTCCGAACTCTCTACGTAAACTAACGAAAGGAACTGCAAGTCTAGCTAGATTTTTAAATTGTCATTCTTCCTCAACACAAACACCATGAAGACGTATCAAAGCCGAAACTTGTGCATTTTAAGTTGTGTTTCGTTGATCCTTGTTCTTTGCTGTGGAGTCGACGCCTCGAAGTCTCTAACTAAACGACTAGGAAATGGTTTCAAAAACAACATACTCGACCGTATGGCCCATGGCTTTGGGAAAAGAACACTTGAAGATACATACGACCTGGCACCATCAATAGCCGAAACAGGAACTAGAAATTGGATGACCCTCAAAGACTTGGCTGACCTTATGACCTACGACAGTGATTTGGCTTACAGAGCGGCACTTAAATTGGACACTAACGGTGATGGAGTGATTTCAATGGACGAGTTTGTAGGAGGTTTGCGTGGACGTGGAACTGCCTAAATTCCAGAAACCTATAGAATCTGCTACCCGCCTTCTTTTCCCAATACACATACATCCATACATAGGCTCACACACACTCGCGGATATGTATTACACACGTATATACACATGCATAATATATATGCATACGTGCATATATGTAAATATGTATGTAGGCATGTGTGCGTATATCGTATATAGT | [GQ466348.1](https://www.ncbi.nlm.nih.gov/nucleotide/GQ466348.1?report=genbank&log$=nucltop&blast_rank=1&RID=CX2C3PZX013) |
| Buccalin | Buccalin | GAAACATCCGTGTCATTGCACGTGGTATGGACCCCATGATGTTCGGTAATCTAGGAAAACGAATGGACCCCAACATGTTTGGATCATTGGGAAAACGATTCAACAGGGACGATAGAGAAAAGCAAGATAAGAAAATAGACCCATATATGTTTGGTAGCCTTGGAAAGAGAATGGATCCTGCAATGTTCGGATCTCTTGGGAAACGAATGGATCCAATGTTGTATGGTGGTTTAGGAAAACGCATGAGCCCCGACTACATCAACTATGGCGGTAAACGATATGATCCAATGTTATTCGGAGGTTTAGGGAAAAGAGTGGACCCCATGTTNTTCGGTGGTTTAGGGAAAAAAGATGGACCCCCAATATGTTTGGAACCCTAGGAAAACGTATGGACTCTATGATGTTCGGTCACCTCGGGAAAAGAGATGTATCTGGAATCAAAGTAGGTGACTTTGAATAAATATACACGTTCTAGGGAACCCATCGTTAGAAC | Published in O. bimaculoides as: [XM_014914567.1](https://www.ncbi.nlm.nih.gov/nucleotide/XM_014914567.1?report=genbank&log$=nucltop&blast_rank=2&RID=CX2K71AK013)  (96.75% sequence similarity) |
| GABA-B-like receptor mRNA | N/A | TNGGTTGGNTAACGNTCNTNAGGGCGACATGANTTAGCGGCCGCGATTCGCCCTCTAAGCTCTCGAAGGCCATCTCTTACAACAGAGGCCATGNTCTTGAACCAACAGCCTACAAAGACTTTGGCAAATATGACATCGCAAAGATATAAAGAACGTCTTGACTCCAAGCTAAAATGTCTCTGACACAAGCCAAGTCACCGGATACCCAGAATCTCCCCTTGCTTATGATGCATTGTGGGCTGTGGCATTAGCGCTAAACAAAACAGCAGCAAGATTAGCATTAAAAAATCTCACATTGGATGAATTCGATTACAATAAAAAGCATATTACAGGATGAAATTTATTCAGCAATGGATTCAACAAGTTTTCTAGGAGTGTCGGGAAATGTAGCTTTCTCTTCNGAAGGGTGACCGAATTGCTTTGACACAAATCGAACAAATGATAAATGGAACCTACCATTTAATGGGATATTATGANTCTGTTACTGATAACCTCACCTGGTTAGATAAAGAACATNGNCT | Published in O. bimaculoides as: [XM_014918957.1](https://www.ncbi.nlm.nih.gov/nucleotide/XM_014918957.1?report=genbank&log$=nucltop&blast_rank=2&RID=CX362VN4013)  (95.42% sequence similarity) |
| VGLUT | vesicular glutamate transporter | TNGTANGAAACGACTCATNAGGGCGANTGANTTAGCGGCCGCGAATTCGCCCTTCGCCATGGAAGAAATTCTTTACGTCTAAACCAGTCTACGCTATCATGGTTGCTAACTTTTGTCGTAGTTGGACATTTTATTTATTAATTATTGAACAACCGACATACTTCAAAGAAGCGTTCCGTTTTAATGTATCTCAGAGTGGCATCGTATCTGCTCTTCCTCATCTTGTTATGGCTATTATTGTACCATTTGGTGGTCAGTTGGCCGATTTTTTGAGGCGAAATGGCTACTTATCAACCACAAATGTAAGGAAAATATTTAACTGCGGAGGTTTCGGAATGGAAGCCGTATTCCTATTAGGTGTTGGCTATACTGCTGACAGAGTTACTGCAATTATATGTCTCACATTAGCAGTTGGCTTCAGTGGATTTTGCTATATCAGGNTTTAACCGTCAACCATCTTGGATATCGCANCCTCGNTACCGCTAGTATACTCATGGGNTTNGNCAC | Published in O. bimaculoides as: [XM_014920898.1](https://www.ncbi.nlm.nih.gov/nucleotide/XM_014920898.1?report=genbank&log$=nucltop&blast_rank=4&RID=CX3FWDZ3013)  (96.64% sequence similarity) |

Alignments for neuropeptides (Winters, 2018):

```
FLRI

                            *        20         *        40         *        60         *        80
OvClone196       : ----------------------------------------------------------MCVTSQWVIYVQAEEASSSDEL :  22
Octopus.b        : -------------------------------------------MAGLWRIVLLATITLMCVTSQWVIYVQAEEASSSDEL :  37
Doryteuthis.p    : -------------------------------SFQHIHNKPTRKMVGFWRILVLGTIGLVVLMTQWASFVRAESPSNGEDL :  49
Sepia.o          : -------------------------------------------MVSFWKILILGTIGVLVLMTQWASFVRAESPSNGEDL :  37
Nautilus.p       : -------------------------------------------MLSSWSVLGFAVTCIFLLYGGGLTGAGAAGASCVDDL :  37
Lottia.g         : ---------------------------------------MECDESSVRRHQPILWS--KDSLLSVIDIGQHLHMTTFDEM :  39
Biomphalaria.g   : --------------MFINIILISVTVLLHGVSGDI----AEDNLEDDKRASSFVRIGRPSSFVRIGRGDNVEDLETDPNY :  62
Charonia.t       : ------------MQMFTFLPLLSLFCPLI--------------LSACSRAA-------PT-------------------- :  27
Crassostrea.g    : MLRPHHVIIVGLFYCYTTNAEINENKLLHPIKTEEGANEILGDKADDKRSRGFFRIGKKSAVENEANDKKFDSKTVKEED :  80


                            *       100         *       120         *       140         *       160
OvClone196       : VNSA------------------------------------NEAEP----------------------------EMDSEDD :  38
Octopus.b        : VNSA------------------------------------NEADP----------------------------EMDSEDD :  53
Doryteuthis.p    : VNAAG-------------------------------AGVESADEPSGRS------------------------VSDSPYD :  74
Sepia.o          : VNAAG-------------------------------AAVESADEPSGRS------------------------VSDSPYD :  62
Nautilus.p       : CSLRK-------------------------------AGEDLAAEDGFDD------------------------LSKSPDE :  62
Lottia.g         : IQLTRQIMEINLVFS-LLLVCSISFVLSH----PFTDDSNNDDQDLKS-----------------------AIQDGDTVP :  91
Biomphalaria.g   : VDLEKKASNFVRIGRYPTMSRFIRIGRTPMEGAGSYEDDSSEEEPIGDDGKRASSFVRIGKRKSSFVRIGKSLAEEDSDE : 142
Charonia.t       : -DVTK-------------------------------DDQSDVAGPVG------------------------QFDDVSGEG :  51
Crassostrea.g    : NYIPEKIQLIRVVNSESETPIYVPVEFDP----ESSDDTADEDEKRAS-----------------------GFFRIGKSA : 133


                            *       180         *       200         *       220         *       240
OvClone196       : --LKRANTFLRIGKA-NGILRLARS-PSSFLRIGRRP----PLHFVRIGKAP---------SSMFLRIGKRV----DDSE :  97
Octopus.b        : --LKRANTFLRIGKA-NGILRLARS-PSSFLRIGRRP----PLHFVRIGKAP---------SSMFLRIGKRV----DDSE : 112
Doryteuthis.p    : --IKRANQFLRIGRG-SHFIRIGRGGASSFLRIGRNP----LSQFVRIGKAP---------SGMFLRIGKSP----AAES : 134
Sepia.o          : --IKRTNQFLRIGRG-SHFIRIGRGGASSFLRIGRNP----LSQFVRIGKAP---------SSMFLRIGKSS----AAGN : 122
Nautilus.p       : EEAKRANSFLRIGKSPSAFLRIGKGYPSTFLRIGRVP----HSSFMRIGRSP---------ASTFLRIGRGTGYDDLDSD : 129
Lottia.g         : QAVKRPSSFVRIGRNPSSFVRIGKAFGR-FIRIGKNDPNKRLSSFVRIGKSD-----PNKRISSFVRIGKSQ----EFN- : 160
Biomphalaria.g   : DVDKRASSFVRIGKSPSSFVRIGKAPSS-FVRIGKS-----PSSFVRIGKSPSSFVRIGKVPSSFVRIGRSI----ENDL : 212
Charonia.t       : DLAKRLSSFVRIGR-PNSFVRIGRG-SR-FVRIGR------PGNFVRIGRG---------YEGDLDDLGYDT----EGD- : 108
Crassostrea.g    : ENVDKRKGFFRIGKSVDQNPMNKKASG--FFRIGRTPIDKRGKGFFRIGKSLNEM--DEKRASGFFRIGKSA----LND- : 204
                      k4   F RIG4      r  4     F RIG4         F RIG4            s f r6G

                            *       260         *       280         *       300         *       320
OvClone196       : NEDLNDYDADKMSGRDTRASPSSFLRIGKSGNEVEDE--------KRVNAFLRIGRQND----------PSSFLRIGKSL : 159
Octopus.b        : NDDLNEYDADKMSGRDTRASPSSFLRIGKSGNEVEDEIDDTDETVKRVNAFLRIGRQND----------PSSFLRIGKSL : 182
Doryteuthis.p    : PVELGELAAGPSSLDDNSID--------------EDE------VIKRASSFLRIGRPN-----------PSTFLRIGKSV : 183
Sepia.o          : P-ELGDLAAGPSSLGDNSID--------------EDE------VLKRASSFLRIGRSN-----------PSTFLRIGKSA : 170
Nautilus.p       : PSELVEQEDTDPSYVRSEPSP--FFRIGGNGN--DDE--------KRAHAFLRIGKSH-----------PSTFLRIGKST : 186
Lottia.g         : -QEPE-----KRQSSFVRIGKSPELNENEIPN-------------KRYSSFVRIGKSMDDGSLENPDKRYSSFVRIGKNI : 221
Biomphalaria.g   : KSELDDIDEEKKASSFVRIGKSPNEELVDSEE-----------EKKRASSFVRIGKSGL--NDQDLFKRVSSFVRIGKSQ : 279
Charonia.t       : ----NYNNADKRASRFVRIGK--------------------------GSRFVRIGKSGQ--VDP-QIKRMSSFVRIGKAD : 155
Crassostrea.g    : -KRSRGFFRIGRSKGFFRIGKAFPLDGEKRASGFFGLAEIHLKNEEKASKFFRIGKSVN--SKEENDKRASGFFRIGKKC : 281
                                                                k    F RIG4              S F RIGK

                            *       340         *       360         *       380         *       400
OvClone196       : N------------------------NEDLSKD----------KR------------------------------------ : 169
Octopus.b        : N------------------------NDDLSKD----------KRTNAFLRIGKIPASSFIRLGRGPFTE-------DNGI : 221
Doryteuthis.p    : SGIDDETA----------------ENEEVATDEVDVPSESVAKRANAFLRIGKIPASSFVRIGRGPYGI-------DNRN : 240
Sepia.o          : GNLDEETA----------------ANEDIVTDDIDVPSESMEKRANAFLRIGKIPASSFVRIGRGPYGI-------DNRS : 227
Nautilus.p       : D-------------------------AGDSVD----------KRASAFLRIGKIPTSTFVRIGRRPFS--------SSSA : 223
Lottia.g         : ENELTNAG---------LEKRPSSFVRIGKSYFAEPGDMDAEKRLSNFVRIGKSGLE---------------EPEMEQ-- : 275
Biomphalaria.g   : GEEDKRVSSFVRIGKSGADEVEDEGKRASSFVRIGKSDTPMDKKASSFVRIGKSSTSPAETSSDSANSAISDEDPINIAS : 359
Charonia.t       : PYSDL--------------DSDDLSKRASSFVRIGR------IPSSAFVRIGRAS------SEDSGE-----------AG : 198
Crassostrea.g    : SGDSDKAGDNLT-----EDKSQSNPENEDTSESFNRNSDEPVRRASQFFRIGKSSSNKVTKRSSGVNS----SPEQNLNL : 352
                                                                  f rig

                            *       420         *       440         *       460         *
OvClone196       : ----------------------------------------------------------------------- :   -
Octopus.b        : NTRGFRGPT----RGFLRIGKRAAIPDGS-----------HADYFSDLNVKSQ------------------ : 259
Doryteuthis.p    : SPRGFLSVG----SRFVRIGKREAIPSEISQA--------HARLLSKLHDQAQ------------------ : 281
Sepia.o          : NPRGFLSVG----SRFVRIGKREAIPSETGPT--------HARLLPNLHDQAQ------------------ : 268
Nautilus.p       : RPNSFLRIGRLGTSSFVRIGKRQAVDEGENDSPLLAKGILASKVLTDGSKEQMK----------------- : 277
Lottia.g         : -KRAFVRIGKIPSSAFVRIGRMPLYDAILQKPLGYYNVARRMGK-SSFVRIGKRNNEA------------- : 331
Biomphalaria.g   : RSSAFVRIGKIPSSAFVRIGKNTNLLTAPSENWKLGFRRGSREGQSSFVRIGK------------------ : 412
Charonia.t       : DFGTFDRIARMGQSSFVRIGKRE----ADPE--KLAAQAHNLKQ--------------------------- : 236
Crassostrea.g    : NKRAFFRIGKVPTSAFMRIGRQHLLQSLVSDPLYRN---GRIQQ-SSFIRIGKRSMSDNHLIDDEQSDSSL : 419
                       f        s f rig


FMRFamide

                            *        20         *        40         *        60         *        80
OvClone197       : MSGL--SPFSLLIFIFIYCLTVYAATAMSLAQACMES---PSM-----CESISL-----SHLGSEDGLKSKRFLRPG-RA :  64
Octopus.b        : MSGL--SPFSLLIFIFIYCLTVYAATAMSLAQACMES---PSM-----CESISL-----SHLSSEEGLKSKRFLRPG-RA :  64
Doryteuthis.p    : MRCW--SPCSLLVVIVIYCLSSHTSEAFDLAQACVES---QRLSLLPICDTIFAVQQEGAQQSADDGMRSKRFIRFG-RA :  74
Nautilus.p       : MQSW--SQWSLWAVIFFKYLCT-NSLATDLVTACAEAEIRQDTSLIPLCETLMA-----VDEAPDNSVRSKRFLRFG-RA :  71
Aplysia.c        : ------MRF------------------------------------------------------------GKRFMRFG-KR :  13
Lymnaea.s        : --MY--SP------TLIVCLSFFHSAV------------------------------------------TKRFLRFG-RA :  27
Mytilus.e        : --MWTKSYATLIVAAIINWISV-KVHADELSRWCLDN---QEI-----CSNLIQ----NLRDTDDTNAQKRNFLRFG-RA :  64
Crassostrea.g    : MGTW--TYLCLLVAFLLNWFTI-ETSANDLIDDCYRN---PEL-----CQEVGI-------LFGQQQPVDKRFLRFGKRA :  62
                             l               a  l   c              c                     4rF6RfG 4a

                            *       100         *       120         *       140         *       160
OvClone197       : L-SGD---AFLRF-------GKNGLNLPFEDKRFLRFGRT-------DRQFEDMLKE-VLQRAETVDN------RQKRST : 119
Octopus.b        : L-SGD---AFLRF-------GKNGLNLPFEDKRFLRFGRT-------DRQFEDMLKE-VLQRAENVDN------RQKRST : 119
Doryteuthis.p    : L-SGD---AFLRF-------GKNVPDLPFEDKRFLRFGRA-------APQLDDLLKQ-ALQRVESLQKADETSVRRKRST : 135
Nautilus.p       : L-SGD---RFLRF-------GRSISPALFEDKRFLRFGRS-------TQSLEELLRE-ALDHMQTLEQMEGQKIRRKRST : 132
Aplysia.c        : E-DGEPDKRFMRF-------GKSMADNDL-DKRFMRFGKR-------FMRFGKSLPD------SEVDK---RFMRFGKSV :  68
Lymnaea.s        : LDTTD---PFIRLRRQFYRIGRGGYQ-PYQDKRFLRFGRS------EQPDVDDYPRDVVLQSEEPLY-------RKRRST :  90
Mytilus.e        : L-AGD---HFFRF-------GRSPYQT--EDKRFLRFGRSGGGGGFDNVGLADILKA-ALVKVESANQ-NGLKIRKRRSI : 129
Crassostrea.g    : L-SGD---HYIRF-------GRNS-----DDKRFLRFGKR-----GEQGSVEDDLRE-ALNKVIKFKQETGLHLRKRRSA : 120
                   l  gd    5 Rf       G4        DKRF6RFG4             d l    l              R  4S

                            *       180         *       200         *       220         *       240
OvClone197       : DN---IPQSSVQ-----DDSSKITKRNADAS---------ENGM--DKRFMKFGKS----GD-PYV--NEWSDKRFMRFG : 173
Octopus.b        : DN---IPQSSVQ-----DDSSKITKRNADAS---------DNGM--DKRFMKFGKS----GDLPYV--NEWSDKRFMRFG : 174
Doryteuthis.p    : DA---APQNNAENPEQKNDSAKITKRYIDDV---------EDSD--VKRFMRFGKRFMRFGRNPSDAGNKLTEKRFMRFG : 201
Nautilus.p       : DV---DPSADA---------------------------------------------------------NRLKEEKALETE : 152
Aplysia.c        : DG----------------DVDKRFMRFGKSV---------DDAV--DKRFMRFGKSV-----------DSDLDKRFMRFG : 110
Lymnaea.s        : EA---GGQSEE-----------MTHRTARSA---------PEPAAENREIMKRET-----GAEDLD-----EEKRFMRFG : 137
Mytilus.e        : DAVKDVPEKKSVT-DNTEPEAEIKKRNVDNSYATNEDSADENLKAADKRFMRFGKRFMRFGK------RENADKRFMRFG : 202
Crassostrea.g    : DP----P-------------------LVKDV---------PEDK--DSNSTEKE--------------DSASEKHKRETD : 152
                   d     p                  r                        m                      k f rfg

                            *       260         *       280         *       300         *       320
OvClone197       : R----E------------PDKRFMRFGKS---D--DKRFMRFGRN--PSELEDRIEE-GK-------------------- : 209
Octopus.b        : R----E------------PDKRFMRFGKS---D--DKRFMRFGRN--PNELEDRLEE-GK-------------------- : 210
Doryteuthis.p    : R----D------------PEKRFMRFGKS---D--DKRFMRFGRN--PSDVEDELEE-DK-------------------- : 237
Nautilus.p       : S----E------------KSKRFMRFGKK---D--EGAGIA-KRN--ADD-QEEFVE-NK-------------------- : 186
Aplysia.c        : KSVGSD-----------EVDKRFMRFGKSVGSDEVDKRFMRFGKSLGTDDVDKRFMRFGK----SLGTDD-------VDK : 168
Lymnaea.s        : R-GDEE------------AEKRFMRFGKS---------FMRFGRD--MSDVDKRFMRFGK-------------------- : 173
Mytilus.e        : KRGDEDFGEEEGDDTYGVEDKRFMRFGRG-GTE--DKRFMRFGRA-GE---DKRFMRFGKRADENFGEDEGDETYGVEDK : 275
Crassostrea.g    : E--VSE------------ENKRFMRFGRT------------------PAEDDPTYMK----------------------- : 177
                                       KRFMRFG4          f rfg                k

                            *       340         *       360         *       380         *       400
OvClone197       : RFMRFGR--GNDEE----EKRFMRFGR----D-P--DSKLMGYGNS---DE----------------------EKRFMRF : 251
Octopus.b        : RFMRFGR--GNEEE----EKRFMRFGR----D-P--DSKLMGYGNS---EE----------------------EKRFMRF : 252
Doryteuthis.p    : RFMRFGR--GGEDDEEEAEKRFMRFGR----D-P--EKKFMRFGKS---GE----------------------DKRFMRF : 283
Nautilus.p       : RFMRFGR--GD-------EKRFMRFGR----DNA--DGR--QVGD----WD----------------------DKRFMRF : 223
Aplysia.c        : RFMRFGKSLGTEDV----DKRFMRFGKSLGTDDV--DKRFMRFGKSLGTDD-----VDKRFMRFGKSLGTEDVDKRFMRF : 237
Lymnaea.s        : RFMRFGREPGT-------DKRFMRFGR----E-PGADKRFMRFGKS---FDGEEENDDDLYYNESDADSNDDVDKRFMRF : 238
Mytilus.e        : RFMRFGR--GGTE-----DKRFMRFGKSMDNDD---EKRFMRFGKS---AE-----ADKRFMRFGKSLEA---DKRFMRF : 334
Crassostrea.g    : RFMRFGR---NPDL----EKKFMRFGK----DGN--EKRFMRFGKR---EDNDNMVTD---------------DKRFMRF : 226
                   RFMRFG4  g         K4FMRFG4    d      4 m  G s                           dKRFMRF

                            *       420         *       440         *       460         *
OvClone197       : GRMSDEADAQKRFMRFGRSVDDDKAN----------------RLKKSNDQTDC------------------------ : 288
Octopus.b        : GRMSDEADAQKRFMRFGRSVDDDKANRLK----KSNDQLRTIRMGRSVDDKKVN--------SANGDAYLRIGQSDE : 317
Doryteuthis.p    : GRNPDEQEADKRFMRFGRGGEDDEV---------STEDKRFMRFGRSADKCK--------------------GCLEG : 331
Nautilus.p       : GRNPDE----KRFMRFGRGLENEDE---------MEEEKRFMRF--------------------------------- : 254
Aplysia.c        : GKSLGTDDVDKRFMRFGKSLGTEDVDKRF-----MRFGKRFMRFGRSVGDSKYRGASSENVMTTDSKQTTEQATNKS : 309
Lymnaea.s        : GKSAEE----KRFMRFGKSQDA------------SRDKKEFLRIGKRESRSA------------EVENNIQIAAKQS : 287
Mytilus.e        : GKSGDD---EKRFMRFGKSVDGEDKEKRFMRFGKSTEDKRFMRFGRDPAEKRFMRFGKS---TTEDKRFMRFGRK-- : 403
Crassostrea.g    : GRDPKDDTLMERFVRNGRSGDD----------------KRFMRFGSTAKKIISR----------------------- : 264
                   G4        kRF6RfG4s                   krf R g
Bradykinin

                         *        20         *        40         *        60         *        80
OvClone194    : -------------------------HSLPKRNDHVITLRSLRQTGLSDSDSRALLQAY-----------ILGKLSNVDIA :  44
Octopus.b     : ------MGINIIHVLCLVAVFASSVHSLPKRNDHVITLRSLRQTGLSDSDSRALLQAY-----------ILGKLSNVDIA :  63
Octopus.v     : ------MGINIIHVLCLVAVFASSVHSLPKRNDHVITLRSLRQTGLSDSDSRALLQAY-----------ILGKLSNVDIA :  63
Sepia.o_NKY3  : ------MTVNAVHVLCIFALLFACAHSLPKRTDHASTLRYLQQSGLSDSDSRALLQAY-----------LIGKLSNGD-S :  62
Doryteuthis.p : ------MTVNAVHVLCIFALLFACVHSLPKRTDHASTLRFLQQSGLSDSDSRALLQAY-----------ILGKLSVGDGS :  63
Conus.g       : -MLTIPHFLSASAILLLVSCALARSLNDETLPKEMESLVAGSGSNNNDPYNDPKVLADLWD--------LLLTGSHVTPY :  71
Clione.l      : -MTSSIYGVLTLAVAAFISHVSCRSMDYYDSEADSYDFPPNDAYRLKS--PGEMLAVIPNQRQYQELLLSMGEPSGMSPY :  77
Helix.l       : -MTSSIYGFITLSVVALISQTTCRSLDLLLDGDFNNGLASFDGSSKWSRLPLEFLAAFDLDP-HQAQGQHLAEAPEAPLM :  78
Tritonia.d    : -MSTLTPSLNHCAILALLSLLLTTYVTARSAQASSDHTEEKRFYIPVVISSDAILASLKN--------HQVMGGPELTRE :  71
Aplysia.c     : -MTSSIYGFITLSVVALISQTTCRSLDLLLDGDFNNGLASFDGSSKWSRLPLEFLAAFDLDP-HQAQGQHLAEAPEAPLM :  78
Aplysia.b     : -MTTSIYGFITLSVVALISQTTCRSLDLLLDGDLNNGLASFDGSSKWSRLPLEFLAAFDLDP-HQVQGQHLAEAPEAPLM :  78
Pinctada.m    : MKTTILLSFSLCVTMTFAKSLSESFRGEWKQDRPSRILSMISKSQQKELELIQTLMRMTADR----NKSLFFDEEDYDPS :  76
                                                     l                6

                         *       100         *       120         *       140         *       160
OvClone194    : GNKEADNSD---------------------------YTTIKRKAFWRPMGYLPFDNH-GGSGASSSNDNSPSGGTGSAVF :  96
Octopus.b     : GNKEADNSD---------------------------YTTIKRKAFWRPMGYLPFDNH-GGSGASSSNDNSPSGGTGSAVF : 115
Octopus.v     : GNKEADNSD---------------------------YTTIKRKAFWRPMGYLPFDNH-GGSGASSSNDNSPSGGTGSAVF : 115
Sepia.o_NKY3  : ICKELETSE---------------------------YPTIKRKAFWRPMGYLPFENH-VGSGASSSNDNAAGTGSASAVF : 114
Doryteuthis.p : IGKELETSE---------------------------YPTIKRKAFWRPMGYLPFENH-AGSGASSSNDNAAGGGSASAVF : 115
Conus.g       : GQPALSMVS----------------AKRSWPNGYLVPSSMKRKMFWTPLGHLPASAR-LGRPQNMRPNME---DSGSPVF : 131
Clione.l      : GGLQRTHVPNQLQLLASGLQGGIKRAQRQGLWRHPLFRNIKRKMFWQPLGYMPASQR-THN-ADMARDSS-SNDSGSKML : 154
Helix.l       : EAMKRSRGP-------------SPRRLRSYLRRAAGLRGMKRKMFWQPLGYMPASAR-AHNNVPEVVNEN-SQDSGTNVF : 143
Tritonia.d    : NGLERKRTAAAR--------------LEYLDSLLPRSTVPKRKMFWSPLGYMSASARRLNQNNPGSRPAASKEDTGSPGF : 137
Aplysia.c     : EAMKRSRGP-------------SPRRLRSYLRRAAGLRGMKRKMFWQPLGYMPASAR-AHNNVPEVVNEN-SQDSGTNVF : 143
Aplysia.b     : ETMKRSRGP-------------SSRRLRSYLRRAAGLRGMKRKMFWQPLGYMPASAR-AHNNVPEVVNEN-SQDSGTNVF : 143
Pinctada.m    : MESEEDKMKN--------------------ILSEDVKSVSKRKVFWQPLGYVPASLR--MNGGSREQSGESSSRAGGNIL : 134
                                                        KRK FW P6Gy6p                      g   f


OvClone194    : RYG :  99
Octopus.b     : RYG : 118
Octopus.v     : RYG : 118
Sepia.o_NKY3  : RYG : 117
Doryteuthis.p : RYG : 118


Conopressin

                              *        20         *        40         *        60         *        80
OvConopressin508Cl : ---------MSSIKSSVFAIL--IVVVLLPLVK-GCFWTSCPIGGKRSNI-PATEP---RQCMSCGPNGEGQCVGSNICC :  64
Octopus.b_Cono/Neu : -----MSQNCFAIVQLLFVLF-TVCSFFIATTD-GCYFRNCPIGGKRAT--PMSEQGSNQKCMSCGPNGEGQCVGSNICC :  71
Octopus.v_Cephalot : -----MSQNCFAIVQLLFVLF-TVCSLFIATTD-GCYFRNCPIGGKRAT--PMSEQGSNQKCMSCGPNGEGQCVGSNICC :  71
Octopus.v_Octopres : ---------MSSIKSSVFAIL--IVVVLLPLVK-GCFWTSCPIGGKRSNI-PATEP---RQCMSCGPNGEGQCVGSNICC :  64
Doryteuthis.p      : --------MASYRWGSLALLLIIVVLPLVSIVE-GCFWTECPIGGKRSS--AAV-----RECMACGPEGKGRCAGPSICC :  64
Sepia_Sepiatocin   : -MGSGRFLFSSTKCQVACVLF-NFCVFLICTTD-ACFFRNCPPGGKRAVA-MNDGVAH-KQCMACGPEGKGRCAGPNICC :  75
Nautilus.p         : -------MICGQSCQVVVVLL-ALGASYIVEVE-GCYFLSCPVGGKRSSVGSSSNTGMNHQCQPCGPNGQGQCFGPQICC :  71
Aplysia.c_Cono     : --MSHSSMSPLSVRTFVLVAG-LAVISFSVVAD-ACFIRNCPKGGKRSM--DMQLLGQ-RQCMACGPEGIGQCVGPNICC :  73
Aplysia.k_Lys-cono : -------MSPLSVRTFVLVAG-LAVISFSVVAD-ACFIRNCPKGGKRSM--DMQLLGQ-RQCMSCGPDGIGQCVGPNICC :  68
Lymnaea.s          : --------MMSSLCGMPLTYLLTAAVLSLSLTD-ACFIRNCPKGGKRSL--DTGMVTS-RECMKCGPGGTGQCVGPSICC :  68
Lottia.g           : --------------------------------N-SCFIRNCPTGGKRAI--EASEIG--HKCMSCGPGNVGQCVGPNICC :  43
Conus.l_cono       : ---MKCSVLQMSRLSWAMCLM-LLMLLLLGTAQ-GCFIRNCPRGGKRAV--DAVQPA--RQCMSCGPDGVGQCVGPSVCC :  71
Crassostrea.g_cono : ---------------------------MLQIVD---------------------------TCMSCGPGLLGQCVGPDICC :  26
Lingula.a_cono/neu : ---------MLPCSTNMLLKHILALMCALTIAS-GCFIRNCPSGGKRTM--EDFDTSHHRQCMSCA-DGRGQCVGPSICC :  67
Platynereis.d_vaso : --------MQFSRPTFTLQYGSAVLLTLVVCCS-ACFVRNCPPGGKRSM--DLPQIHSTRQCMRCGPQGLGQCFGPNICC :  69
Capitella.t        : MRHSSDAVNSVVLCGRLLLLF-ACISCCIETTS-GCFIRNCPIGGKRSSV-PSRISAQ-KECMACGPNGLGQCVGPNTCC :  76
Asterias.r         : -----------MGMKSMVALWTGVLVALWVQSQ-ACLVQDCPEGGKRSS--YNTI----RQCLSCGPGGLGQCVGSAICC :  62
Strongylocentrotus : ---------MMSVKSIVTCLFLSLVLALWIGGSFACFISNCPKGGKRSNSRPL------RQCLECGPGGVGRCMGPGICC :  65
Danio.r_vaso/neuro : -----------MSDSLLSVCV-LCVLALSTLSS-ACYIQNCPRGGKRSQ--PEPI----RQCMSCGPGGVGRCFGPSICC :  61
Xenopus.t_vaso/neu : -----------MPEASVPACF----LCLLALSS-ACYIQNCPRGGKRSY--PDTEL---RQCMQCGPGNRGNCFGPNICC :  59
Rattus.n_Oxy       : -----------MACPSLACCL----LGLLALTS-ACYIQNCPLGGKRAA--LDLDM---RKCLPCGPGGKGRCFGPSICC :  59
Bos.t_vaso/neuro   : -----------MPDATLPACF----LSLLAFTS-ACYFQNCPRGGKRAM--SDLEL---RQCLPCGPGGKGRCFGPSICC :  59
Mus.m              : -------MLARMLNTTLSACF----LSLLAFSS-ACYFQNCPRGGKRAI--SDMEL---RQCLPCGPGGKGRCFGPSICC :  63
Homo.s_Vaso        : -----------MPDTMLPACF----LGLLAFSS-ACYFQNCPRGGKRAM--SDLEL---RQCLPCGPGDKGRCFGPSICC :  59
                                                        c    cp ggkr              C  Cgp g G C Gp  CC

                              *       100         *       120         *       140         *       160
OvConopressin508Cl : HKD-GCIIGTLAKE-CNEENESTTACSVKGVPCGTDG-QGRCVADGVCCDESACSSNARC------------NTRGGRSK : 129
Octopus.b_Cono/Neu : HKD-GCIIGTLAKE-CNEENESTTACSVKGVPCGTDG-QGRCVADGVCCDESACSSNVRC------------NTSGGRSK : 136
Octopus.v_Cephalot : HKD-GCIIGTLAKE-CNEENESTTACSVKGVPCGTDG-QGRCVADGVCCDESSCFTTDRC------------DRENHRSM : 136
Octopus.v_Octopres : HKD-GCIIGTLAKE-CNEENESTTACSVKGVPCGTDG-QGRCVADGVCCDESSCFTTDRC------------DRENHRSM : 129
Doryteuthis.p      : SND-GCIIGEMAKE-CMQEDEGTTACEVKGMPCGAEG-QGRCTALGVCCDTTACSSNSHCGPVP--------TRSSNQRQ : 133
Sepia_Sepiatocin   : QKE-GCIIGDMAKE-CMQEDEGTTVCEVKGIPCGAEG-QGRCVAAGVCCDTSACSTNSHCGSALP-------RTSSRRQE : 145
Nautilus.p         : HSS-GCVVGQLASE-CEKENESTTPCLVTGKSCGSEG-QGHCGAHGICCDNEACSANSHCKESAE-------VVPYNRQE : 141
Aplysia.c_Cono     : SPHFGCHIGTPETEICQKENQSTSPCSVRGETCGYRD-SGNCVANGICCDSESCAANDRCRLRKETSRIGFDDTQSSRAE : 152
Aplysia.k_Lys-cono : SPHFGCHIGTPETEVCQKENQSTSPCSVRGETCGYRD-TGNCVANGICCDSESCAVNDRCRLRKETSRSGFDDTQSSRAE : 147
Lymnaea.s          : GQDFGCHVGTAEAAVCQQENDSSTPCLVKGEACGSRD-AGNCVADGICCDSESCAVNDRCRDLDG-------NAQANRGD : 140
Lottia.g           : GR-FGCYIGTKETEICEHENDSTVACRVEGKLCGSRQ-QGQCVANGICCDS----------------------------- :  92
Conus.l_cono       : GLGLGCLMGTPETEVCQKENESSVPCAISGRRCGMDN-TGNCVADGICCVEDACSFNSLCRVDTDQE-----DSVSARQE : 145
Crassostrea.g_cono : GP-FGCQMGTSESNICGKENESTTACAISGPPCGSRN-QGNCVADGICCDTGACSFNTKCKL----------NSEPRDVQ :  94
Lingula.a_cono/neu : VEKGGCHVGTSEADVCKKENESNTPCSVKGKSCDSSLFMGKCTADGICCNSESCVMDDNCRTA---------FSSKYSKE : 138
Platynereis.d_vaso : GPSIGCYINTLESEECSKENDVRTPCDITAEICGVDG-QGRCGADGVCCTDEKCTLDSSCDKLDM-------DKPRFSPD : 141
Capitella.t        : GQDIGCFMGTQEAKMCGEENDSPIPCRVDGAACGRND-GGRCVAEQICCNEDKCSHDSSCQSKAKR------QHDNLSQD : 149
Asterias.r         : GNTFGCFLGTKETFVCREESQLSTPCEVVGETCESIT-DGKCVSNGFCCNERSCSLDVACR-----------ETDTEQRD : 130
Strongylocentrotus : GPTIGCHINTQHTLSCMRENEISTPCELPGNPCQTVP-SGTCGAMGVCCNSNSCSEDASCLMIEE-------DDSLKRFE : 137
Danio.r_vaso/neuro : GPGLGCVLGSPEAQVCMEEEQLSGPCETGGTSCGDRG--GRCAAEGICCDSESCAVDPDC-------------PEGSSAG : 126
Xenopus.t_vaso/neu : GEDMGCYIGTPETLRCVEENFVPSPCEAGGRPCST-G--GRCAAPGICCNDESCSLDSACL-----------DDESERRR : 125
Rattus.n_Oxy       : ADELGCFVGTAEALRCQEENYLPSPCQSGQKPCGS-G--GRCATAGICCSPDGCRTDPAC------------DPESAFSE : 124
Bos.t_vaso/neuro   : GDELGCFVGTAEALRCQEENYLPSPCQSGQKPCGS-G--GRCAAAGICCNDESCVTEPECREGVGFPRRVRANDRSNATL : 136
Mus.m              : ADELGCFVGTAEALRCQEENYLPSPCQSGQKPCGS-G--GRCAAVGICCSDESCVAEPECH--DGFFRLTRAREPSNATQ : 138
Homo.s_Vaso        : ADELGCFVGTAEALRCQEENYLPSPCQSGQKACGS-G--GRCAAFGVCCNDESCVTEPECREV--FHRRARASDRSNATQ : 134
                         GC 6g      C  E      C   g  Cg     G C a g CC    c     c

                              *       180         *
OvConopressin508Cl : TLK-------------DLLLILNKMYRDKN--------- : 146
Octopus.b_Cono/Neu : TLK-------------DLLLILNKMYRDKN--------- : 153
Octopus.v_Cephalot : AMQ-------------KLLEIRDGIYYKK---------- : 152
Octopus.v_Octopres : AMQ-------------KLLEIRDGIYYKK---------- : 145
Doryteuthis.p      : LLNL---------LK-SLIKKVK---------------- : 146
Sepia_Sepiatocin   : LFSL---------LK-RLINKVN---------------- : 158
Nautilus.p         : LLAL---------LK-RIIKRVEN--------------- : 155
Aplysia.c_Cono     : VLKL---------IQ-KLLRAKEED-------------- : 167
Aplysia.k_Lys-cono : VLKL---------IQ-KLLRAKEED-------------- : 162
Lymnaea.s          : LIQL---------IH-KLLKVRDYD-------------- : 155
Lottia.g           : --------------------------------------- :   -
Conus.l_cono       : LLTL---------IR-RLLVNRQYD-------------- : 160
Crassostrea.g_cono : ILSL---------LK-TLGLIETKAYNNAGMS------- : 116
Lingula.a_cono/neu : FLLF---------VK-NLLESAKKDQH------------ : 155
Platynereis.d_vaso : ILRM---------LK-KIFDRRSYPGRRK---------- : 160
Capitella.t        : LLRY---------MH-QLMTLKSLGGRR----------- : 167
Asterias.r         : LKNR---------LKERLLDALLRQP------------- : 147
Strongylocentrotus : QMSREENGSTRKDLRVKLLDLLLNMQDQ----------- : 165
Danio.r_vaso/neuro : LKSI---------SGETLLRLLNLATRRQRPF------- : 149
Xenopus.t_vaso/neu : APLEKNTTVMDGSASDFLLRLMHMANRQQQAKHQYY--- : 161


 Myomodulin-1-like

                         *        20         *        40         *        60         *        80
OvClone198    : -------------------------------------------------------------------------------- :   -
Octopus.b     : -MLFNITPIICALVLCVLFPRGNCNEDTNASQSKSKLNENNVA----EQHLDKRSSGEYNP---------------HDLK :  60
Doryteuthis.p : -MNLTLKLICIVLCLQ-IQQGACESNDNNNNNHDVNAAEAGAS----PLLRERRAVGMLRLGRGVQMLRLGKRAPYDDLK :  74
Sepia.b       : -MNLTLTLICVLLCLQQLRQGACEDNESNNNNN--NAVETGAT----PALRERRAVGMLRLGRGVQMLRLGKRAPYDDLK :  73
Nautilus.p    : IIRSRCWAALISLISILLTKSLRFRKFVVRGFRPIRRIYKRPVQIETVITSKASSLCKMRFSVTLVCLAVCLHS------ :  74
Aplysia.c     : -MQVYMLLPLAVFASLTYQGACEETAAAQTSSDASTSSASSEHAENELSRAKRGSYRMMRLGRGLHMLRLGK-------- :  71
Lymnaea.s     : -MQGAFLITFIVLTMTLVSIGNTEESGGQTSSNDKTEPAQ--------SRTKRRSREMGRVVRGLQMLRLGK-------- :  63
Capitella.t   : -MKVALLFSLQCIYVIASADDSARARRDLSMLRMGKRSEFSSLPPLVPPSYDLSADDFDERQVWMKIPRVGKD------- :  72
Charonia.t    : -MMKKILLSLTLCFLFQLHRANSEEASAAAETDNDSSSNSGE----ELSRLKRNAMDMRRLGRGIQMLRLGK-------- :  67
Haliotis.a    : -MNAALFAKAVTVLLLVICKQGSSEE--QTQSESQDEPAS---------REKRG-LNMLRLGRGLQMLRLGK-------- :  59


                         *       100         *       120         *       140         *       160
OvClone198    : -------------------------------------------------------------------------------- :   -
Octopus.b     : TIVAAILERQEQGQKQSSALRSMN---ELAGDQYNGDALSRVVQLLRRSMPSDFD--DFERRLAPVPRLGRLKKRS---- : 131
Doryteuthis.p : TIVASILNQQDQQ-NRQAPLPRYGKEEDLLTEAYPAESLSQVTQLLRRSNPSYIDEDDLLRQEAPVPRLGYMHKRS---- : 149
Sepia.b       : TIVATILGRQEQQFNRQAPLPRYGKEEGLLVDAYPADSPSQVSQILRRSYPSYFEDEDLLHQEAPLPHLGYLQKRSAEIR : 153
Nautilus.p    : --------------------------------------WEGSCENSAEN-----------PNGLLRVRRGGLNMLRLG-- : 103
Aplysia.c     : ---------------------------------------RGGP-VEPES------------EENLETLLNLLQGYY---- :  95
Lymnaea.s     : ---------------------------------------RDSV-STSED------------PGDIDDILSLLQAYQ---- :  87
Capitella.t   : --------------------------------------IDEDTPHLRLA------------RYPPVPRLGSALDSL---- :  98
Charonia.t    : ---------------------------------------RGVP-MLRLG------------RSSPEDTLS-LEDLL---- :  90
Haliotis.a    : ---------------------------------------RGGLNMLRLG------------RSSPS--SDELSDYL---- :  82


                         *       180         *       200         *       220         *       240
OvClone198    : --------------------------------------DNILFPRQITLPRYGKDEGSDAADAEQYLSSDCEIFDAYGTC :  42
Octopus.b     : ---------------VLSNPNKEVDYDEIEANDNNESADNILFPRQITLPRYGKDEGSDAADAEQYLSSDCEVFDAYGTC : 196
Doryteuthis.p : -APLPRYGKGPDYEDLSNEATSEEDSTDGEEE--SEVENSHLFDRQPPLPRYGKDE---------LVGLECEAYDESGNC : 217
Sepia.b       : HAPLPRYGKGPDYEDLTNEVSSEEGGDDHEDENVSDTVSSEMFERQPPLPRYGKDE---------IAGIDCEAYDESGNC : 224
Nautilus.p    : -------------------------RGVHMLRLGKRNSGGDLSTEELRTLILALLQR-------------HRMFSERQTP : 145
Aplysia.c     : -------------------------SDVPEYPSEFDDTDLAYPYEEYDAPAHPRYR---------------RSTPPTDGV : 135
Lymnaea.s     : -------------------------EQNPEFA--FNEEERELSGDELEVPEHHRFR---------------RSTENSG-- : 123
Capitella.t   : -------------------------IEEYRRDIGSNDDEDEIRQVLFKIPRVGRNR---------------RSADDQENV : 138
Charonia.t    : -------------------------NAMEESQ----YLDDFYPYPLPEEPVHGRFR---------------RSADHSQES : 126
Haliotis.a    : -------------------------YDLLQNEY-LVNPEVLNGEDDMMYPDSGRYR---------------RSADGVRAV : 121
                                                                 p

                         *       260         *       280         *       300         *       320
OvClone198    : VQYK-IPRENVK-----RQVRMLRLGKRQMHPIKIDDE-TGMDKRSE---DFNVDGNSGESKRAVSMLRLGRS------- : 105
Octopus.b     : VQYK-IPGENVK-----RQVRMLRLGKRQMHPIKIDDESTGMDKRSE---DFNADGNSGENKRAVSMLRLGRS------- : 260
Doryteuthis.p : LGYNDIEKKDVRMLRMGRQVNMLRMGKRALSMLRMGRN--AENKRAVGMLRLGRSDNTDESKRALAMLRLGRSGEKRAVS : 295
Sepia.b       : LRFEDIEKKDVRMLRMGRQVNMLRMGKRALSMLRMGRN--GGDKRAVSMLRLGRSDNVEDSKRALAMLRLGRSGEKRAVS : 302
Nautilus.p    : LPRYGKDIIGDS-----DFDSNQDDALTRVAQLLRRSR---------VYPSQFSDDDEEYFSSLPPSPRLGRIKRS---- : 207
Aplysia.c     : VAPDVLQKGSSE----FEDFGDSQLDESDEGYYGYDPE-NYLYGDFEDYLEPEEG-GLGEEKRSLSMLRLGKRGLS---- : 205
Lymnaea.s     : VAPQEVQQSQS-----FKDSGEHELKLEEAEPYLYFPDGDFYYGDVDELLEGDNE-DGSADKRQIPMLRLGKRSMS---- : 193
Capitella.t   : EMKEQVKRAAP-----LPRLGMLEERAAPLPRLGLYER--------AAFLPRLGYRDLDEEERAAPLPRLGVREED---- : 201
Charonia.t    : VPADSVFADGA-----NKSAGDVKRSVDETAEPLEEEE---------SYFNEQGD-GDDIEKRPMSMLRLGKRPMS---- : 187
Haliotis.a    : ALPRIG-----------------------------------------KELEDEEY-GDEYEKRALSMLRLGRSGEG---- : 155
                                                                             kr    lRLG

                         *       340         *       360         *       380         *       400
OvClone198    : ---FNAGGESEENEIPVFDEAKRAVSMLRLGRSGPFLDKRALSMLRLGRSEFDQENTALPIMYPGPNGFESNKRAVSMLR : 182
Octopus.b     : ---FNAGGQSEENEIPVFDEAKRAVSMLRLGRSGPFLDKRALSMLRLGRSEFDQENTALPILYPGQNGFENNKRAVSMLR : 337
Doryteuthis.p : MLRLGRSGLGEDN-MPV-EEQKRALAMLRLGRSGSDEQKRAVSMLRLGRSGEGEEKRAVSMLRLGRSGPDSEKRAVSMLR : 373
Sepia.b       : MLRLGRSGLDEES-MPS-EEQKRALAMLRLGRSGSEEQKRAVSMLRLGRSGEGDQKRAVSMLRLGRSGPDGQKRAVSMLR : 380
Nautilus.p    : -------AENSVEDIENQKDFERAPPIPRIGRLIDDEDVDDSDESSFNRRMT-------PLPRFGWCEFYDDNGDCLSFD : 273
Aplysia.c     : ---MLRLGKREGE-EGDEMDKKQDESLN--DAFENDDIKRTLSMLRLGKRP-------MSMLRLG-------KRPMSMLR : 265
Lymnaea.s     : ---MLRLGKREED-DADFDEEKRSLSMLRLGKREDDNFEDSFDEE-NDKRS-------LSMLRLG-------KRPMSMLR : 254
Capitella.t   : ---YENEVGDNSY-LKEED--ERAAPLPRLGYYE----KRNVGMLRMGKRP-------MSMLRMG-------KRPMSMLR : 257
Charonia.t    : ---MLRLGKRPMS-MLRLG--KRPMSMLRLGKRP-------MSMLRLGKRP-------MSMLRLG-------KRPMNMLR : 240
Haliotis.a    : ---VEDEAEGLEP-EEEEE--KRALGMLR-----------------LGKRP-------MNMLRLG-------KRPMNMLR : 198
                                     kr   6 r g                g4           6 r G       kr   mlr

                         *       420         *       440         *       460         *       480
OvClone198    : LGRS------------------------MSADDKRAVSMLRLGRSGFDTMKRAVSMLRLGRN---SGYPSEKRAVSMLRL : 235
Octopus.b     : LGRS------------------------MSAGDKRAVSMLRLGRSGFDTMKRAVSMLRLGRN---SGYPSEKRAVSMLRL : 390
Doryteuthis.p : LGRNS----------------------PSYDAEKRAVSMLRLGRSGSDEEKRAVSMLRLGRSG--AGEEADKRAVSMLRL : 429
Sepia.b       : LGRNF----------------------PVSDAEKRAVSMLRLGRSGSDEEKRAVSMLRLGRS---AEEEAEKRAVSMLRL : 435
Nautilus.p    : RSTRG------------------------------GLGMLRLG-------KRRLNMLRMG-----------KRHLSMLRL : 305
Aplysia.c     : LGKR-------------------------------PMSMLRLG-------KRPMSMLRLG-----------KRPMSMLRL : 296
Lymnaea.s     : LGKR-------------------------------PMSMLRLG-------KRPMSMLRLG-----------KRPMSMLRL : 285
Capitella.t   : MGKR-------------------------------PMSMLRMG-------KRPMSMLRMGRSMDEAQPEQQKRAMSMLRM : 299
Charonia.t    : LGKRDGEDAEPIDEEEFIEPFEEEGQVEEFPAEKRPMSMLRLG-------KRPMSMLRLG-----------KRPMSMLRL : 302
Haliotis.a    : LGKR-------------------------------PMNMLRLG-------KRPVNMLRLG-----------KRPMNMLRL : 229
                 g                                  6sMLR6G       KR 6sMLR6G           KR 6sMLR6

                         *       500         *       520         *       540         *       560
OvClone198    : GRSGSDEDKRAVSMLRLGRS-GSDED---------------------KRAVSMLRLGRSGADIEDEKRAVSMLRLGRGGA : 293
Octopus.b     : GRSGSDEDKRAVSMLRLGRS-GADIEDEKRAVSMLRLGRSG--ADNMKRAVSMLRLGRSGSDDMNTKRAVSMLRLGRSGN : 467
Doryteuthis.p : GRSGSDGKKRAVSMLRLGRS-GPEQDESKRAVSMLRLGRSGPESDETKRAVSMLRLGRSGSESDQSKRAVSMLRLGRSGS : 508
Sepia.b       : GRSGAD--KRAVSMLRLGRR-NQGADDEKRAVSMLRLGRSGPESDESKRAVSMLRLGRSGPETDESKRAVSMLRLGRSGP : 512
Nautilus.p    : G-------KRQDEELAGE-----------------------------KRSLRLLRLG---------KKSESEDSHTIKN- : 339
Aplysia.c     : G-------KRPMSMLRLG----------KRPMSMLRLG---------KRPMSMLRLG---------KRPMSMLRLGKR-- : 339
Lymnaea.s     : G-------KRPMSMLRLG----------KRPMSMLRLG---------KRPMSMLRLG---------KRPMSMLRLGKR-- : 328
Capitella.t   : G-------KRGMNLLRMGRSEQPAEEEEKRAMSMLRMGRSEPAVEAEKRPMSMLRMG---------KRPMSMLRMGKREM : 363
Charonia.t    : G-------KRPMSMLRLG----------KRPMSMLRLG---------KRPMSMLRLG---------KRPMSMLRLGKR-- : 345
Haliotis.a    : G-------KRPMNMLRLG----------KRPMNMLRLG---------KRPMNMLRLG---------KREDETEGEEKR-- : 272
                G       KR    Lr g          kr   mlr g         KR 6s6LR6G         K4  smlr g

                         *       580         *       600         *       620         *       640
OvClone198    : DNM---------------------KR------------------------------------------------------ : 298
Octopus.b     : DNVGE------------------DKRAVSMLRLGRSDNNA-NNKRAMAMLRLGRSNDTSSKET----------------- : 511
Doryteuthis.p : EADESKRAVSMLRLGRSD---SADKRAVSMLRLGRSDKNSDAEKRAVSMLRLGRSDHATDVKQS---------------- : 569
Sepia.b       : ETEESKRAVSMLRLGRSDEKFGADKRAVSMLRLGRSDKNTDADKRAVSMLRLGRSAEEEAEKRAVSMLRLGRSGADKRAV : 592
Nautilus.p    : -------------------------RALSMLRLGRSEQTSEDKQR----------------------------------- : 359
Aplysia.c     : --------------------------PMSMLRLGKRDDDEKEKKSLNMLRLGKRSTQ----------------------- : 370
Lymnaea.s     : --------------------------PMSMLRLGKREDDE-EKRSLAMS------------------------------- : 350
Capitella.t   : DDEVIPVV-------------EAEKRPMNMLRMGKRDTDDNQPIVEEDQQAQSS-------------------------- : 404
Charonia.t    : --------------------------PMSMLRLGKRESE----------------------------------------- : 358
Haliotis.a    : --------------------------ALGMLRLGKRSDEKVDGDAGVSTQQ----------------------------- : 297
                                             mlr g

                         *       660         *       680         *       700
OvClone198    : ------------------------------------------------------------------ :   -
Octopus.b     : ------------------------------------------------------------------ :   -
Doryteuthis.p : ------------------------------------------------------------------ :   -
Sepia.b       : SMLRLGRRNQGADEEKRAVSMLRLGRSGPESDESKRAVSMLRLGRSGPETDESKRAVSMLRLGRSG : 658
Nautilus.p    : ------------------------------------------------------------------ :   -
Aplysia.c     : ------------------------------------------------------------------ :   -
Lymnaea.s     : ------------------------------------------------------------------ :   -
Capitella.t   : ------------------------------------------------------------------ :   -
Charonia.t    : ------------------------------------------------------------------ :   -
Haliotis.a    : ------------------------------------------------------------------ :   -


Sp191
                              *        20         *        40         *        60         *        80
OvClone191         : -------MKTYQSRNLCILSCVSLILVLCCG---------------VDASKSLTKRLGNGFKNNILDRMAHGFGKRTLED :  58
Octopus.b          : -------MKTYQSRNLCILSCVSLILVLCCG---------------VDASKSLTKRLGNGFKNNILDRMAHGFGKRTLED :  58
Doryteuthis.p      : -------MNVSGAWGLCTLCCLGLLVLFASSD--------------AHASVVVPNRPQRGFKDNVSNRIAHGFGKRTFQD :  59
Sepia.o            : -------MNVSGARGLCTLCCLGLLVLLASSD--------------AHASVVVPNRPQRGFKDNVSNRIAHGFGKRTFQD :  59
Nautilus.p         : -------MKFTLS-----LCCLVLAVVTQAYP--------------QPEAHSREKR---AFKDSVANRIAHGFGKRTALS :  51
Charonia.t_Allatot : ----------MMRTSLVLIGLVVIALGQALP-----------------THS--NSRQKRGFRINSSSRVAHGYGKRTFNT :  51
Crassostrea.g_Alla : ----MQMVKVVLVTVLLTLCVIVDCLPHSS------------------TKMR----QKRGFRQSIVDRMGHGFGKRANTD :  54
Aplysia.b_Allatotr : ----MLSAPSIAHTVVALLVLMCLCPFSQST-----------------EAS--LSRAKRGFRLNSASRVAHGYGKRGYAS :  57
Aplysia.c_Allatotr : ----MLSAPSIAHTGVALLVLMCLCPFSQST-----------------EAS--LSRAKRGFRLNSASRVAHGYGKRGYAS :  57
Capitella.t_Allato : -----------MKVSICFIVVALVVCIEVMT-----------------SHAANLSRSKRGFRMGAADRFSHGFGKRGGDF :  52
Chaetoderma.n_Alla : ----MYITSSFLVAGLAAMLVFCDAYPKSDDV--------------GLLEMR--DITKRGFRSSVADRIAHGFGKRTPDR :  60
Conus.g_Allatotrop : ---------MRIFISLLLIGLVVLALGQALP-----------------TQS--HSRQKRGFRMNSSNRVAHGFGKRQSGT :  52
Dolabrifera.d_Alla : ----MFSTMSIVQSVALTLLLICLCPLAESR-----------------EVS--LNRAKRGFRLNSASRVAHGYGKRGYAS :  57
Falcidens.c_Allato : ----MYITSSLFVAGLAVMLVCCDAYPKSDDV--------------GLLEMRDGDMSKRGFRSSVADRIAHGFGKRTPVR :  62
Melibe.l_Allatotro : -------------------------------------------------------------------------------- :   -
Platynereisd_Allat : -----------MKVVLCLFVVVFVVSVNGMD-----------------LRAP---RAKRGFRTGAYDRFSHGFGKRGE-- :  47
Aplysia.b_Sensorin : -----------MMVILCIVCLALQAVAANAT---------------------RSKNNVPRRFPRARYRVGYMFGKRSSSE :  48
Aplysia.c_Sensorin : -----------MMVVLCIVCLALQAVAANAT---------------------RSKNNVPRRFPRARYRVGYMFGKRSSSE :  48
Aplysia.d_Sensorin : -----------MMVVLCIVCLALQAVAANAT---------------------RSKNNVPRRFPRARYRVGYMFGKRSSSD :  48
Aplysia.k_Sensorin : -----------MMVVLCIVCLALQAVAANAT---------------------RSKNNVPRRFPRARYRVGYMFGKRSSSE :  48
Biomphalaria.g_Sen : -------MSSNRTSKVHFLPVVLLVLVTCCA---------------------MMD-LASAH----KYRVGYLFGKRG-IE :  46
Clione.l_Sensorin  : ------------MVALCILCLVLHIVVTSAE---------------------QDR-EVPRR--ATRYRVGYMFGKRSPSE :  44
Conus.g_Sensorin   : ------MLMMTTMPRPVLLVLVLLMVHLSLSHCLSLGHLDQPGEDAMAERVGQGAKRDGRRITRGRYRMGYMFGKRSDVE :  74
Dolabrifera.d_Sens : MASYASCLSLQLLVALCVLCLALHNVCVGLA---------------------TERPSVPRRFPRARYRVGYMFGKRSSTE :  59
Helisoma.t_Sensori : ----MCRSSLTLQVGVVLVA-ICLFDVVVSE-----------------DRL--MSRQKRGFRMNSASRVAHGYGKRGY-Q :  55
Lymnaea.s_Sensorin : -------MSPTNMSSG-LVPVLVLTVVACLM---------------------LTDNVSASA----RYRVGYMFGKRG-IE :  46
Melibe.l_Sensorin  : MLPRASPIYSQLMVALCILCLVLHIVVTSAE---------------------QDR-EVPRR--ATRYRVGYMFGKRSPSE :  56
Pleurobranchaea.c_ : -----MTSPTLWTQMLAVVLSACLLSVAHTQ-----------------EAT--LSRHRRGFKKTAASRMGHMYGKRDFLG :  56
Tritonia.d_Sensori : ----MIRSMLSLHIGVVLILLVCIADCLFLK-----------------DKVKDHSRQRRGFRMNTASRVAHGYGKRGYSN :  59
                                                                                        r     gkr

                              *       100         *       120         *       140         *       160
OvClone191         : TYDLAPSIAETGT------------------RNWMTLKDLADLMTYDSDLAYRAALKLD-TNGDGVISMDEFVGGLRGRG : 119
Octopus.b          : TYDLAPSIAETGT------------------RNWMTLKDLADLMTYDSDLAYRAALKLD-TNGDGVISMDEFVGGLRGRG : 119
Doryteuthis.p      : TYDLAPSIDDR--------------------KNLITPRKLAELIMYDKDLAYIVALKLD-TNGDGLISMNELILKDYI-- : 116
Sepia.o            : TYDLAPSLDDS--------------------KNLITPRKLAELIMYDRNLAYIVALKLD-SNGDGVISM-ELILKDYV-- : 115
Nautilus.p         : -LDLLP---EDNS------------------RYWMAIDEVAGRIMTDYKLARTLAQKIVDSNGDGLITSNELLQFSRDL- : 108
Charonia.t_Allatot : MDDSLFPSPSSNG--------------------LMTVQELAQLAAENPSLSEALIQKFIDADGDGIVSTQELFGVAVE-- : 109
Crassostrea.g_Alla : LYDPY----LTKTPN------------------LMTADELTTNILNSEDLAQAIVKKFIDLDGDGIISTAELLRTTS--- : 109
Aplysia.b_Allatotr : SSGAVP--YPELARDVLDNLRAEEEEKE-LEWSIMSVDELASLLQSHPKLARALVKKFVDINGDNLVTAEELFRPPTRK- : 133
Aplysia.c_Allatotr : SSGAVP--YPELARDVLDNLRAEEEEKE-LEWSIMSVDELASLLQSHPKLARALVKKFVDINGDNLVTAEELFRPPTRK- : 133
Capitella.t_Allato : NSLIDGESDM-----------------------VMSDEDLTEIIRADARLAQTFVKRFIDTDGDGFVSRQELFEA----- : 104
Chaetoderma.n_Alla : TSSSQSYRIQSDPPSNS---------------VLMTSEQLTKVLSNSPDLIMKVIQRFIDVNGDGIITVQELQTVHRRK- : 124
Conus.g_Allatotrop : ADDDLFAVVPSSD--------------------LMTVQELAQLSVLHPSLAETLIQKFIDRDGDGILTTQELFDLAVE-- : 110
Dolabrifera.d_Alla : PSSDVT--YPELARDALENLQSMEDEKD-LDWSIMNVDELSSLLQSHPKLARALVKKFIDVNGDNLVTAE---------- : 124
Falcidens.c_Allato : TSSSQ---MLNRPLSNS---------------VLMSSEQLTKVLSNSPDLIMKVIQRFIDVNGDGMITVQELQTVHHRK- : 123
Melibe.l_Allatotro : ---------------------------------MMRLDELSSILRSHPRLLKVLLRKFVDVDGDDVITAEELIRSPMVKL :  47
Platynereisd_Allat : ---LTNEDDG-----------------------LMSVEDMAELITNTPKLALSFVKRYMDRNDDGVISKEELLFVPEQ-- :  99
Aplysia.b_Sensorin : TYSTN-LINLLSR-------------------QLVSQEELRAILEKQPILLDEVVKILDRNDDGYI-TVADLL------- : 100
Aplysia.c_Sensorin : TYSTN-LINLLSR-------------------QLVSQEELRAILEKQPILLDEVVKILDRNDDGYI-TVADLL------- : 100
Aplysia.d_Sensorin : AYSTN-LINLLSR-------------------QLVSQEELRAILEKQPILLDEVVKVLDRNGDGYV-TVADLL------- : 100
Aplysia.k_Sensorin : TCSTN-LINLLSR-------------------QLVSQEELRAILEKQPILLDEVVKILDRNGDGYV-TVADLL------- : 100
Biomphalaria.g_Sen : SPSTS-LIDIVSN-------------------DMKTKQELESLILKKPELLSELITILDRNDDGYI-TGTDLII------ :  99
Clione.l_Sensorin  : LYSVP-LIDLLSR-------------------NLLRQGHRRSSEEKEHPVLEEVFDLVPDSDEDFARMVNDLV------- :  97
Conus.g_Sensorin   : ENPASVLVKKLMG-------------------KAMSVDDLRTALQSDPAFADRVARHLDRNGDGFV-AVSELL------- : 127
Dolabrifera.d_Sens : TISTS-LVNLLSR-------------------QLVSQAELRAILEKESTLLEEVVKILDKNADGFI-TVADLL------- : 111
Helisoma.t_Sensori : LPSSSSSGESQDQLEVLEELSDG---------PLMTVNEFTQLMTSHPNLARALVKKFVDINGDDVISTEELFRPVMKK- : 125
Lymnaea.s_Sensorin : SPSTS-FIDTISR-------------------EMKTKQEVESLILKNPEILSEVLTILDKNDDGYI-TVSDVL------- :  98
Melibe.l_Sensorin  : LYSVP-LIDLLSR-------------------NLLRQGHRRSSEEKEHPVLEEVFDLVPDSDEDFARMVNDLV------- : 109
Pleurobranchaea.c_ : TLASQYGGLSGTG--LDDRMQLPISDVDDADVPIMTVPELTALLTRHEPLARAVVRKFLDLDGDNVVRTGELFRHLQKK- : 133
Tritonia.d_Sensori : IFNDQSNTLSEANKAVPDFLQGGDRSSDNSNSPFLSAQDFSRLIQSNEQLADVIVRKFIDVNDDDLISTDELFRRIRE-- : 137


Buccalin

                          *        20         *        40         *        60         *        80
ObClone614     : MTPSSMWMRVLFCSILLHLYVAQYTRSLETHSEDD------------------------------------------SES :  38
OvClone195     : -------------------------------------------------------------------------------- :   -
Octopus.b      : MTPSSMWMKVLFCSILLHLYVAQYTRSLETHSEDD------------------------------------------SES :  38
Octopus.v      : MTPSSMWMKVLFCSILLHLYVAQYTRSLETHSEDD------------------------------------------SES :  38
Helix.a        : --LRKTPLQQLLLLALVLTLNQISASDVSDTAD----------------------------------------------- :  31
Aplysia.c      : MAHHRGHRHILLYVSLALSLGLALAEDATDPSDDTGSFDDVEAVSEEADLDPYSMSQELNKRPNVDPYSYLPSVGKRAFD :  80
Lottia.g       : MAARK--HELVLVLTSVLCFVSSIVGDPNVPSDSQ---------------DNSALTQD--------------DFAKRGMD :  49
Lymnaea.s      : -------------------------MDRYGFFG----------------------------------------------- :   8
Pinctada.f     : -------------------------------------------------------------------------------- :   -
Biomphalaria.g : -MLPKNWLHSFVLCLVVLSSCRAYDPDTNEFDGER-------------------------------------PEDLVTSD :  42
Crassostrea.g  : MWSTNYATTVFGFFCFVQVFVLTVSQHISSHNDDY-------------------------------------TKHLENIK :  43


                          *       100         *       120         *       140         *       160
ObClone614     : DKRSIIDDDFKSLDPTYYPSGLDKRQ-------------------------QQYSKLGRNIRVIARGMDPMMFG-NLGKR :  92
OvClone195     : -----------------------------------------------------------NIRVIARGMDPMMFG-NLGKR :  20
Octopus.b      : DKRSIIDDDFKSLDPTYYPSGLDKRQ-------------------------QQYSKLGRNIRVIARGMDPMMFG-NLGKR :  92
Octopus.v      : DKRSIIDDDFKSLDPTYYPSGLDKRQ-------------------------QQYSKLGRNIRVIARGMDPMMFG-NLGKR :  92
Helix.a        : SPDAFNDITASKDAADIEHAVLSARE--------------------------DQDSEDPDDV-EKRKLDSYGFYGGIGKR :  84
Aplysia.c      : HYGFTGGLGKRKIDHFGFVGGLGKRQIDPLGFSGGIGKRYDSFAYSAGLGKRGMDSLAFSGGLGKRGMDSLAFSGGLGKR : 160
Lottia.g       : KFGFAGGVGKRGLDKFGFTGQLGKRDMDSFGFAG-------------QLGKRGLDQYGFTGQLGKRGLDQYGFTGQLGKR : 116
Lymnaea.s      : ------GIGKRRLDRFGFYGGIGKRDAGGF----------------------DDWNGATEDL-EKRQLDPFGFSGGIGKR :  59
Pinctada.f     : -------------------------------------------------------------------MDPYMFRGYLGKR :  13
Biomphalaria.g : SLDNSEAMDKRKLDRYGFHMGIGKRD--------------------------DEEDGDLEDVYEKRRIDPFAFSGGIGKR :  96
Crassostrea.g  : FLNKEAEISPKQPADD--DVDFGNSD-------------------------TADDLSDLTEE-EKRALDRYSFSGSLGKR :  95
                                                                                  r 6D   F   6GKR

                          *       180         *       200         *       220         *       240
ObClone614     : -MDPNMFG-SLGKR----FNTDGREKQDKKIDPYMFGSLGKR-MDPAMFG-SLGKR-MDPMLYG-SLGKRMSPDYINYA- : 161
OvClone195     : -MDPNMFG-SLGKR----FNRDDREKQDKKIDPYMFGSLGKR-MDPAMFG-SLGKR-MDPMLYG-GLGKRMSPDYINYG- :  89
Octopus.b      : -MDPNMFG-SLGKR----FNTDGREKQDKKIDPYMFGSLGKR-MDPAMFG-SLGKR-MDPMLYG-SLGKRMSPDYINYA- : 161
Octopus.v      : -MDPNMFG-SLGKR----FNRDDREKQDKKIDPYMFGSLGKR-MDPAMFG-SLGKR-MDPMLYG-GLGKRMSPDYINYG- : 161
Helix.a        : RVDRFLFAEGIGKRQLDPFSFAGHLWKRSIN---NTGGSGNRRLDKNGFAGQIGKRSLDPFGFAGQLGKRGLDKSGS--- : 158
Aplysia.c      : GMDSLAFSGGLGKRGMDSLAFSGGLGKRGMDSLAFSGGLGKRGMDSLAFSGGLGKRGMDSFTFAPGLGKRGMDSLAFAGG : 240
Lottia.g       : GLDQYGFTGQLGKRGLDQYGFTGQLGKRGLDQYGFTGQLGKRGLDQYGFTGQLGKRGLDQYGFAGQLGKRGLDQYGFTG- : 195
Lymnaea.s      : KMDRFSFTGGIGKRRMDRYGFSGGIGKRRLDRFSFTGGIGKRRMDPFSFTGGIGKRRLDRFGFAGGIGKRGLD------- : 132
Pinctada.f     : -MDSRMFSGQLGKRALDRKMFISQLGKR-LDNRMFFGRLGKR-MDYRMFSGQLGKRGLDNHMFVGHLGKRLIP------- :  83
Biomphalaria.g : RLDRFSFAGGIGKRGIDRYGFVAGIGKRRLDRFGFSGGIGKRGIDRFNFAGGIGKRPIDRFSFAGGIGKRGFDRYGFYG- : 175
Crassostrea.g  : GLDRYSFYGGLGKRALDRYGFFGGLGKRALDQYGFAGSLGKRALDRYSFMGGLGKRKLDQYGFAGRLGKRALDRYGFVG- : 174
                  6D   F   6GKR                      G  GkR 6D   F   6GKR 6D   5   6GKR

                          *       260         *       280         *       300         *       320
ObClone614     : -------------------------------------------------------------------------------- :   -
OvClone195     : -------------------------------------------------------------------------------- :   -
Octopus.b      : -------------------------------------------------------------------------------- :   -
Octopus.v      : -------------------------------------------------------------------------------- :   -
Helix.a        : -----------------------------------------------------------------------------AGA : 161
Aplysia.c      : LGKRMDGFAFAPGLGKRMDSFAFAPGLGKRGMDSLAFAGGLGKRMDSFAFAPGLGKRMDSFAFAPGLGKRGLDRYGFVGG : 320
Lottia.g       : -----------------------------------------------------------------QLGKRGLDQYGFAGQ : 210
Lymnaea.s      : -------------------------------------------------------------------------------- :   -
Pinctada.f     : -------------------------------------------------------------------------------- :   -
Biomphalaria.g : -----------------------------------------------------------------GIGKRPFDRYAFAGG : 190
Crassostrea.g  : -----------------------------------------------------------------TLGKRKLDQYSFMGN : 189


                          *       340         *       360         *       380         *       400
ObClone614     : ---GKRYDPVLFGGLGKR---------------MDPMLFGGLGKKMDPNMFG----TLGKRMDSMMFGH----------- : 208
OvClone195     : ---GKRYDPMLFGGLGKR---------------VDPM-FGGLGKKDGPPIC----------LEP---------------- : 124
Octopus.b      : ---GKRYDPVLFGGLGKR---------------MDPMLFGGLGKKMDPNMFG----TLGKRMDSMMFGH----------- : 208
Octopus.v      : ---GKRYDPMLFGGLGKR---------------VDPMLFGGLGKKMDPNMFG----TLGKRMDSMMFGH----------- : 208
Helix.a        : IEKRQLDTNGLAGGIGKRQA----------NIKIDIYGFAGGIGKRSEEDG-----FKYDDIDDVEAES----------- : 215
Aplysia.c      : LGKRGMDHFAFTGGLGKRDSGEASGDLEEGKRGLDAYSFTGALGKRGLDRYGFVGGLGKRGMDDFAFSPGLGKKRMDSFM : 400
Lottia.g       : LGKRGLDHYGFAGQLGKR--------------GLDQYGFAGQLGKRGFDQFGFAGQLGKRGLDHYGFAG----------- : 265
Lymnaea.s      : -------KFGFAGGIGKR--------------RLDRFNFAGGIGKRTPD-------DFEDNVVDFEKQA----------- : 173
Pinctada.f     : --VRNSAYYYRYGSRGRPIYT--------QTRGMDRFSFAARLGKRDAQNG----------------------------- : 124
Biomphalaria.g : IGKRPIDKFGFYGGIGKR--------------RLDRFSFAGGIGKRIPD-------LEEVSAAELSEAA----------- : 238
Crassostrea.g  : LGKRRLDSHRYFGSLGKR--------------ALDRYGFFGGLGKRADTLG-----NSQENIQGADKDE----------- : 239
                             G  G4r               6D   F g   K

                          *       420         *       440         *       460         *       480
ObClone614     : ---------------LGKRDVSGIKVGDFE-------------------------------------------------- : 223
OvClone195     : -------------------------------------------------------------------------------- :   -
Octopus.b      : ---------------LGKRDVSGIKVGDFE-------------------------------------------------- : 223
Octopus.v      : ---------------LGKRDVSGIKVGDFE-------------------------------------------------- : 223
Helix.a        : ----ENDRHVDKR--SAQLDKQVHVI------------------------------------------------------ : 235
Aplysia.c      : FGSRLGKRGMDRFSFSGHLGKRKMDQFSFGPGLGKR-GFDHYGFTGGIGKRGFDHYGFTGGIGKRQLDPMLFSGRLGKRS : 479
Lottia.g       : ---QLGKRGLDQLGFTGQLGKRQMDIFGYRGQLGKRQSIDKYSFLG-------------AGIGKR-----------SVKN : 318
Lymnaea.s      : ----DEVRELDKR--STDTAESHMVA------------------------------------------------------ : 193
Pinctada.f     : ---------------TLTFSRQVFFP------------------------------------------------------ : 135
Biomphalaria.g : ----NDVEKRSVP--SAKSEKETVKST----------------------------------------------------- : 259
Crassostrea.g  : ---KFEQKRLYPYWYYRQGGSPIYTQTR---------GIDRFSFAARLGRR----------------------------- : 278


                          *       500
ObClone614     : -------------------------- :   -
OvClone195     : -------------------------- :   -
Octopus.b      : -------------------------- :   -
Octopus.v      : -------------------------- :   -
Helix.a        : -------------------------- :   -
Aplysia.c      : SSEQEEEDVRQVEKRSTTEEQSSKSL : 505
Lottia.g       : TAGIKKDDA----------------- : 327
Lymnaea.s      : -------------------------- :   -
Pinctada.f     : -------------------------- :   -
Biomphalaria.g : -------------------------- :   -
Crassostrea.g  : -------------------------- :   -


```
